# PanKmer: *k*-mer based and reference-free pangenome analysis

**DOI:** 10.1101/2023.03.31.535143

**Authors:** Anthony J. Aylward, Semar Petrus, Allen Mamerto, Nolan T. Hartwick, Todd P. Michael

## Abstract

**Summary:** Pangenomes are replacing single reference genomes as the definitive representation of DNA sequence within a species or clade. Pangenome analysis predominantly leverages graph-based methods that require computationally intensive multiple genome alignments, do not scale to highly complex eukaryotic genomes, limit their scope to identifying structural variants (SVs), or incur bias by relying on a reference genome. Here, we present PanKmer, a toolkit designed for reference-free analysis of pangenome datasets consisting of dozens to thousands of individual genomes. PanKmer decomposes a set of input genomes into a table of observed *k*-mers and their presence-absence values in each genome. These are stored in an efficient *k*-mer index data format that encodes SNPs, INDELs, and SVs. It also includes functions for downstream analysis of the *k*-mer index, such as calculating sequence similarity statistics between individuals at whole-genome or local scales. For example, *k*-mers can be “anchored” in any individual genome to quantify sequence variability or conservation at a specific locus. This facilitates workflows with various biological applications, e.g. identifying cases of hybridization between plant species. PanKmer provides researchers with a valuable and convenient means to explore the full scope of genetic variation in a population, without reference bias.

**Availability and implementation:** PanKmer is implemented as a Python package with components written in Rust, released under a BSD license. The source code is available from the Python Package Index (PyPI) at https://pypi.org/project/pankmer/ as well as Gitlab at https://gitlab.com/salk-tm/pankmer. Full documentation is available at https://salk-tm.gitlab.io/pankmer/.

**Supplementary information:** Supplementary data are available online

## 1 Introduction

Pangenomes consolidate genomic sequence data from multiple individual organisms into a single data structure representing the genomes of a population, species, or clade. Pangenomics was first applied to microbes, but has grown to include a wide variety of eukaryotes ([1–3]). The advent of plant and animal pangenomes coincided with the reduced cost of NGS sequencing technologies. Crop plants were an early subject of pangenome studies, including soybean, brassica, and wheat species ([4–6]). These pangenomes focused on genic sequence only, and they were built by collecting short-read whole-genome sequencing datasets and mapping to a reference, allowing relatively simple if biased assembly of multiple individual genomes. Improvements in long-read sequencing technology and assembly methods have enabled the construction of dozens or hundreds of high-quality genomes for a single plant or animal species. This has facilitated a wave of pangenome studies based on collections of entire assembled genomes ([7–13]).

The transition from representation and analysis of single genomes to pangenomes presents significant challenges. Most studied populations include extensive structural variation (SV), which means only a fraction of genes or intergenic sequences are present in all individuals. This fraction is referred to as the “core” genome, while the remainder is variously called “dispensable”, “variable”, or “accessory” ([2, 14, 15]). Furthermore, the core genome cannot be defined by a simple linear coordinate system. A useful pangenomic dataset must identify the core genome and facilitate analysis of the variable regions.

Currently the dominant methodology is pangenome sequence graphs, which consist of nodes representing segments of genomic sequence connected by edges which allow any individual genome to be traced as a path through the graph ([7,16–19]). These graphs have replaced “iterative assembly” methods to advance our understanding of genomic diversity, especially of SV, and demonstrated utility for crop breeding ([7,8,11,20]). However, their construction is far from a solved problem and generally relies either on computationally expensive multiple genome alignment or on biased alignment to a single reference. Their application to highly complex eukaryotic genomes, such as those of plants, is limited ([21–23]). Such graphs may be limited to as few as a dozen input genomes ([12, 24]). Other methods gain efficiency by focusing on specific categories of variants ([17, 19]).

Several methods have adopted a format based on a De Bruijn graph (DBG) representing overlaps of *k*-mers (genomic substrings of length k) rather than the sequence graph ([25–28]). One such method is PanTools, which defines the pangenome as a comprehensive representation of multiple annotated genomes and provides functions enabling gene-level analysis, sequence alignment, and phylogenomics.

*k*-mer decomposition is an alternative to graph-based pangenomes. In this framework the space of genomic sequences across a population is represented by a set of *k*-mers (sequences of length k). Each individual is represented by a subset: all *k*-mers observed in a single genome. This view does not depend on any coordinate system and therefore sidesteps the difficulties of multiple genome alignment and graph construction. Kmer-db demonstrated the use of *k*-mer decomposition for efficient analysis of microbial genomes ([29]). *k*-mers have many applications in genomics and pangenomics, and they have recently been used as markers for GWAS ([21, 25, 26, 30–36]).

Here we present PanKmer, a non-graphical *k*-mer decomposition method designed to efficiently represent and analyze many forms of variation in large pangenomic datasets, with no reliance on a reference genome and no assumption of annotation.

## 2 Features & Implementation

### 2.1 *k*-mer index

The foundational component of PanKmer’s pangenome representation is the *k-mer index*. It is constructed from a set *G* of input genomes by decomposing them into a set *K* of all unique canonical *k*-mers and noting for each input genome which *k*-mers are present and which are absent (Fig. 1a). This is similar to the content of Kmer-db’s *k*-mer database ([29]). Each *k*-mer is considered equivalent to its reverse complement and is recorded in canonical form. The *k*-mer index *X* is then a |*K*| by|*G*| table of binary values indicating presence/absence of each canonical *k*-mer in each genome. The index can integrate an arbitrary number of genomes from one or several species, requires no reference genome, and enables a range of downstream analyses.

**Figure 1.**
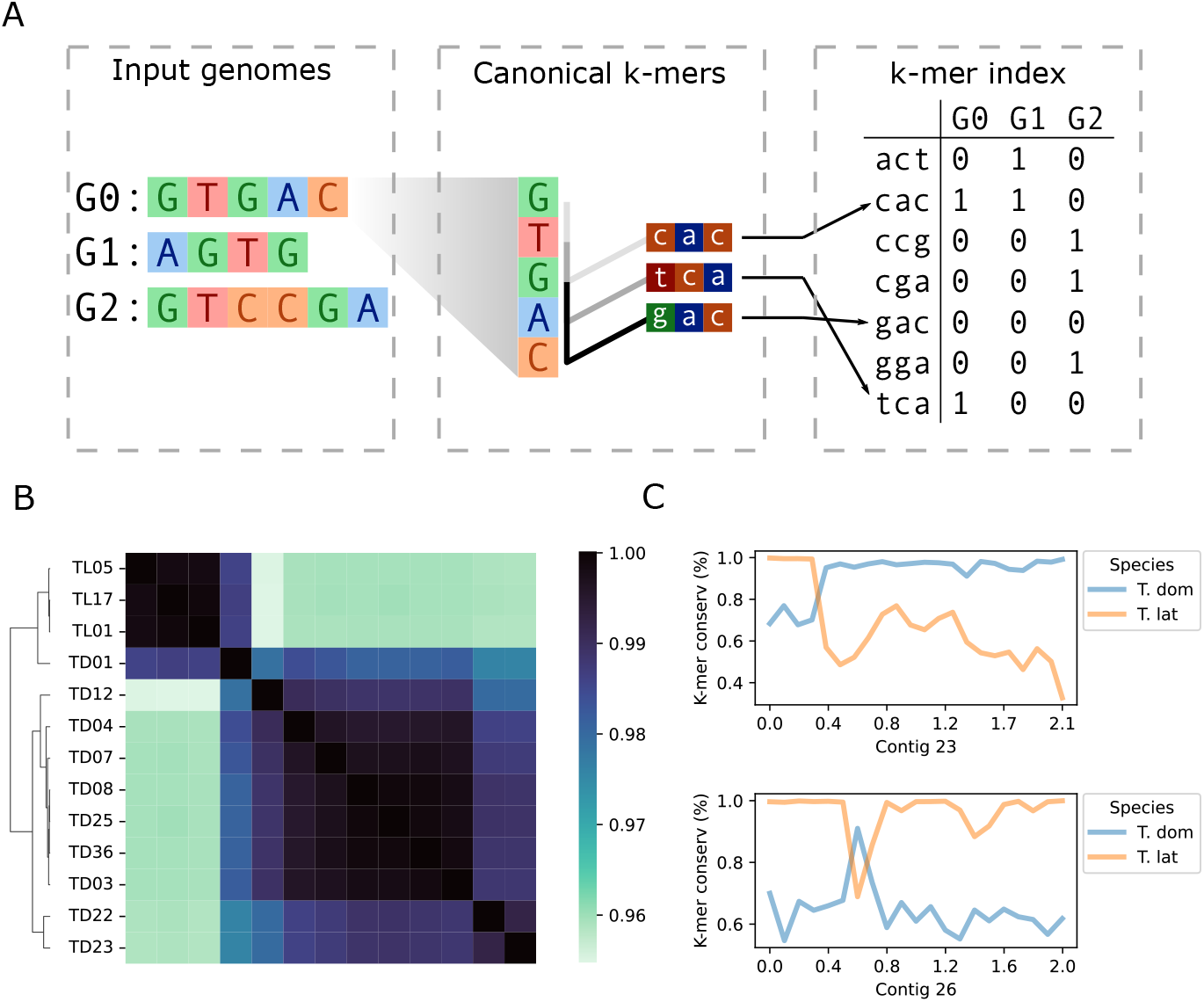
PanKmer enables the rapid estimation of relatedness across the pangenome as well as analysis of specific loci. A) Schematic of procedure for constructing the *k*-mer index. In the example, each genome G0-G2 is decomposed into canonical 3-mers. Each 3-mer is equivalent to its reverse complement, and the lexicographically first is the canonical form. Each 3-mer is assigned an integer value, and its presence/absence is recorded for each genome in the index. B) Relatedness heatmap of Typha pangenome, ANI values shown. C) Genome anchoring plots of representative contigs in TD01. Average *k*-mer conservation of 100-kb bins shown, where *k*-mer conservation is the fraction of TD or TL genomes that include each *k*-mer along the contig.

The index is constructed by scanning all |*G*| input genomes sequentially, recording newly encountered *k*-mers, and updating presence/absence values with each new genome scanned. To make efficient use of all available CPU’s, this process is parallelized across the theoretical *k*-mer space. The set *κ* of all possible canonical *k*-mers has size 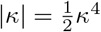, which is divided among *n* segments *κ*_1_…*κ*_*n*_. Index construction is then divided into *n* subprocesses, each of which constructs a sub-index *X*_*i*_ skipping *k*-mers not included in *κ*_*i*_. To efficiently store and access the *k*-mer index on disk, each *k*-mer is converted to an integer value.

The resultant index is robust to varying contiguity in the input genomes, so chromosome-level assemblies can be directly compared to unscaffolded contigs or unaligned reads. The implementation presented here uses *k* = 31, chosen for three reasons. First, 31-mers are short enough to be encoded as 64-bit integers ([37]). Second, they are long enough to impose a low rate of non-unique *k* –mers occurring by chance ([25]). Finally, 31-mers have been used successfully to define variation in previous studies ([34, 37]).

### 2.2 Adjacency matrix

Once the *k*-mer index is constructed, each input genome *g*_*i*_ is represented by |*K*| binary values representing presence or absence of each *k*-mer in *K*. This provides a natural means of calculating pairwise similarity/adjacency values for the input genomes. PanKmer includes a function to calculate the number of shared *k*-mers between all pairs of input genomes and return them as an adjacency matrix. Subsequently, the adjacency values can be used to perform a hierarchical clustering of input genomes and plot adjacency values as a heatmap. The adjacency values may also be converted to Jaccard, QV as described in MerQury ([38]), a symmetric version of QV (SQV), or Average Nucleotide Identity (ANI) (Fig. 1b).

### 2.3 Genome anchoring

While the *k*-mer index does not rely on any specified reference genome, it can be used to contextualize individual sequences. Given a sufficient *k*-mer length (e.g. the default *k* = 31), we can assume that each *k*-mer present in an individual genome *g*_*i*_ occurs approximately once in *g*_*i*_. Therefore, we can quantify variation across any locus in an “anchor” genome *g*_*i*_ by walking along the sequence, checking the *k*-mer that corresponds to each position, and calculating the fraction of genomes in *G* which share that *k*-mer. We refer to this fraction as the “*k*-mer conservation” value at each position. High *k*-mer conservation values indicate core loci which are conserved in many individuals across the pangenome, while low values indicate variable loci which are present only in *g*_*i*_ and a small number of other genomes. Hence, core sequences will have high *k*-mer conservation levels in all target genomes, while variable sequences will have relatively lower levels in each genome that features them.

## 3 Results

To demonstrate the utility of PanKmer, we constructed a cross-species pangenome of cattail downloaded from NCBI SRA: 10 *Typha domingensis* (TD) and 3 *Typha latifolia* (TL) genomes (Fig. 1b) (Table S1). Three of the TD genomes (TD01, TD22, TD23) were highly heterozygous (Figure S1) (Table 1). We used *k*-mer profiles to compute two measures of adjacency, the number of shared *k*- mers and Average Nucleotide Identity (ANI) (Fig. 1b). The 3 TL genomes were highly similar to one another, with an average Jaccard similarity value of 0.92 (186M shared *k*-mers) for TL-TL comparisons, while the average similarity of TD-TD comparisons was 0.60 (151M shared *k*-mers) (Table S2-3). The average similarity of TL-TD comparisons was 0.19 (64M shared *k*-mers). We also constructed single-species pangenomes of T. domingensis and T. latifolia (Figure S2) (Table S4-7). One TD genome, TD01, was an outlier relative to other TD, with Jaccard similarity of only 0.38 (127M shared *k*-mers) on average. Conversely, TD01 showed relatively high similarity to TL genomes, averaging 0.48 (148M shared *k*-mers). To inspect TD01 more closely, we calculated average *k*-mer conservation in 100kb bins across its assembled contigs and observed a mixture of TD and TL sequences. A close examination of variability anchored in TD01 revealed the presence of large and small introgressions (Fig. 1c, Figure S5). High heterozygosity together with the presence of introgressions suggest TD01 is the descendant of a recent TD-TL hybridization event.

**Table 1.** Basic statistics of *T. domingensis* and *T. latifolia* genome assemblies.

| Name | Predicted<br>genome<br>size (Mb) | Heterozygosity<br>(%) | Contig N50<br>(Mb) | GC content<br>(%) |
| --- | --- | --- | --- | --- |
| TD01 | 214 | 4.12 | 1.0 | 37.7 |
| TD03 | 222 | 0.13 | 11.5 | 37.6 |
| TD04 | 226 | 0.45 | 4.8 | 37.7 |
| TD07 | 263 | 0.14 | 7.7 | 37.6 |
| TD08 | 213 | 0.05 | 13.8 | 37.7 |
| TD12 | 224 | 0.22 | 6.1 | 37.5 |
| TD22 | 218 | 2.60 | 1.0 | 37.8 |
| TD23 | 213 | 2.60 | 0.7 | 37.8 |
| TD25 | 210 | 0.12 | 8.3 | 37.7 |
| TD36 | 215 | 0.15 | 8.9 | 37.6 |
| TL01 | 234 | 0.07 | 14.5 | 37.9 |
| TL05 | 218 | 0.07 | 14.5 | 37.8 |
| TL17 | 215 | 0.07 | 13.5 | 37.7 |

To explore suitability for larger eukaryote genomes and sample sizes, we benchmarked PanKmer on several published pangenomes and super-pangenomes, including *Solanum, Zea, H. Sapiens, A. thaliana* (supplementary material) (Table 2, Figure S7, Table S12) ([7, 39–41]). PanKmer successfully built *k*-mer indexes for all input pangenomes (Figure S8). We compared the performance of PanKmer to Kmer-db (supplementary material) (Figure S9, Table S13). We found that PanKmer had a smaller memory footprint than Kmer-db, due in part to its ability to divide *k*-mer decomposition into multiple rounds (supplementary material).

**Table 2.**
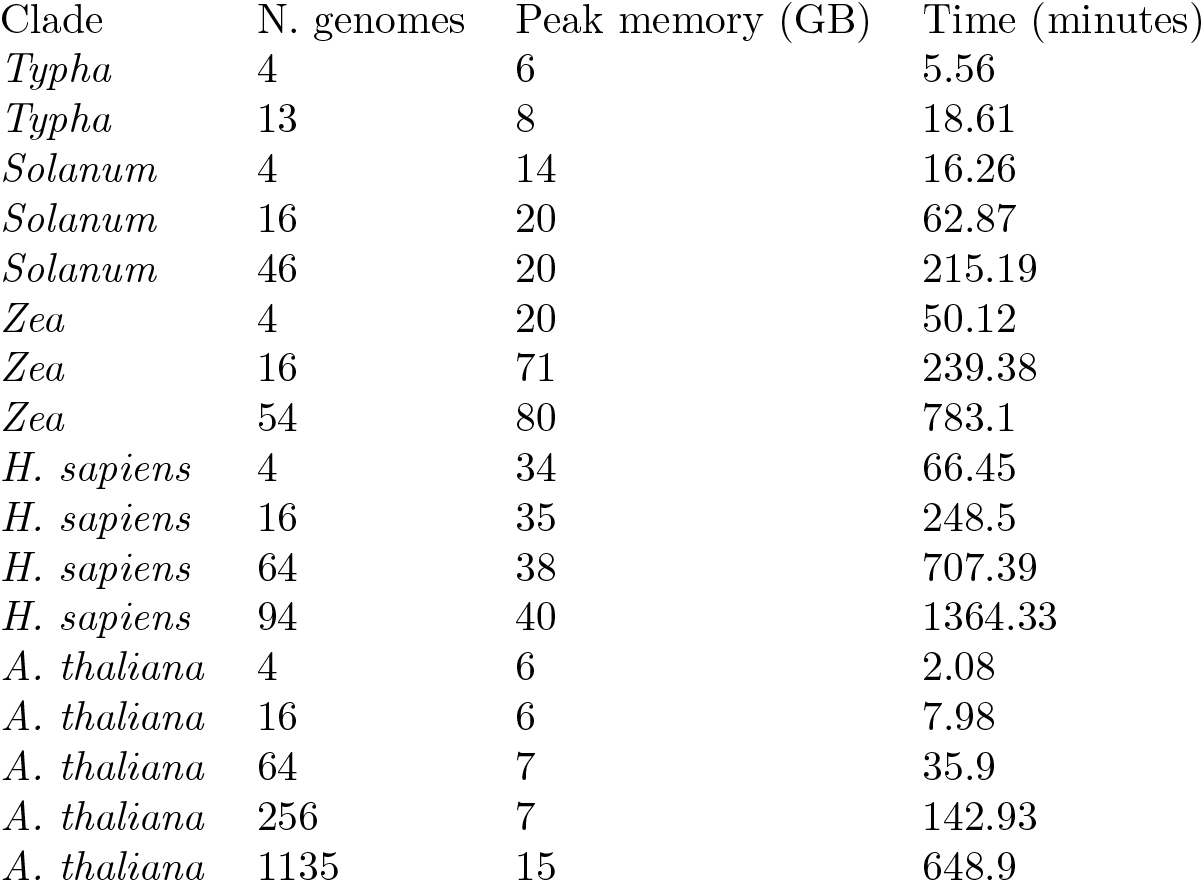
Benchmarking results of PanKmer configured for moderate memory usage and moderate runtime (16 rounds of *k*-mer decomposition)

| Clade | N. genomes | Peak memory (GB) | Time (minutes) |
| --- | --- | --- | --- |
| *Typha* | 4 | 6 | 5.56 |
| *Typha* | 13 | 8 | 18.61 |
| *Solanum* | 4 | 14 | 16.26 |
| *Solanum* | 16 | 20 | 62.87 |
| *Solanum* | 46 | 20 | 215.19 |
| *Zea* | 4 | 20 | 50.12 |
| *Zea* | 16 | 71 | 239.38 |
| *Zea* | 54 | 80 | 783.1 |
| *H. sapiens* | 4 | 34 | 66.45 |
| *H. sapiens* | 16 | 35 | 248.5 |
| *H. sapiens* | 64 | 38 | 707.39 |
| *H. sapiens* | 94 | 40 | 1364.33 |
| *A. thaliana* | 4 | 6 | 2.08 |
| *A. thaliana* | 16 | 6 | 7.98 |
| *A. thaliana* | 64 | 7 | 35.9 |
| *A. thaliana* | 256 | 7 | 142.93 |
| *A. thaliana* | 1135 | 15 | 648.9 |

## 4 Conclusion

PanKmer enables *k*-mer based analysis of pangenome datasets. It accepts as input a collection of genome assemblies or unaligned reads in FASTA format, and produces a *k*-mer index which is similar to the database of kmer-db ([29]). The index is agnostic to the contiguity of assemblies. A powerful feature of PanKmer is the ability to anchor the index in any individual genome and identify core or variable loci. This allows users to explore the full scope of genetic variation, without incurring bias from the choice of a single reference. PanKmer includes tools for downstream processing of the index which provide data visualizations and biological insights, such as identifying a hybridization event between *T*.*latifolia* and *T. domingensis*.

The primary advantage of PanKmer over other pangenome analysis tools is its ability to capture all forms of presence-absence variation, including SNPs, INDELs, SVs, and any variant that adds or removes a *k*-mer from the genome. Our reference-free and alignment-free algorithm is also more computationally tractable than graph-based methods. On the other hand, PanKmer is limited by inability to detect copy number variants in repetitive sequences (Figure S6), and by the loss of spatial relationships between *k*-mers in the index. However, their spatial context can be rescued by projecting the index onto an anchor genome. Currently PanKmer does not have genotyping functions, but these are planned for future releases.

## Supporting information

Software manual

## 5 Tutorials & documentation

Complete tutorials and documentation are available in the supplementary material and online: https://salk-tm.gitlab.io/pankmer/

## 6 Author contributions

A.A.: software (main developer, testing), writing (original draft, review and editing) funding acquisition. S.P.: software (original developer), writing (review and editing). A.M.: software (pipeline management, testing), writing (review and editing). N.H.: software (optimization) T.M.: Supervision, funding acquisition, writing (review and editing).

## 7 Acknowledgments

We thank Edward S Buckler, Bethany F Econopouly, Zachary R Miller, Peter J Bradbury, and Matt Wiese of the United States Dept. of Agriculture-Agricultural Research Service and Institute for Genomic Diversity at Cornell University for helpful discussion and collaboration on software features and implementation.

## 8 Funding

This research was funded by a Bill & Melinda Gates Foundation (BMGF) grant (INV-040541) to T.P.M., and the Tang Genomics Fund.

### 8.1 Conflict of interest

No conflict of interest is declared.

## 9 Data availability

The raw reads have been deposited in the NCBI short read archive (SRA) under BioProject PRJNA742003. Accession numbers for individual experiments are provided in the supplementary material. The genome assemblies are available for download from Michael lab AWS storage at: https://salk-tm-pub.s3.us-west-2.amazonaws.com/pub-supplementary/PRJNA742003-ASSEMBL.tar

---

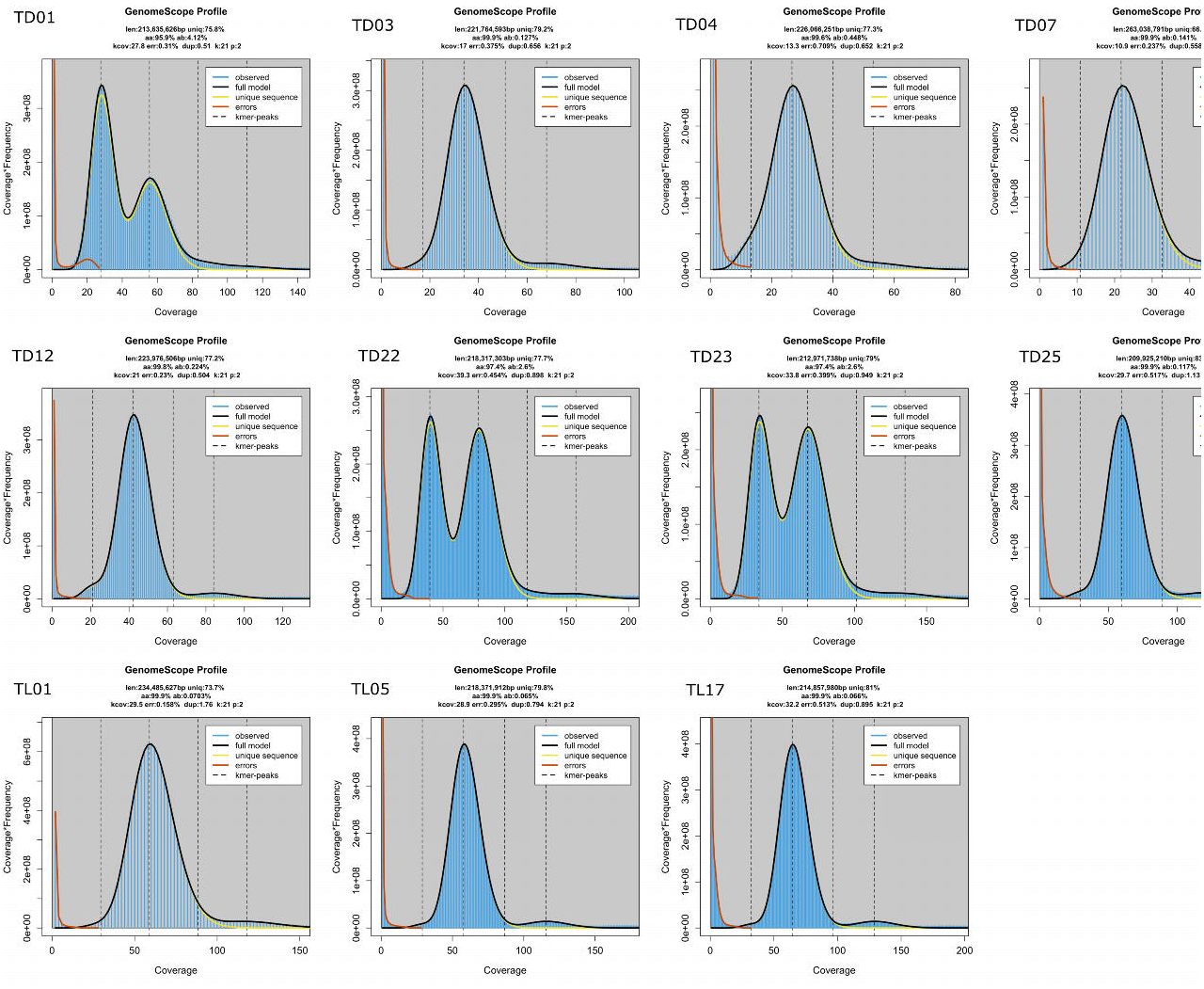

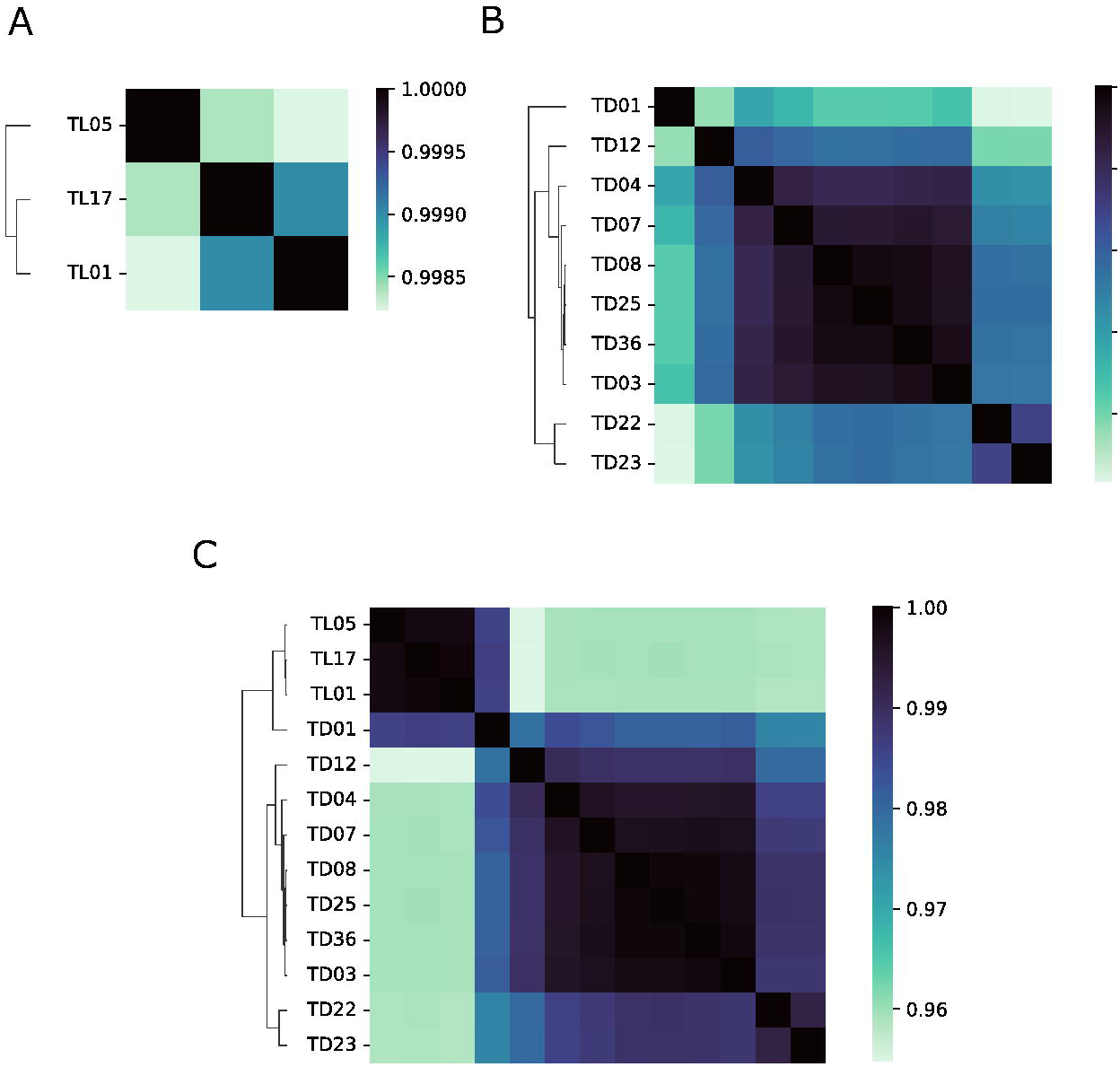

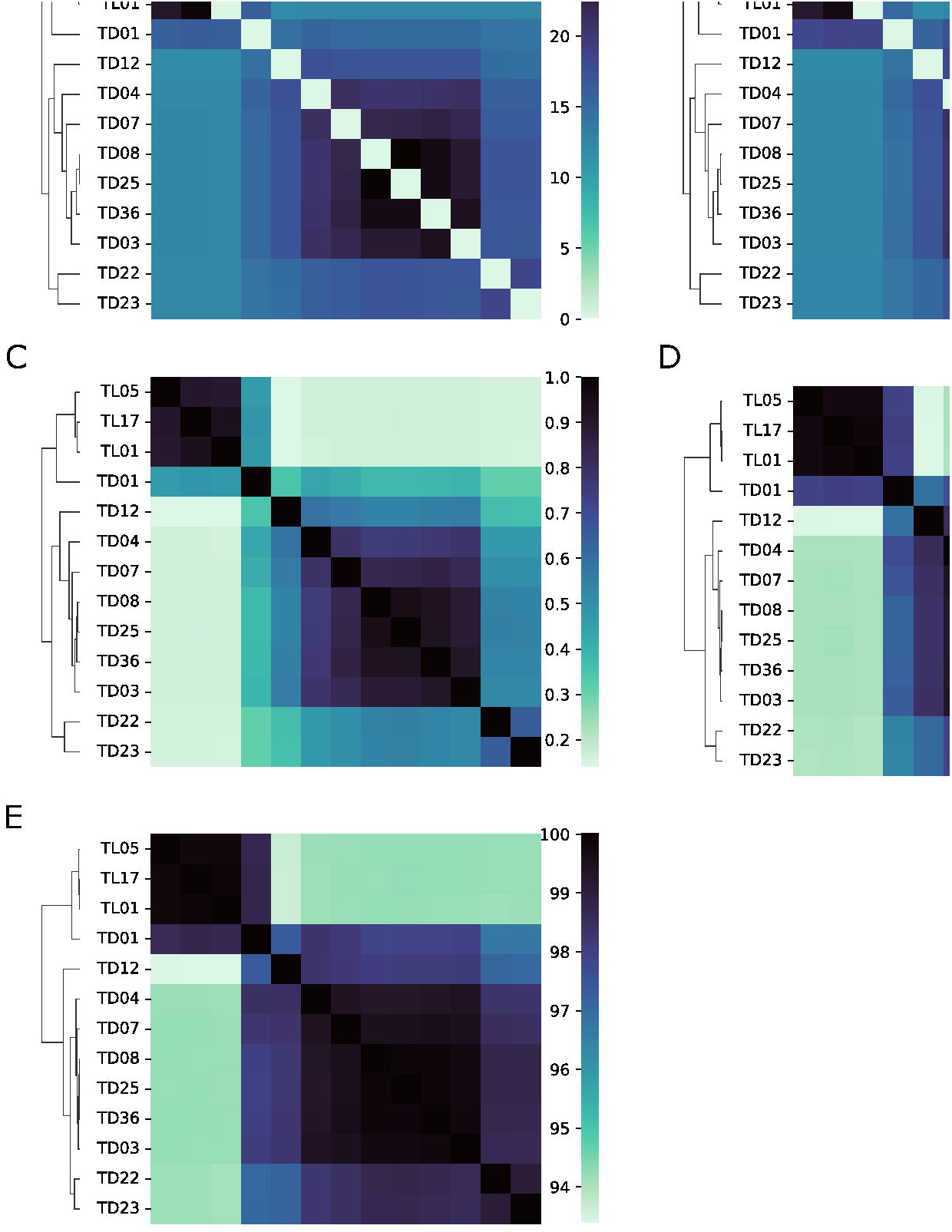

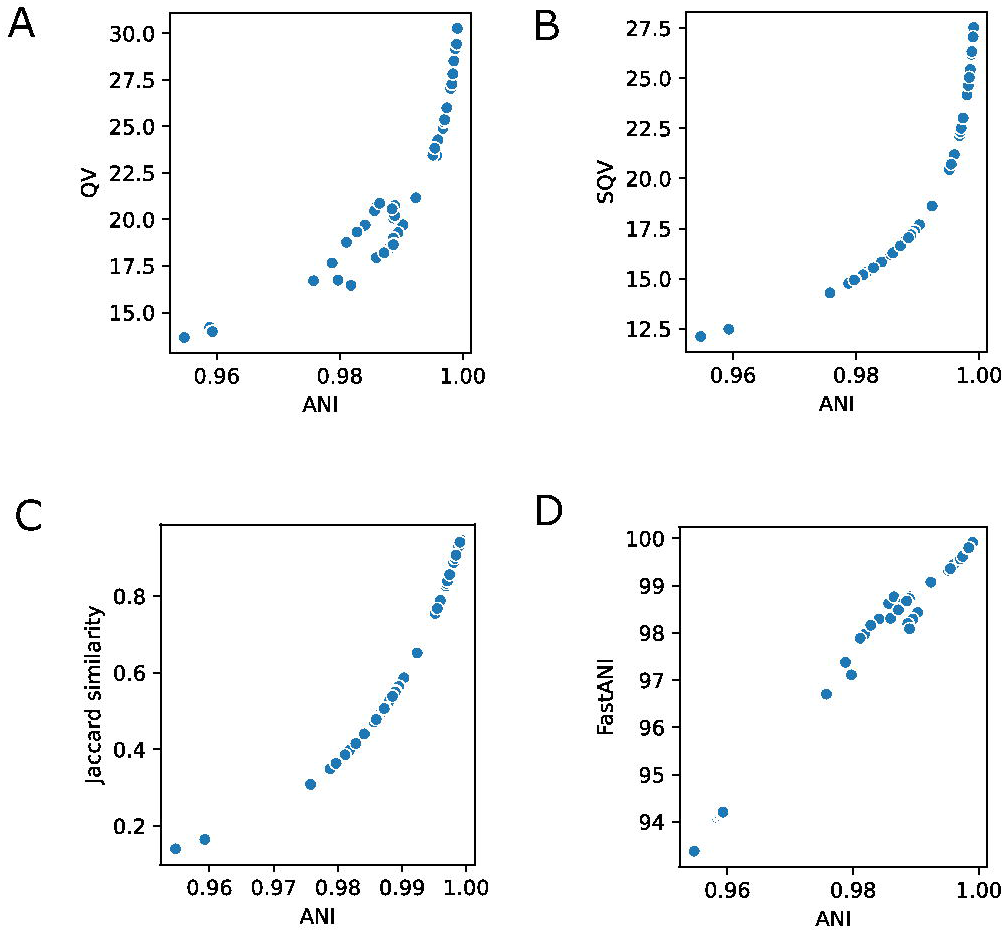

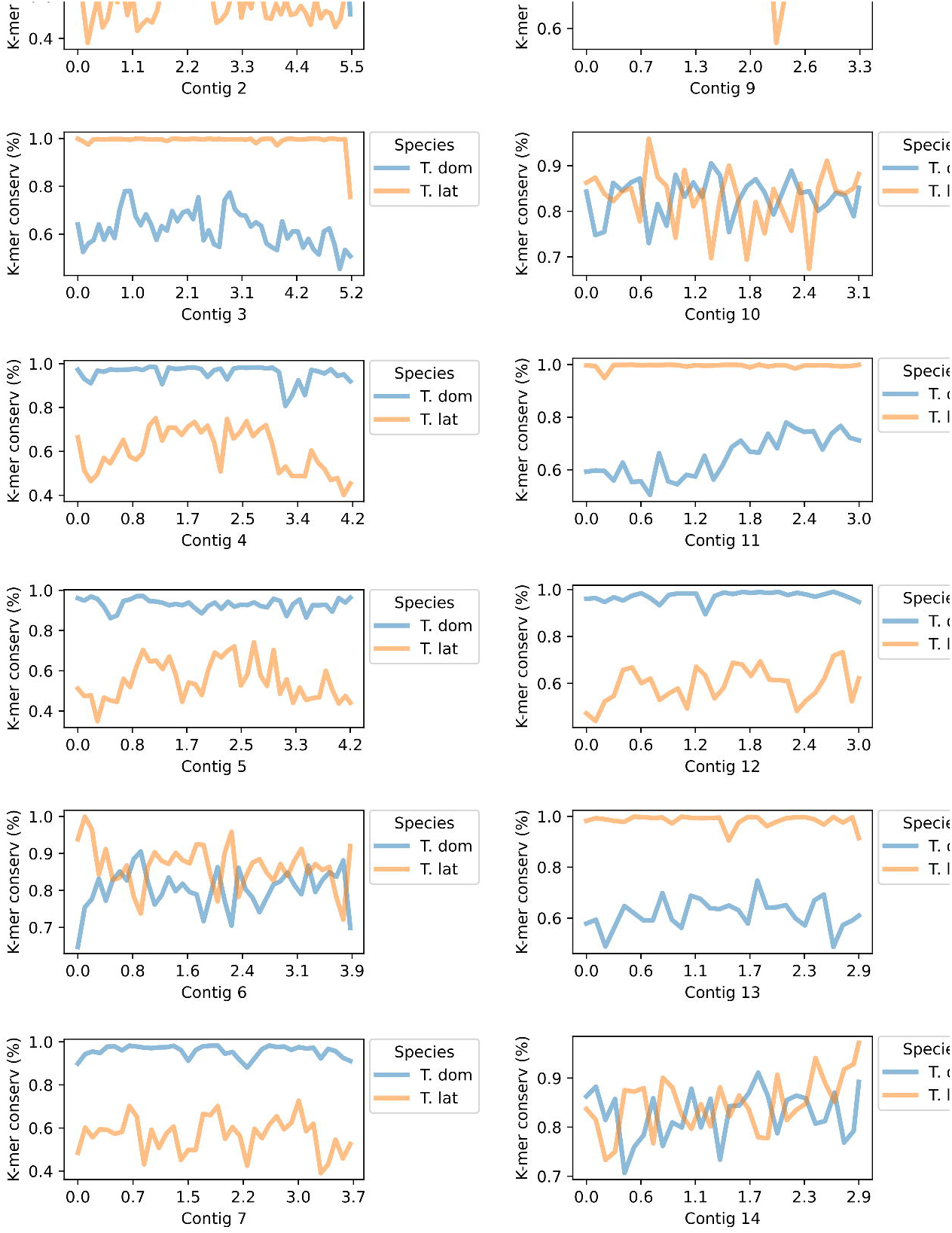

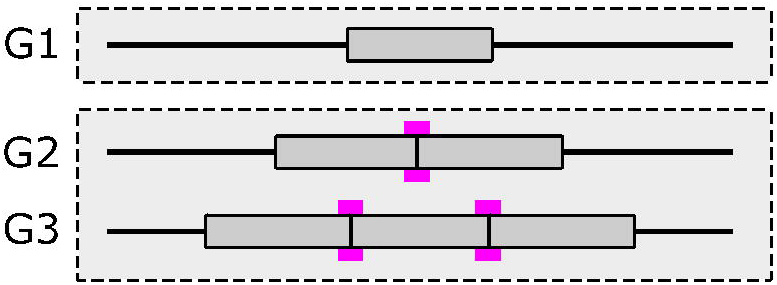

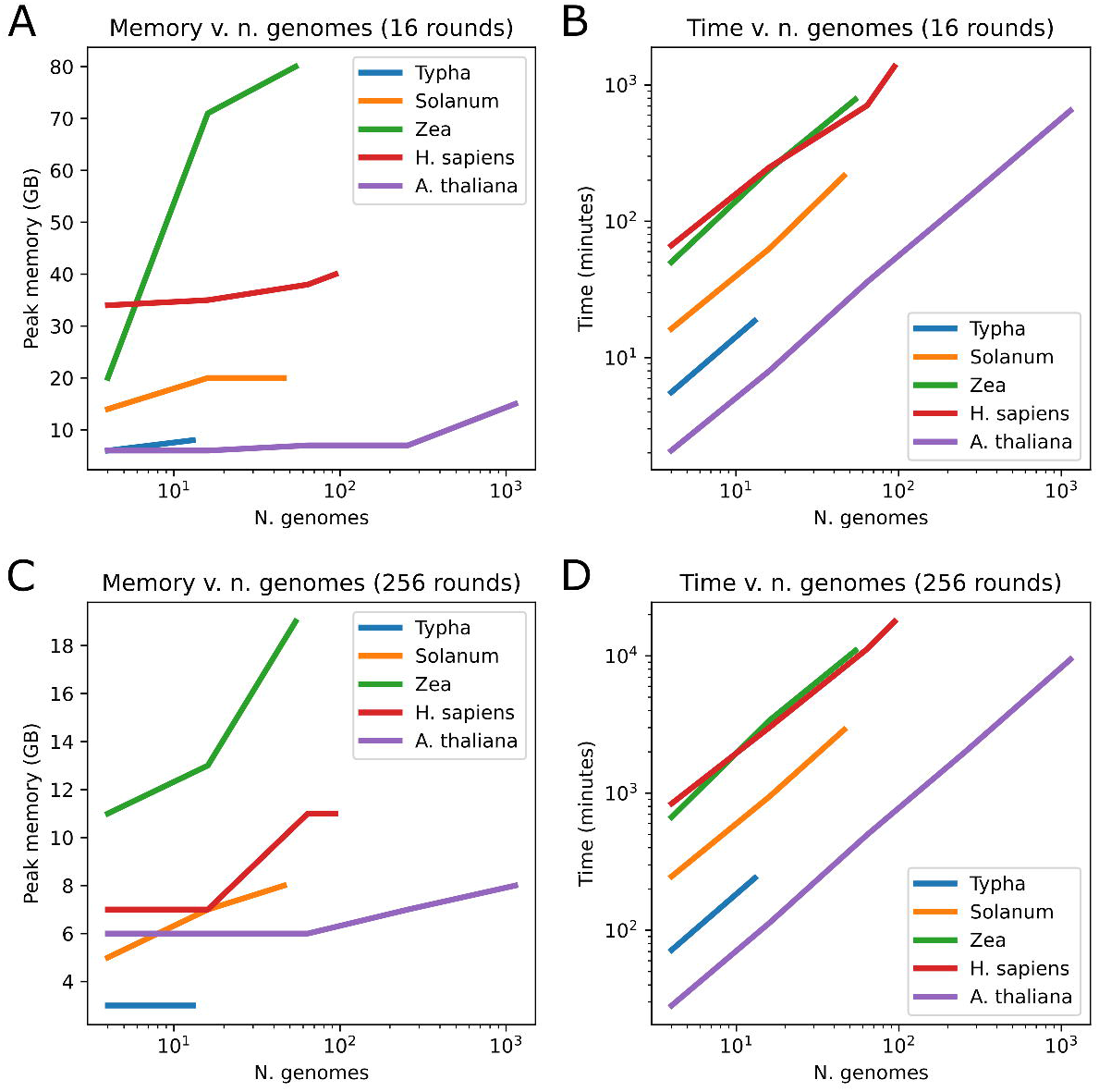

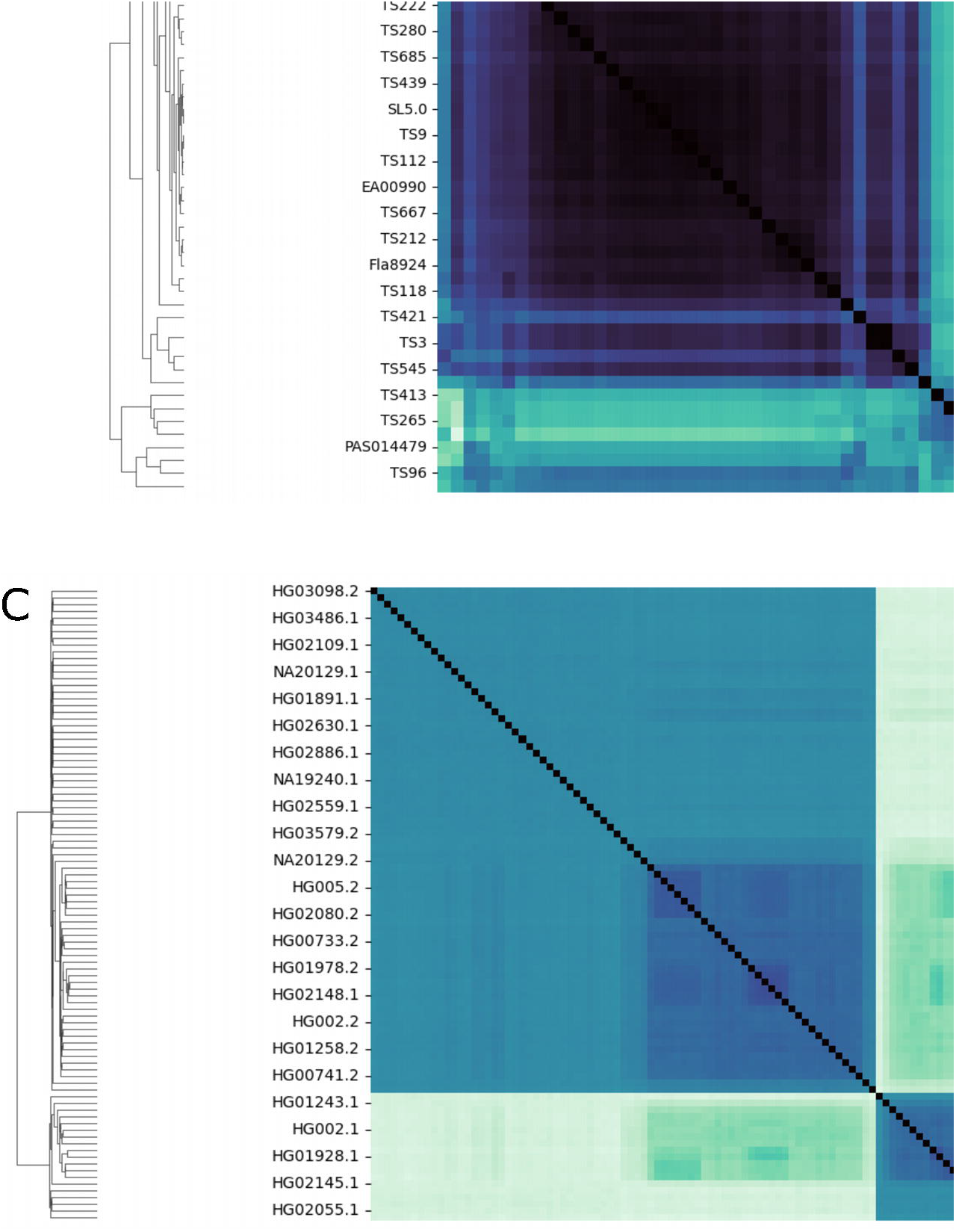

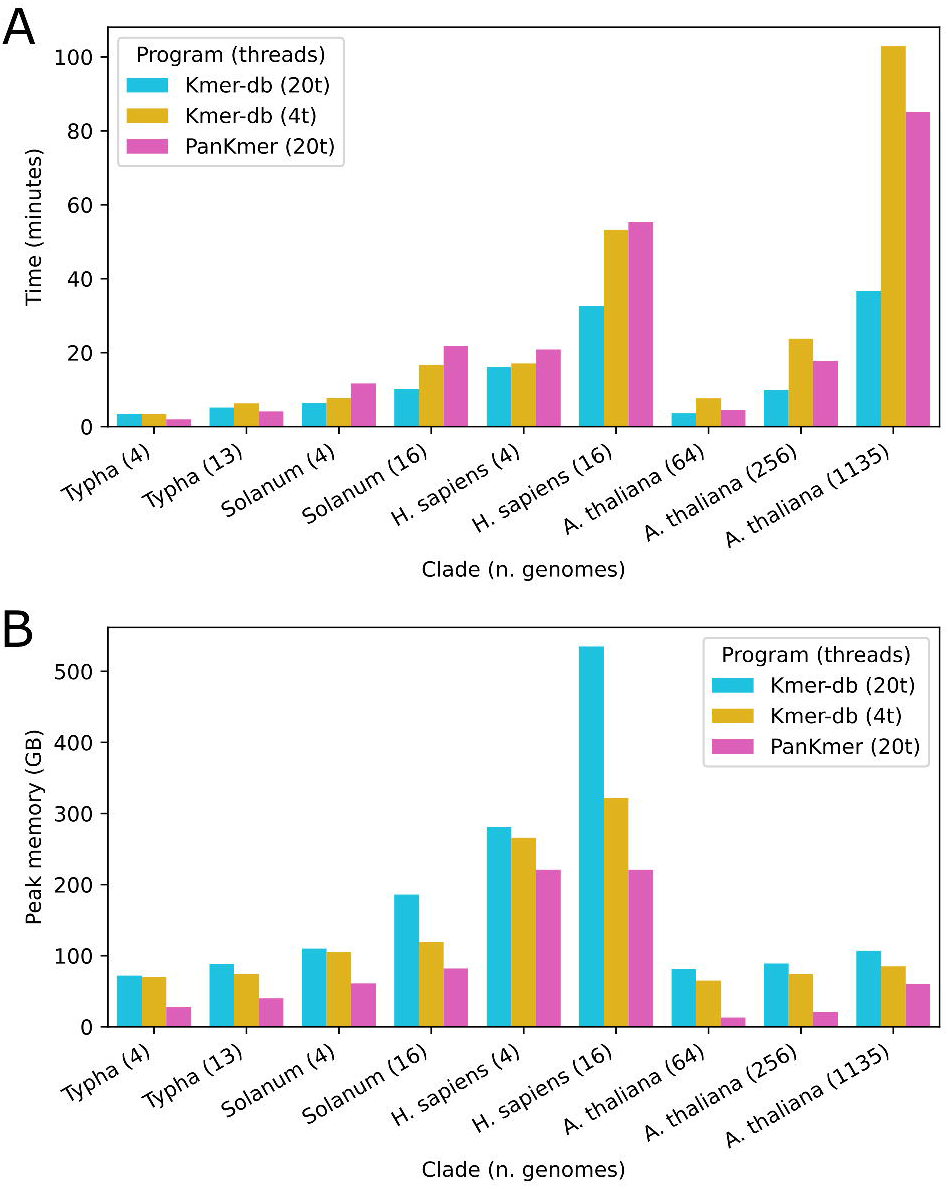

## Notes

https://pypi.org/project/pankmer/

https://gitlab.com/salk-tm/pankmer

https://salk-tm.gitlab.io/pankmer/

### Competing Interest Statement

The authors have declared no competing interest.

### Summary of Updates

Minor correction to abstract

## References

[1] D. Medini, C. Donati, H. Tettelin, V. Masignani, and R. Rappuoli. The microbial pan-genome. Curr. Opin. Genet. Dev., 15:589–594, 2005.

[2] A. A. Golicz, P. E. Bayer, P. L. Bhalla, J. Batley, and D. Edwards. Pangenomics comes of age: From bacteria to plant and animal applications. Trends Genet., 36:132–145, 2020.

[3] W. Li. et al. Plant pan-genomics: recent advances, new challenges, and roads ahead. J. Genet. Genomics, 49:833–846, 2022.

[4] Y. Li. et al. De novo assembly of soybean wild relatives for pan-genome analysis of diversity and agronomic traits. Nat. Biotechnol., 32:1045–1052, 2014.

[5] A. A. Golicz. et al. The pangenome of an agronomically important crop plant brassica oleracea. Nat. Commun., 7:13390, 2016.

[6] J. D. Montenegro. et al. The pangenome of hexaploid bread wheat. Plant J., 90:1007–1013, 2017.

[7] J. D. Montenegro. et al. Graph pangenome captures missing heritability and empowers tomato breeding. Nature, 606:527–534, 2022.

[8] L. Shang. et al. A super pan-genomic landscape of rice. Cell Res., 32:878–896, 2022.

[9] X. Tong. et al. High-resolution silkworm pan-genome provides genetic insights into artificial selection and ecological adaptation. Nat. Commun., 13:5619, 2022.

[10] X. Tong. et al. Improved pea reference genome and pan-genome highlight genomic features and evolutionary characteristics. Nat. Genet., 54:1553–1563, 2022.

[11] S. Gui. et al. A pan-zea genome map for enhancing maize improvement. Genome Biol., 23:178, 2022.

[12] H. Li. et al. Graph-based pan-genome reveals structural and sequence variations related to agronomic traits and domestication in cucumber. Nat. Commun., 13:682, 2022.

[13] H. Li. et al. Genome evolution and diversity of wild and cultivated potatoes. Nature, 606:535–541, 2022.

[14] H. Li. et al. Pangenomics in microbial and crop research: Progress, applications, and perspectives. Genes, 13:598, 2022.

[15] L. Lei. et al. Plant pan-genomics comes of age. Annu. Rev. Plant Biol., 72:411–435, 2021.

[16] J. A. Baaijens. et al. Computational graph pangenomics: a tutorial on data structures and their applications. Nat. Comput., 21:81–108, 2022.

[17] P. J. Bradbury. et al. The practical haplotype graph, a platform for storing and using pangenomes for imputation. Bioinformatics, 38:3698–3702, 2022.

[18] G. Hickey. et al. Genotyping structural variants in pangenome graphs using the vg toolkit. Genome Biol., 21:265, 2020.

[19] H. Li, X. Feng, and C. Chu. The design and construction of reference pangenome graphs with minigraph. Genome Biol., 21:265, 2020.

[20] P. Ruperao. et al. Sorghum pan-genome explores the functional utility for genomic-assisted breeding to accelerate the genetic gain. Front. Plant Sci., 12, 2021.

[21] A. W. Khan. et al. Super-pangenome by integrating the wild side of a species for accelerated crop improvement. Trends Plant Sci., 25:148–158, 2020.

[22] M. F. Danilevicz, C. G. Tay Fernandez, J. I. Marsh, P. E. Bayer, and D. Edwards. Plant pangenomics: approaches, applications and advancements. Curr. Opin. Plant Biol., 54:18–25, 2020.

[23] P. E. Bayer, A. A. Golicz, A. Scheben, J. Batley, and D. Edwards. Plant pan-genomes are the new reference. Nat. Plants, 6:914–920, 2020.

[24] X. Zhang. et al. Pan-genome of raphanus highlights genetic variation and introgression among domesticated, wild, and weedy radishes. Mol. Plant, 14:2032–2055, 2020.

[25] S. Sheikhizadeh, M. E. Schranz, M. Akdel, D. de Ridder, and S. Smit. Pantools: representation, storage and exploration of pan-genomic data. Bioinformatics, 32, 2016.

[26] E. M. Jonkheer. et al. Pantools v3: functional annotation, classification and phylogenomics. Bioinforma. Oxf. Engl., 38:4403–4405, 2022.

[27] G. Holley and P. Melsted. Bifrost: highly parallel construction and indexing of colored and compacted de bruijn graphs. Genome Biol., 21:249, 2020.

[28] F. Almodaresi, H. Sarkar, A. Srivastava, and R. Patro. A space and time-efficient index for the compacted colored de bruijn graph. Bioinformatics, 34, 2018.

[29] S. Deorowicz, A. Gudyśs, M. D/lugosz, M. Kokot, and A. Danek. Kmer-db: instant evolutionary distance estimation. Bioinformatics, 35:133–136, 2019.

[30] P. K. Gupta. Gwas for genetics of complex quantitative traits: Genome to pangenome and snps to svs and k-mers. BioEssays, 43, 2021.

[31] M. Jayakodi, M. Schreiber, N. Stein, and M. Mascher. Building pan-genome infrastructures for crop plants and their use in association genetics. DNA Res., 28, 2021.

[32] G. Holley, R. Wittler, and J. Stoye. Bloom filter trie: an alignment-free and reference-free data structure for pan-genome storage. Algorithms Mol. Biol., 11:3, 2016.

[33] E. Aun, A. Brauer, V. Kisand, T. Tenson, and M. Remm. A k-mer-based method for the identification of phenotype-associated genomic biomarkers and predicting phenotypes of sequenced bacteria. PLoS Comput. Biol., 14, 2018.

[34] Y. Voichek and D. Weigel. Identifying genetic variants underlying phenotypic variation in plants without complete genomes. Nat. Genet., 52:534–540, 2020.

[35] J. Ebler. et al. k-mer-based genome-wide association studies in plants: Advances, challenges, and perspectives. Nat. Genet., 54:518–525, 2022.

[36] B. Karikari, M.A. Lemay, and F. Belzile. Pangenome-based genome inference allows efficient and accurate genotyping across a wide spectrum of variant classes. Genes, 14:1439, 2023.

[37] A. Rahman, I. Hallgrśımsdśottir, M. Eisen, and L. Pachter. Association mapping from sequencing reads using k-mers. eLife, 7, 2018.

[38] A. Rhie, B. P. Walenz, S. Koren, and A. M. Phillippy. Merqury: reference-free quality, completeness, and phasing assessment for genome assemblies. Genome Biol., 21:245, 2020.

[39] M. R. Woodhouse. et al. A pan-genomic approach to genome databases using maize as a model system. BMC Plant Biol Biol., 21:385, 2021.

[40] W.-W. Liao. et al. A draft human pangenome reference. Nature, 617:312–324, 2023.

[41] C. Alonso-Blanco. et al 1,135 genomes reveal the global pattern of polymorphism in arabidopsis thaliana. Cell, 166:481–491, 2016.

