## Supplemental Information for "PanKmer: *k*-mer based and reference-free pangenome analysis"

### Supplementary material for article: PanKmer: $k$-mer based and reference-free pangenome analysis

August 29, 2023

**2 Supplementary figures** **7**

### 1 Supplementary methods

#### 1.1 *Typha* Genome assembly

Basecalled ONT fastq files passing QC (`fastq_pass`) were assembled using our previously described pipeline[1]. Briefly, fastq files were filtered by length for the longest 30x using a Illumina kmer-based genome size estimate. The 30x fastq files were overlapped using minimap2[2], the initial assembly was generated with miniasm[3], the resulting graph (gfa) was visually checked with Bandage[4], the assembly fasta was extracted from the gfa (`awk '/^S/{print ">"$2"\n"$3}' assembly_graph.gfa | fold > assembly_graph.fasta`), the consensus was generated with 3 iterative cycles of mapping the 30x reads back to the assembly with minimap2 followed by Racon[5], and the final assembly was polished iteratively 3 times with using 2x150 bp paired-end Illumina reads mapped using minimap2 (> 98% mapping) followed by pilon[6].

#### 1.2 Genome assembly statistics

Genome size and heterozygosity estimates of Typha domingensis and Typha latifolia genome assemblies were determined by genomescope[7] (Table 1, Figure S1). GC content was counted using SeqKit[8]. We also calculated completeness scores using MerQury[9] with $k = 21$. After consensus and polishing, all 13 Typha genomes had 100% completeness scores.

#### 1.3 Indexing algorithm

PanKmer indexing breaks down genomes to unique canonical $k$-mers ($k = 31$), and assigns each $k$-mer a bit vector indicating which samples include it. $k$-mers are represented as integers as in $k$-mer counters such as Jellyfish and Kmer-db[10, 11] For each $k$-mer in a contig, we encode the $k$-mer and its complement as unsigned 64-bit integers. The larger integer (lexicographically smaller $k$-mer) is selected as the canonical $k$-mer. Each base has a 2-bit integer substitute:

- A: 11
- C: 10
- G: 01
- T: 00

For example, the 5-mer ACGTC becomes 11-10-01-00-10 which equals 1938 in base 10. The reverse complement GACGT then becomes 01-11-10-01-00 which equals 1508 in base 10. Since 1938 is greater than 1508, it becomes the representative of the two, and is added to the index. Any $k$-mers containing a base that is not one of A, C, G, or T are skipped.

Each $k$-mer is added/updated in a hash table of $k$-mers wherein the keys are $k$-mers and the values are bit vectors called scores. A score indicates presence or absence of the $k$-mer in each sample. Scores are n bits where n is the number of samples. Each bit indicates presence/absence with sample 1 at the right most bit and sample n at the left most bit. For example, for a sample set of four genomes the score 0011 indicates presence in samples 3 and 4, while 1101 indicates presence in samples 1,2, and 4. After all $k$-mers in all contigs have been updated in the hash table, the $k$-mers and scores are written to their corresponding files on disk, along with a metadata file that completes the index.

#### 1.4 Choice of $k$

PanKmer's implementation represents each $k$-mer as an unsigned 64-bit integer, as in $k$-mer counting tools such as Jellyfish and Kmer-db[10, 11]. Hence the maximum value of $k$ is 32. Choosing $k \geq 31$ limits the theoretical rate of non-unique $k$-mers occurring by chance to fewer than 1 per 100 million $k$-mers[12]. This is important because some of PanKmer's downstream analysis functions, such as genome anchoring, work best when the non-unique k-mer rate is low. Hence, $k = 31$ and $k = 32$ are both suitable values for our application, and we chose $k = 31$. Furthermore, 31-bp k-mers have been applied successfully to define variation in previous studies[13, 14].

##### 1.5 Computation of QV, symmetric QV, and ANI

The QV of two sequences was defined by the authors of MerQury[9] as follows. Let K0, K1, be the sets of k-mers observed in each sequence. Then:

$$K_{shared} = |K_0 \cap K_1| \tag{1}$$
$$K_{total} = |K_0| \tag{2}$$
$$P = \left(\frac{K_{shared}}{K_{total}}\right)^{\frac{1}{k}} \tag{3}$$
$$E = 1 - P \tag{4}$$
$$QV(K_0, K_1) = -10\log_{10}(E) \tag{5}$$

Where $K_{shared}$ is the number of k-mers shared by $K_0$ and $K_1$, $K_{total}$ is the total number of $k$-mers in $K_0$. All QV values in Figures S3, S4, and Table S8 were computed using this formula, with $k = 31$. We define the symmetric QV (SQV) by the following:

$$K_{union} = |K_0 \cup K_1| \tag{6}$$
$$\hat{P} = \left(\frac{K_{shared}}{K_{union}}\right)^{\frac{1}{k}} \tag{7}$$
$$\hat{E} = 1 - \hat{P} \tag{8}$$
$$SQV(K_0, K_1) = -10\log_{10}(\hat{E}) \tag{9}$$

Where $K_{union}$ is the number of $k$-mers in the union of $K_0$ and $K_1$. We compute ANI values using the formula of FastANI[15]:

$$ANI(K_0, K_1) = 1 + \frac{1}{k}\ln\left(\frac{2J(K_0, K_1)}{1 + J(K_0, K_1)}\right) \tag{10}$$

Where $J(*, *)$ is the Jaccard similarity metric. All ANI values in Figures 1, S2, S3, S4, and Tables S3, S5, S7 were computed using this formula, with $k = 31$. Finally, we compared ANI values computed by PanKmer to values computed by FastANI.

#### 1.6 Copy number variation

The presence or absence of 31-mers can be used to detect most cases of SNP, INDEL, and SV including deletion, insertion, inversion, and translocation[13, 16]. However, k-mer presence/absence cannot tag CNVs unless their junctions produce unique K-mers. In practice, while some special cases can be tagged, a large class of CNVs are invisible to PanKmer (Figure S6).

#### 1.7 Configuration for large pangenomes

A challenge of building the PanKmer index (or Kmer-db database) from a large set of eukaryote genomes is the significant memory footprint of the operation. PanKmer includes the `--rounds` option to regulate peak memory usage. The default value of `--rounds` is 1, which means each input genome will be read once by each thread and the entire index will be built on the first pass. Increasing the value will divide the k-mer space into discrete blocks and construct a sub-index for each block, in sequence, finally merging all sub-indexes into the complete index. When the number of rounds is greater than 1, this scheme reduces peak memory usage by reducing the number of k-mers held in memory, at the expense of increased runtime since multiple k-mer decomposition passes must be made over each genome. Hence, when the pangenome size is small or large memory resources are available, fewer rounds are recommended, while more rounds are recommended for large pangenomes or environments with minimal memory.

#### 1.8 Benchmarks

##### 1.8.1 Benchmarking results

PanKmer was benchmarked on several pangenome datasets of various sizes(Figure S7, S8, Table S12, S13, S14):

1. The *Typha* super-pangenome dataset presented in this study (13 genomes total)
2. *Solanum* super-pangenome (46 genomes total)[17]
3. *Zea* super-pangenome (54 genomes total)[18]
4. *H. sapiens* pangenome (94 haplotypes total)[19]
5. *A. thaliana* pseudo-pangenome (1,135 pseudo-genomes total)[20]

Memory usage and runtime were also compared to Kmer-db (Figure S9)[11]. Shell scripts to execute all PanKmer-only benchmarking runs and all comparative PanKmer vs Kmer-db benchmarking runs can be found in the PanKmer GitLab repository at: https://gitlab.com/salk-tm/pankmer/-/tree/main/benchmark . All PanKmer runs had the `--threads` parameter set to execute with 20 threads. In all cases except the H. sapiens pangenome, the input genomes were compressed. The hard-coded $k$-mer size of 31 bp was in place for all cases. For

PanKmer only runs, the `--rounds` parameter was set either to 16 for moderate memory usage with moderate runtimes or to 256 for low memory usage with long runtimes. For PanKmer vs Kmer-db runs, `--rounds` was set to 1 for highest memory usage with shortest runtimes. All Kmer-db runs had parameters `-k 30` to use the maximum $k$-mer size of 30 and either `-t 20` or `-t 4` to use either 20 or 4 threads.

Building the PanKmer index or Kmer-db database from up to thousands of eukaryote genomes is computationally intensive. For each pangenome dataset, we executed 2-5 benchmarks using differently sized subsets of the input genomes, to demonstrate the resource requirements of PanKmer at different scales. The highest peak memory consumption occurred in the full-sized Zea super-pangenome benchmarks: 80 GB under the moderate memory/runtime parameter setting (`--rounds 16`) and 19 GB under the low memory / long runtime setting (`--rounds 256`). This is due to the large number of $k$-mers ($> 8 \times 10^9$). in the *Zea* pangenome dataset. The longest runtime occurred in the full-sized *H. sapiens* pangenome benchmarks: 1364 minutes under moderate memory/runtime (approx. 23 hours) and 17801 minutes under low memory / long runtime (approx. 12 days). This is due to the long length of the human genomes (> 3 gigabases). While 12 days is an exceptionally long time for a modern bioinformatics analysis, we note that this enabled building the PanKmer index of all 94 *H. sapiens* haplotypes using less than 12 GB of memory. In Kmer-db, the tradeoff of memory consumption against runtime is regulated by the number of threads. We chose two Kmer-db configurations, high memory/short runtime (`-t 20`) and low memory/long runtime (`-t 4`). We ran benchmarks comparing PanKmer to Kmer-db on *Typha*, *Solanum*, *H. sapiens* and *A. thaliana* datasets. In general, Kmer-db with `-t 20` was about twice as fast as PanKmer but had a much higher memory footprint, requiring 535 GB RAM to build a database from 16 *H. sapiens* genomes, while PanKmer required 221 GB to build the same pangenome at its most memory-intensive parameter setting. With `-t 4`, Kmer-db had similar runtimes to PanKmer but still had a higher memory footprint.

##### 1.8.2 Computing environment

The computer used for benchmarking was of the following configuration:

- 2 Intel Xeon Gold 6326 CPUs, 16 double-threaded cores per CPU, clocked at 2.9 GHz,
- 1024 GB RAM,
- 12 HDDs of size 2.4 TB each, `hdparm -t` reported buffered read speed 2317.27 MB/sec,
- Ubuntu 20.04 x86-64 operating system.

PanKmer was compiled with Rust version 1.71.0. Kmer-db binary was installed via Bioconda.

### References

[1] T.P. et al. Michael. High contiguity arabidopsis thaliana genome assembly with a single nanopore flow cell. *Nat. Commun.*, 9:541, 2016.

[2] H. Li. Minimap2: pairwise alignment for nucleotide sequences. *Bioinforma. Oxf. Engl.*, 34:3094–3100, 2018.

[3] H. Li. Minimap and miniasm: fast mapping and de novo assembly for noisy long sequences. *Bioinformatics*, 32:2103–2110, 2016.

[4] R. R. Wick, M. B. Schultz, J. Zobel, and K. E. Holt. Bandage: interactive visualization of de novo genome assemblies. *Bioinformatics*, 31:3350–3352, 2015.

[5] R. R. Wick, M. B. Schultz, J. Zobel, and K. E. Holt. Fast and accurate de novo genome assembly from long uncorrected reads. *Genome Res.*, 27:737–746, 2017.

[6] R. R. Wick, M. B. Schultz, J. Zobel, and K. E. Holt. An integrated tool for comprehensive microbial variant detection and genome assembly improvement. *PLOS ONE*, 9, 2014.

[7] T. R. Ranallo-Benavidez, K. S. Jaron, and M. C. Schatz. An integrated tool for comprehensive microbial variant detection and genome assembly improvement. *Nat. Commun.*, 11:1432, 2020.

[8] W. Shen, S. Le, Y. Li, and F. Hu. Seqkit: A cross-platform and ultrafast toolkit for fasta/q file manipulation. *PLOS ONE*, 11, 2016.

[9] A. Rhie, B. P. Walenz, S. Koren, and A. M. Phillippy. Merqury: reference-free quality, completeness, and phasing assessment for genome assemblies. *Genome Biol.*, 21:245, 2020.

[10] G. Marçais and C. Kingsford. A fast, lock-free approach for efficient parallel counting of occurrences of k-mers. *Bioinformatics*, 27:764–770, 2011.

[11] S. Deorowicz, A. Gudyś, M. Długosz, M. Kokot, and A. Danek. Kmer-db: instant evolutionary distance estimation. *Bioinformatics*, 35:133–136, 2019.

[12] S. Sheikhizadeh, M. E. Schranz, M. Akdel, D. de Ridder, and S. Smit. Pantools: representation, storage and exploration of pan-genomic data. *Bioinformatics*, 32, 2016.

[13] Y. Voichek and D. Weigel. Identifying genetic variants underlying phenotypic variation in plants without complete genomes. *Nat. Genet.*, 52:534–540, 2020.

[14] A. Rahman, I. Hallgrímsdóttir, M. Eisen, and L. Pachter. Association mapping from sequencing reads using k-mers. *eLife*, 7, 2018.

[15] C. Jain, L. M. Rodriguez-R, A. M. Phillippy, K. T. Konstantinidis, and S. Aluru. High throughput ani analysis of 90k prokaryotic genomes reveals clear species boundaries. *Nat. Commun.*, 9:5114, 2018.

[16] B. Karikari, M.A. Lemay, and F. Belzile. Pangenome-based genome inference allows efficient and accurate genotyping across a wide spectrum of variant classes. *Genes*, 14:1439, 2023.

[17] J. D. et al. Montenegro. Graph pangenome captures missing heritability and empowers tomato breeding. *Nature*, 606:527–534, 2022.

[18] M. R. et al. Woodhouse. A pan-genomic approach to genome databases using maize as a model system. *BMC Plant Biol Biol.*, 21:385, 2021.

[19] W.-W. et al. Liao. A draft human pangenome reference. *Nature*, 617:312–324, 2023.

[20] C. et al Alonso-Blanco. 1,135 genomes reveal the global pattern of polymorphism in arabidopsis thaliana. *Cell*, 166:481–491, 2016.

### 2 Supplementary figures

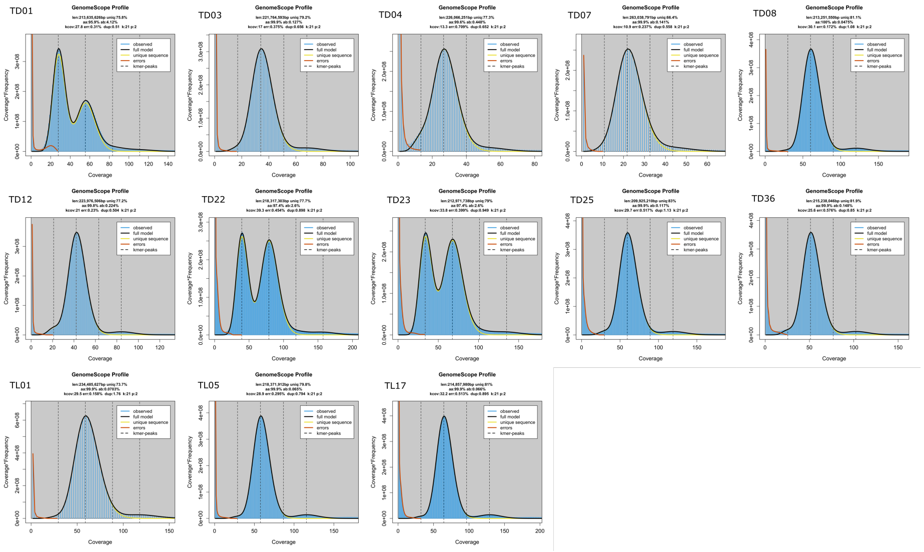

Figure S1: ***k*-mer spectra of Typha genomes.**

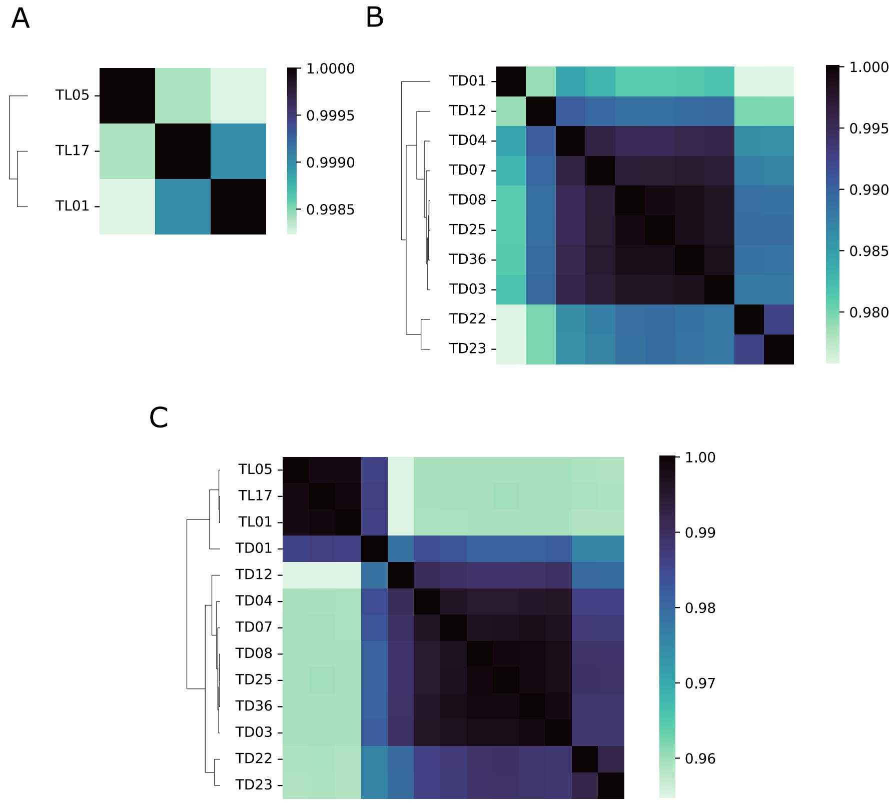

Figure S2: **Pairwise Average Nucleotide Identity across Typha pangenomes.** A) T. latifolia pangenome B) T. domingensis pangenome C) Typha super-pangenome

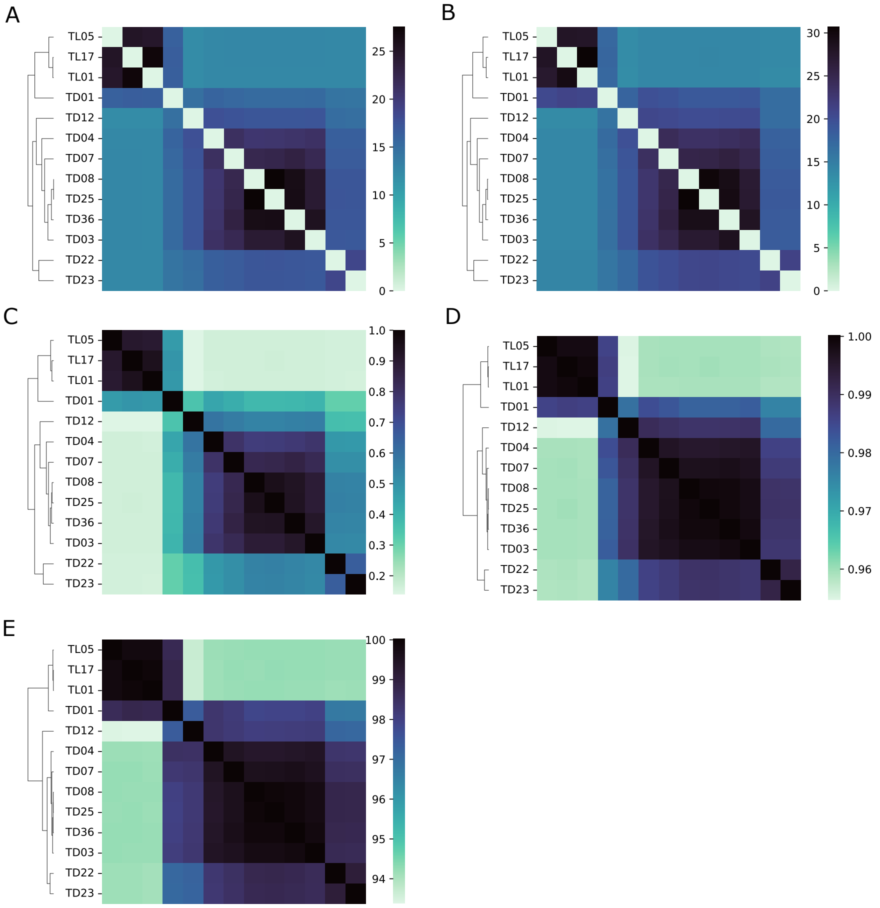

Figure S3: **Similarity metrics.** A) Symmetric QV B) MerQury QV C) Jaccard similarity D) ANI E) FastANI

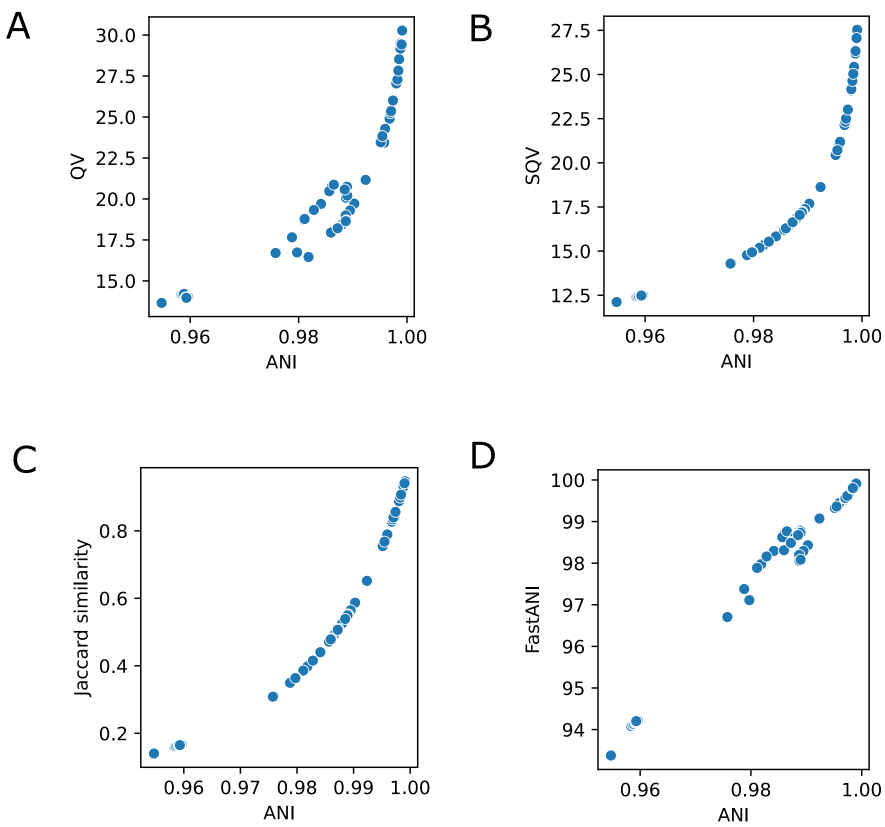

Figure S4: **Comparison of similarity metrics.** A) Average Nucleotide Identity vs. MerQury QV (SQV) B) ANI vs. Symmetric QV C) ANI vs. Jaccard similarity D) ANI vs. ANI computed by FastANI

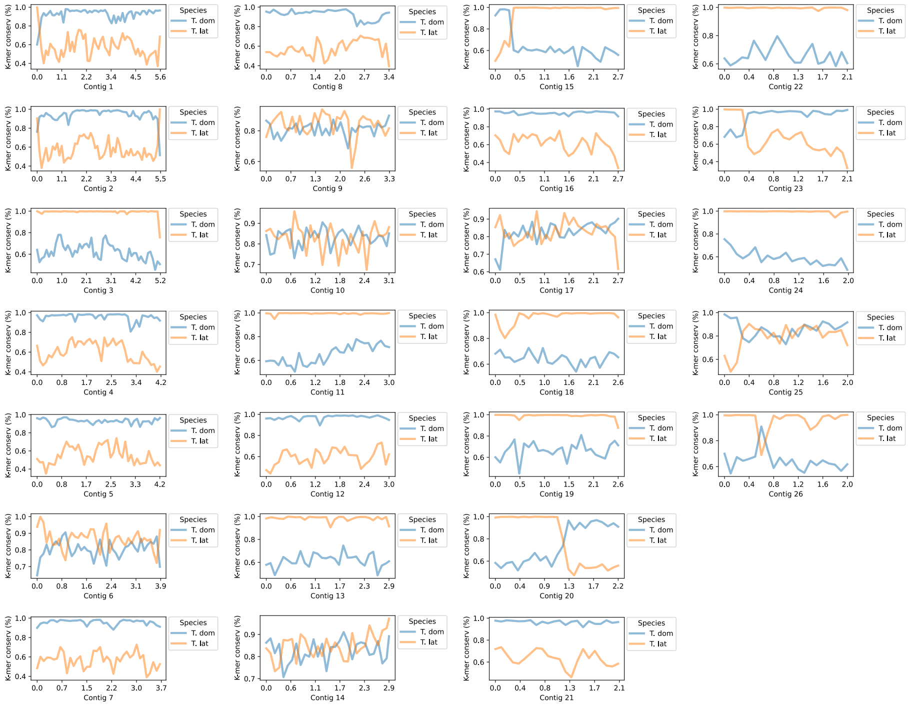

Figure S5: **k-mer conservation anchored in TD01 contigs.** All contigs larger than 2 Mb shown

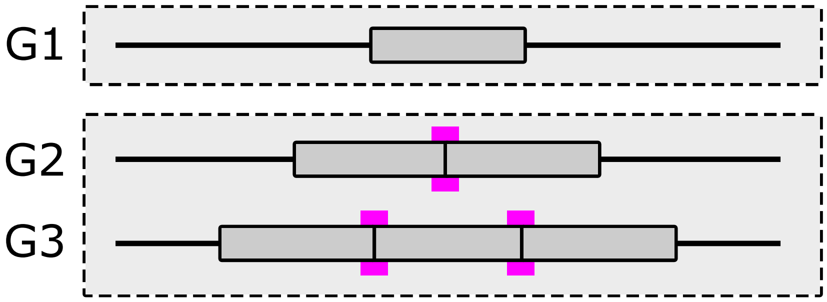

Figure S6: **Limitations of *k*-mers for tagging copy number variation.** Genomes G1, G2, and G3 have 1, 2, and 3 tandem copies of a repeat, respectively. K-mers produced by the copy junction highlighted in magenta can distinguish G2 and G3 from G1. However, G2 cannot be distinguished from G3 since the junction is present in both.

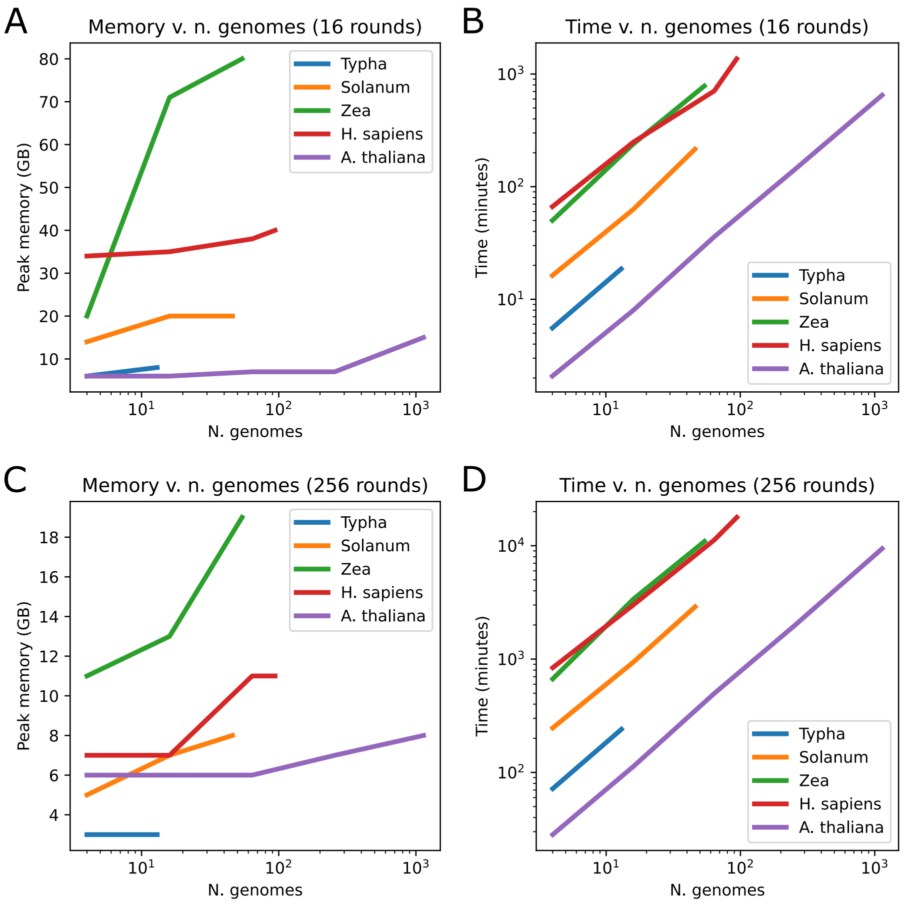

Figure S7: **Benchmarks.** Peak memory usage and runtime for pangenomes and super-pangenomes of various sizes, at either 16 rounds or 256 rounds of k-mer counting. A) Peak memory vs n. genomes with 16 rounds. B) Runtime vs n. Genomes with 16 rounds. C) Peak memory vs n. genomes with 256 rounds. D) Runtime vs n. genomes with 256 rounds.

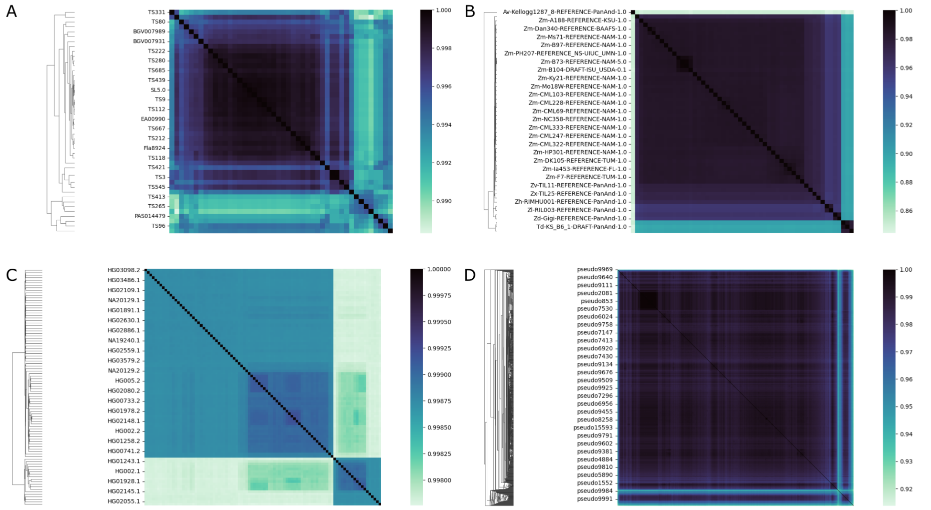

Figure S8: **Heatmaps of PanKmer benchmarking pangenome datasets.** Average Nucleotide Identity values shown. A) Solanum B) Zea C) H. sapiens D) A. thaliana

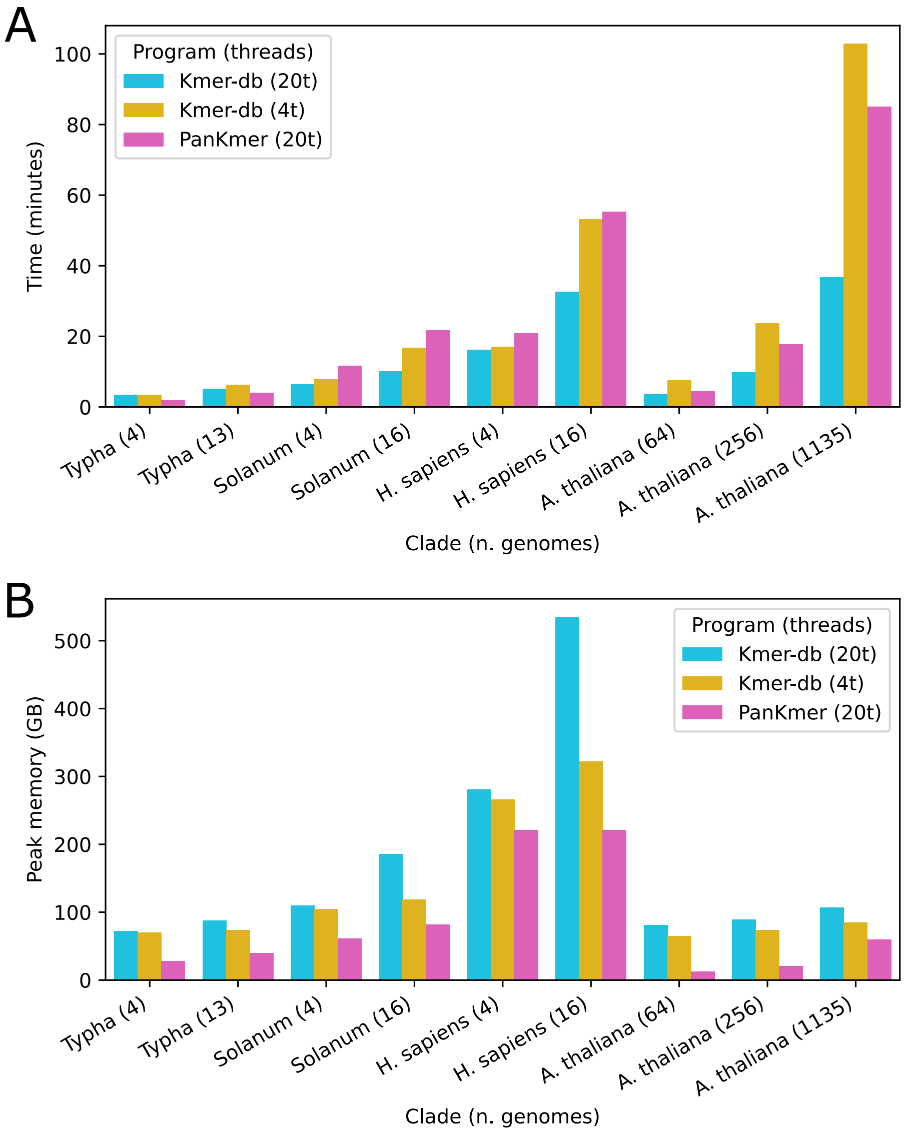

Figure S9: **PanKmer vs Kmer-db.** Comparison of runtime and memory usage for Kmer-db and PanKmer applied to several pangenome datasets. Kmer-db was run once with 20 and once with 4 threads for each dataset, PanKmer was run once with 20 threads. A) Runtime. B) Peak memory usage.
