## Supplementary material for "PanKmer: *k*-mer based and reference-free pangenome analysis": Software manual

### PanKmer

*Release 0.17.0*

**Michael lab**

**Aug 25, 2023**

**13 API reference** **57**

**Index** **63**

CHAPTER

ONE

### INSTALLATION

#### 1.1 In a conda environment

First create an environment that includes all dependencies:

```
conda create -c conda-forge -c bioconda -n pankmer cython \
  gff2bed more-itertools pybedtools python-newick pyfaidx \
  rust seaborn upsetplot urllib3
```

Then install PanKmer with `pip`:

```
conda activate pankmer
pip install pankmer
```

Alternatively, to install from source:

```
conda activate pankmer
git clone https://gitlab.com/salk-tm/pankmer.git
cd pankmer
pip install .
```

#### 1.2 With pip

PanKmer is built with Rust , so you will need to install it if you have not already done so. Then you can install PanKmer with `pip`:

```
pip install pankmer
```

#### 1.3 Check installation

Check that the installation was successful by running:

```
pankmer --version
```

See the available subcommands by running `pankmer --help`:

```
usage: pankmer [-h] [--version]
               {index,count,collect,upset,subset,adj-matrix,tree,clustermap,similarity,
↪distance,anchor-region,anchor-genome,anchor-plot,anchor-heatmap,anchormap,dryrun,
↪download-example}
               ...

positional arguments:
  {index,count,collect,upset,subset,adj-matrix,tree,clustermap,similarity,distance,
↪anchor-region,anchor-genome,anchor-plot,anchor-heatmap,anchormap,dryrun,download-
↪example}
    index               generate k-mer index
    count               count k-mers in one or more indexes
    collect             calculate k-mer collection curve
    upset               generate upset plot
    subset              subset an index
    adj-matrix          generate adjacency matrix
    tree                generate a heirarchical clustering tree from an adjacency matrix
    clustermap          plot a clustered heatmap from the adjacency matrix
    similarity          generate a similarity matrix from an adjacency matrix
    distance            generate a distance matrix from an adjacency matrix
    anchor-region       anchor k-mers in a region
    anchor-genome       anchor k-mers in a genome
    anchor-plot         generate a plot from genome anchoring results
    anchor-heatmap      draw anchor heatmap
    anchormap           export an anchormap for downstream visualization
    dryrun              perform a dry run of indexing and print metadata
    download-example    download an example dataset
```

CHAPTER

TWO

### DEPENDENCIES

Pending shipment of binaries, installing PanKmer requires the Rust programming language and its crate manager Cargo.

PanKmer also depends on the following Python packages: biopython seaborn urllib3 newick pyfaidx gff2bed upsetplot pybedtools cython more-itertools

### CHAPTER THREE

### TUTORIAL

#### 3.1 Download example dataset

The `download-example` subcommand will download a small example dataset of Chr19 sequences from *S. polyrhiza*[1]
.

```
pankmer download-example -d .
```

After running this command the directory `PanKmer_example_Sp_Chr19/` will be present in the working directory. It contains FASTA files representing Chr19 from three genomes, and GFF files giving their gene annotations.

```
ls PanKmer_example_Sp_Chr19/*
```

```
PanKmer_example_Sp_Chr19/README.md

PanKmer_example_Sp_Chr19/Sp_Chr19_features:
Sp9509_oxford_v3_Chr19.gff3.gz Sp9512_a02_genes_Chr19.gff3.gz

PanKmer_example_Sp_Chr19/Sp_Chr19_genomes:
Sp7498_HiC_Chr19.fasta.gz Sp9509_oxford_v3_Chr19.fasta.gz Sp9512_a02_genome_Chr19.fasta.
↪gz
```

**Note:** GFF files are included in the example data, but are not required for basic pangenome analysis and are not used in this tutorial. Examples using GFF data can be found in the *examples* section.

To get started, navigate to the downloaded directory.

```
cd PanKmer_example_Sp_Chr19/
```

[1] Harkess A. et al. Improved Spirodela polyrhiza genome and proteomic analyses reveal a conserved chromosomal structure with high abundance of chloroplastic proteins favoring energy production. *Journal of Experimental Botany* 2021

#### 3.2 Build a *k*-mer index

The *k*-mer index is a table tracking presence or absence of *k*-mers in the set of input genomes. To build an index, use the `index` subcommand and provide a directory containing the input genomes.

```
pankmer index -g Sp_Chr19_genomes/ -o Sp_Chr19_index.tar
```

After completion, the index will be present as a tar file `Sp_Chr19_index.tar`.

```
tar -tvf Sp_Chr19_index.tar
```

```
Sp_Chr19_index/
Sp_Chr19_index/kmers.b.gz
Sp_Chr19_index/metadata.json
Sp_Chr19_index/scores.b.gz
```

**Note:** The input genomes argument provided with the `-g` flag can be a directory, a tar archive, or a space-separated list of FASTA files.

If the output argument provided with the `-o` flag ends with `.tar`, then the index will be written as a tar archive. Otherwise it will be written as a directory.

#### 3.3 Create an adjacency matrix

A useful application of the *k*-mer index is to generate an adjacency matrix. This is a table of *k*-mer similarity values for each pair of genomes in the index. We can generate one using the `adj_matrix` subcommand, which will produce a CSV or TSV file containing the matrix.

```
pankmer adj-matrix -i Sp_Chr19_index.tar -o Sp_Chr19_adj_matrix.csv
pankmer adj-matrix -i Sp_Chr19_index.tar -o Sp_Chr19_adj_matrix.tsv
```

**Note:** The input index argument provided with the `-i` flag can be tar archive or a directory.

#### 3.4 Plot a clustered heatmap

To visualize the adjacency matrix, we can plot a clustered heatmap of the adjacency values. In this case we use the Jaccard similarity metric for pairwise comparisons between genomes:

```
pankmer clustermap -i Sp_Chr19_adj_matrix.csv \
  -o Sp_Chr19_adj_matrix.svg \
  --metric jaccard \
  --width 6.5 \
  --height 6.5
```

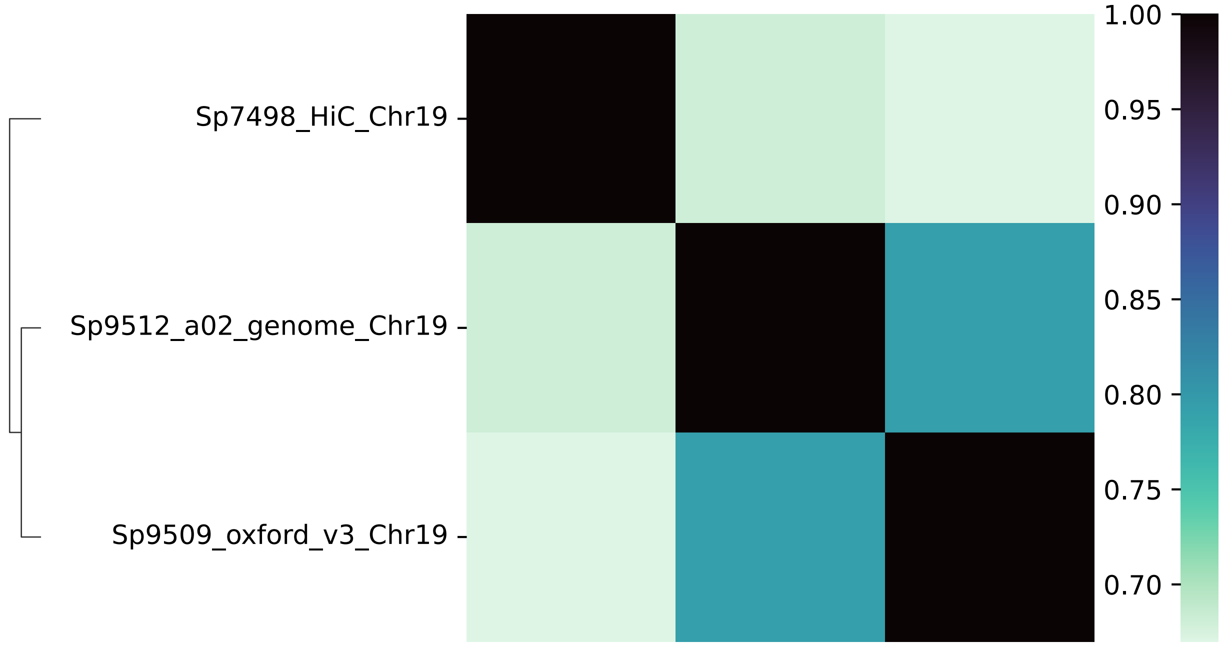

Fig. 1: Clustered heatmap generated from *S. polyrhiza* adjacency matrix.

CHAPTER

FOUR

### EXAMPLES

#### 4.1 Generate a Newick tree

Generate a hierarchical clustering of genomes in Newick tree format (e.g. for input to Cactus alignment).

First navigate to the example data directory `PanKmer_example_Sp_Chr19/` as shown in the *tutorial*.

Then, use `pankmer tree` with the `--newick` option to generate the tree. In this example we use the Jaccard metric for pairwise comparison of genomes.

```
pankmer tree --newick --metric jaccard -i Sp_Chr19_adj_matrix.csv
```

```
(Sp7498_HiC_Chr19:0.33006755822027156,(Sp9512_a02_genome_Chr19:0.20699749712873816,
↪Sp9509_oxford_v3_Chr19:0.20699749712873816)3:0.1230700610915334);
```

#### 4.2 Count total and diagnostic K-mers

Simply use `pankmer count`:

```
pankmer count -i Sp_Chr19_index.tar
```

```
index	total	diagnostic
Sp_Chr19_index.tar	4764864	836962
```

#### 4.3 Calculate collection curves

Use `pankmer collect` to calculate collection curves and assess pangenome completeness:

```
pankmer collect -i Sp_Chr19_index.tar -o Sp_Chr19_collect.svg
```

At the cost of some extra computation, 95% confidence intervals can be estimated using the `--conf` option:

```
pankmer collect --conf -i Sp_Chr19_index.tar -o Sp_Chr19_collect_conf.svg
```

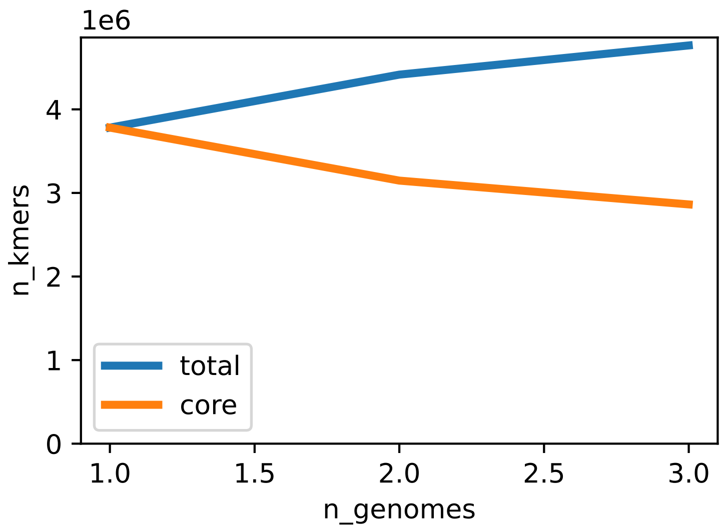

Fig. 1: *S. polyrhiza* pangenome collection curves.

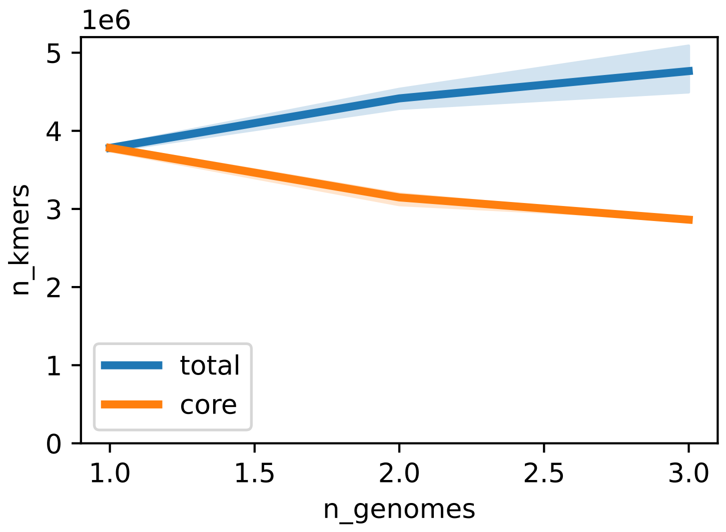

Fig. 2: *S. polyrhiza* pangenome collection curves with confidence intervals.

#### 4.4 Draw an UpSet plot

Use `pankmer upset` to draw an UpSet plot:

```
pankmer upset -i Sp_Chr19_index.tar -o Sp_Chr19_upset.svg -g Sp7498_HiC_Chr19 \
  Sp9509_oxford_v3_Chr19 Sp9512_a02_genome_Chr19
```

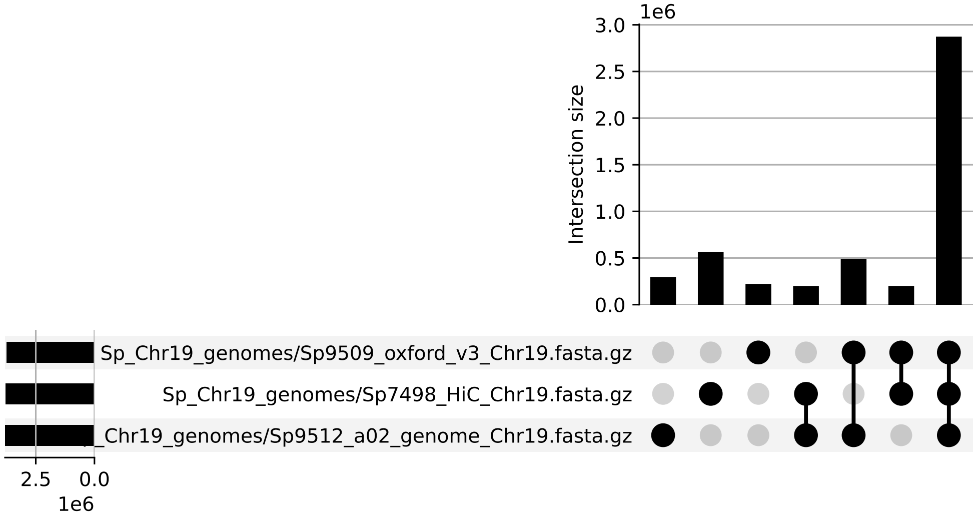

Fig. 3: *S. polyrhiza* UpSet plot.

**Note:** The output image of `pankmer upset` is generated by the upsetplot python library .

#### 4.5 Extract a subset pangenome

An index for a subset of genomes can be generated using `pankmer subset`:

```
pankmer subset -i Sp_Chr19_index.tar -o Sp7_Chr19_index -g Sp7498_HiC_Chr19
pankmer subset -i Sp_Chr19_index.tar -o Sp9_Chr19_index.tar \
  -g Sp9509_oxford_v3_Chr19 Sp9512_a02_genome_Chr19
```

The `-g` flag defines the genome subset.

#### 4.6 Genome anchoring: regional

Anchor the pangenome in a specific region of an anchor genome. This provides a view of variability across genomes in the index.

```
mkdir Sp_Chr19_anchor/
pankmer anchor-region -i Sp_Chr19_index.tar -r Sp_Chr19_genomes/Sp9509_oxford_v3_Chr19.fasta.gz \
  -c Chr19:384629-385934 > Sp_Chr19_anchor/Sp19g00080.bdg
```

We can visualize the resulting bedGraph file by loading it into a genome browser. Alternatively, we can apply gtracks .

```
pip install gtracks
gtracks-gff3-to-bed12 Sp_Chr19_features/Sp9509_oxford_v3_Chr19.gff3.gz | bgzip -c > Sp_Chr19_anchor/Sp9509_oxford_v3_Chr19.bed.gz
cd Sp_Chr19_anchor/
gtracks --genes Sp9509_oxford_v3_Chr19.bed.gz Chr19:384629-386134 Sp19g00080.bdg Sp19g00080.svg
```

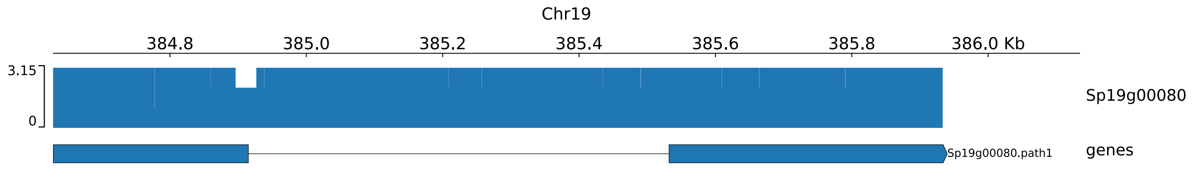

Fig. 4: K-mer coverage across the Sp19g00080 gene in the Sp9509 reference.

**Note:** `anchor-region` expects 1-based coordinates. To calculate coverage across an entire chromosome or contig, use coordinates of the format `<contig>:1-<contig-size>`

**Note:** For converting GFF3 to BED format, see also the UCSC genome browser utility GFF3ToGenePred . Also on bioconda

#### 4.7 Genome anchoring: regional from multiple indexes

We can also use `pankmer anchor-region` on multiple indexes. For example, using the subset indices generated above:

```
pankmer anchor-region -i Sp7_Chr19_index Sp9_Chr19_index.tar \
  -r Sp_Chr19_genomes/Sp9509_oxford_v3_Chr19.fasta.gz \
  -c Chr19:384629-385934 > Sp_Chr19_anchor/Sp19g00080_multi.bdg
```

This will produce a bedGraph file with two coverage columns.

```
head Sp_Chr19_anchor/Sp19g00080_multi.bdg
```

```
Chr19       384628  384629  1       1
Chr19       384629  384630  1       1
Chr19       384630  384631  1       1
Chr19       384631  384632  1       1
Chr19       384632  384633  1       1
Chr19       384633  384634  1       1
Chr19       384634  384635  1       1
Chr19       384635  384636  1       1
Chr19       384636  384637  1       1
Chr19       384637  384638  1       1
```

We can visualize these data with `gtracks`, but need to separate them into two bedGraph files.

```
cd Sp_Chr19_anchor/
cut -f 1-4 Sp19g00080_multi.bdg > Sp19g00080_Sp7.bdg
cut -f 1-3,5 Sp19g00080_multi.bdg > Sp19g00080_Sp9.bdg
gtracks --genes Sp9509_oxford_v3_Chr19.bed.gz --plot-type line:2 \
--overlay Chr19:384629-386134 Sp19g00080_Sp7.bdg Sp19g00080_Sp9.bdg \
Sp19g00080_multi.svg
```

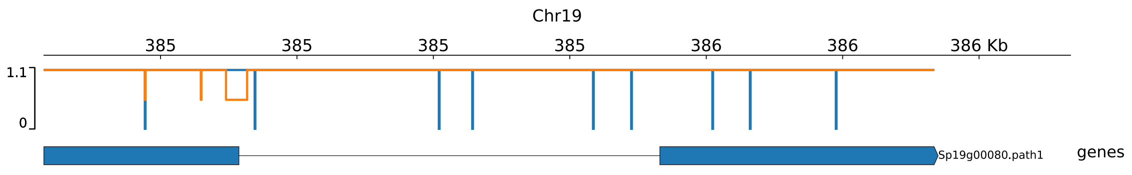

Fig. 5: K-mer coverage across the Sp19g00080 gene in the Sp9509 reference, from two indexes overlaid.

#### 4.8 Genome anchoring: genomic

To plot coverage across an entire chromosomes or contigs, use the `anchor-genome` subcommand.

```
mkdir Sp_Chr19_anchor_genome/
pankmer anchor-genome -i Sp_Chr19_index.tar \
  -r Sp_Chr19_genomes/Sp9509_oxford_v3_Chr19.fasta.gz \
  -c Chr19 \
  -t Sp_Chr19_anchor_genome/Sp9509_Chr19_coverage.tsv \
  -o Sp_Chr19_anchor_genome/Sp9509_Chr19_coverage.svg
```

Anchoring across the entire chromosome is computationally intensive so even the small example dataset will take a few minutes. The result is a plot of average coverage levels in 1-megabase bins across the chromosome.

The `anchor-genome` subcommand can also be used with multiple indexes.

```
pankmer anchor-genome --groups "Sp 7" "Sp 9" --legend \
  --x-label "Chromosome 19" \
  -i Sp7_Chr19_index Sp9_Chr19_index.tar \
  -r Sp_Chr19_genomes/Sp9509_oxford_v3_Chr19.fasta.gz \
  -c Chr19 \
```

(continues on next page)

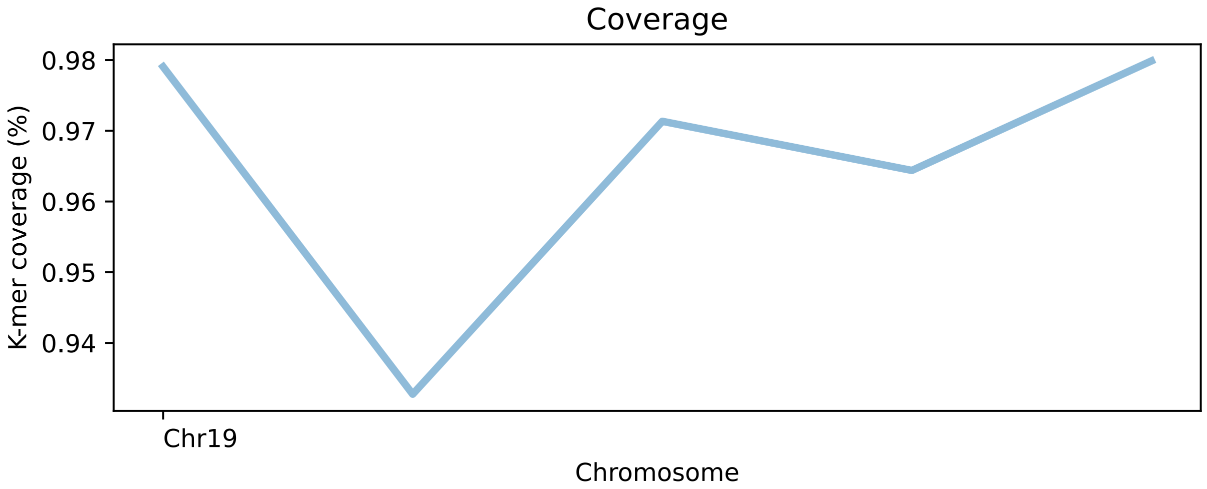

Fig. 6: K-mer anchoring across chromosome 19 in the Sp9509 reference.

(continued from previous page)

```
  -t Sp_Chr19_anchor-genome/Sp9509_Chr19_anchor_multi.tsv \
  -o Sp_Chr19_anchor-genome/Sp9509_Chr19_anchor_multi.svg
```

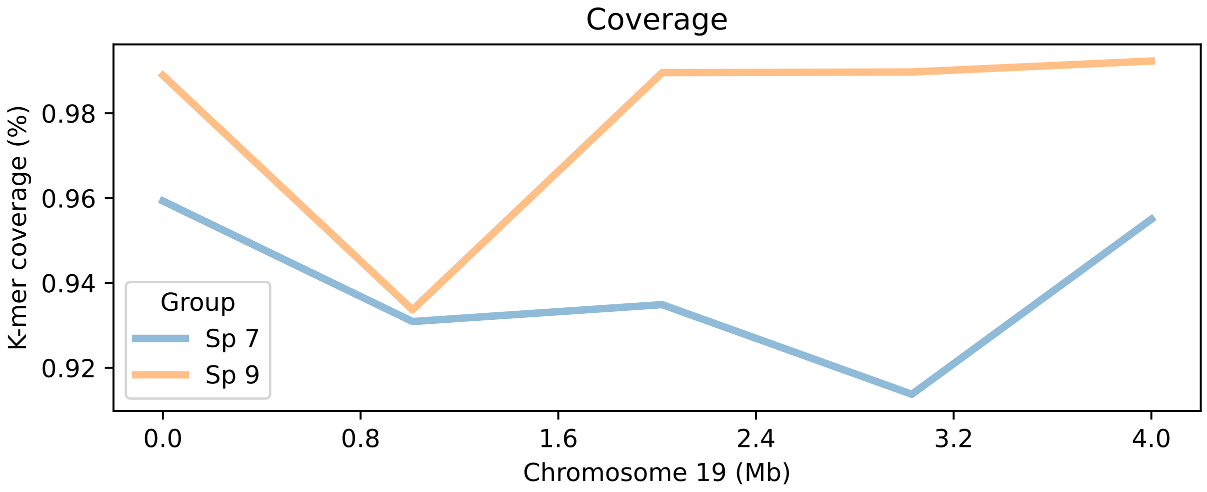

Fig. 7: K-mer anchoring across chromosome 19 in the Sp9509 reference, from two indexes.

To re-generate a plot with e.g. different styling, you can apply the `agplot` subcommand to the TSV file generated by `anchor-genome` when the `-t` / `--table` argument is provided. For example, to draw a new plot like the one above with the Sp19g00235 locus marked:

```
faidx -i chromsizes Sp_Chr19_genomes/Sp9509_oxford_v3_Chr19.fasta.gz > Sp9509_oxford_v3_
↪Chr19.chrom.sizes
pankmer agplot --groups Sp7 Sp9 --legend \
  --loci "Chr19:1625956:Sp19g00235" \
  --chromsizes Sp9509_oxford_v3_Chr19.chrom.sizes \
```

(continues on next page)

(continued from previous page)

```
  -t Sp_Chr19_genome_coverage/Sp9509_Chr19_anchor_multi.tsv \
  -o Sp_Chr19_genome_coverage/Sp9509_Chr19_anchor_Sp19g00235_locus.svg
```

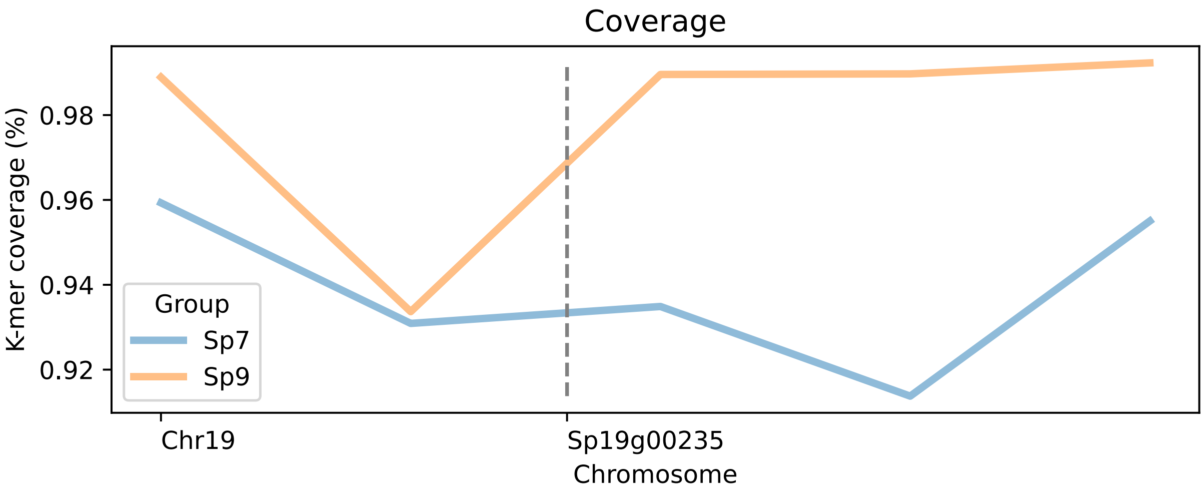

Fig. 8: K-mer anchoring across chromosome 19 in the Sp9509 reference, with Sp19g00235 locus marked.

#### 4.9 Calculate time elapsed while indexing

To calculate the time required to generate an index, you can use the `--time` option

```
pankmer index --time -g Sp_Chr19_genomes/ -o Sp_Chr19_index.tar
...
Indexed in 0.15 minutes
```

CHAPTER

FIVE

### USE CASE: HYBRIDIZATION EVENT IN *TYPHA*

#### 5.1 *Typha* super-pangenome dataset

For this use case we will apply PanKmer to a dataset of 10 *Typha domingensis* (TD) and 3 *Typha latifolia* (TL) genomes. The raw reads can be downloaded from NCBI SRA BioProject PRJNA742003 (see accession IDs in Table 1). The assembled genomes can be downloaded from AWS by:

```
curl -O https://salk-tm-pub.s3.us-west-2.amazonaws.com/pub-supplementary/PRJNA742003-
↪ASSEMBL.tar
```

Table 1: SRA accessions

| Experiment | Accession |
|---|---|
| TL01_WGS_ONT | SRX11987434 |
| TL01_WGS_ILL | SRX11987431 |
| TL05_WGS_ONT | SRX11987435 |
| TL05_WGS_ILL | SRX11977055 |
| TL17_WGS_ONT | SRX11987422 |
| TL17_WGS_ILL | SRX11977056 |
| TD01_WGS_ONT | SRX11987417 |
| TD01_WGS_ILL | SRX11977057 |
| TD03_WGS_ONT | SRX11987418 |
| TD03_WGS_ILL | SRX11977058 |
| TD04_WGS_ONT | SRX11987419 |
| TD04_WGS_ILL | SRX11977059 |
| TD07_WGS_ONT | SRX11987420 |
| TD07_WGS_ILL | SRX11977060 |
| TD08_WGS_ONT | SRX11987421 |
| TD08_WGS_ILL | SRX11987415 |
| TD12_WGS_ONT | SRX11987423 |
| TD12_WGS_ILL | SRX11987416 |
| TD22_WGS_ONT | SRX11987424 |
| TD22_WGS_ILL | SRX11977061 |
| TD23_WGS_ONT | SRX11987425 |
| TD23_WGS_ILL | SRX11977052 |
| TD25_WGS_ONT | SRX11987426 |
| TD25_WGS_ILL | SRX11977053 |
| TD36_WGS_ONT | SRX11987428 |
| TD36_WGS_ILL | SRX11977054 |

#### 5.2 Genome size and heterozygosity

We apply jellyfish 2[1] and genomescope 2[2] to calculate genome size and heterozygosity statistics (Table 2), and to plot k-mer spectra (Figure 1).

Table 2: Basic statistics of *Typha* genomes

| Name | Predicted genome size (Mb) | Heterozygosity (%) | N. contigs | GC content (%) |
|---|---|---|---|---|
| TD01 | 214 | 4.12 | 1743 | 37.7 |
| TD03 | 222 | 0.13 | 84 | 37.6 |
| TD04 | 226 | 0.45 | 578 | 37.7 |
| TD07 | 263 | 0.14 | 148 | 37.6 |
| TD08 | 213 | 0.05 | 93 | 37.7 |
| TD12 | 224 | 0.22 | 113 | 37.5 |
| TD22 | 218 | 2.60 | 1138 | 37.8 |
| TD23 | 213 | 2.60 | 1204 | 37.8 |
| TD25 | 210 | 0.12 | 130 | 37.7 |
| TD36 | 215 | 0.15 | 139 | 37.6 |
| TL01 | 234 | 0.07 | 89 | 37.8 |
| TL05 | 218 | 0.07 | 39 | 37.8 |
| TL17 | 215 | 0.07 | 46 | 37.7 |

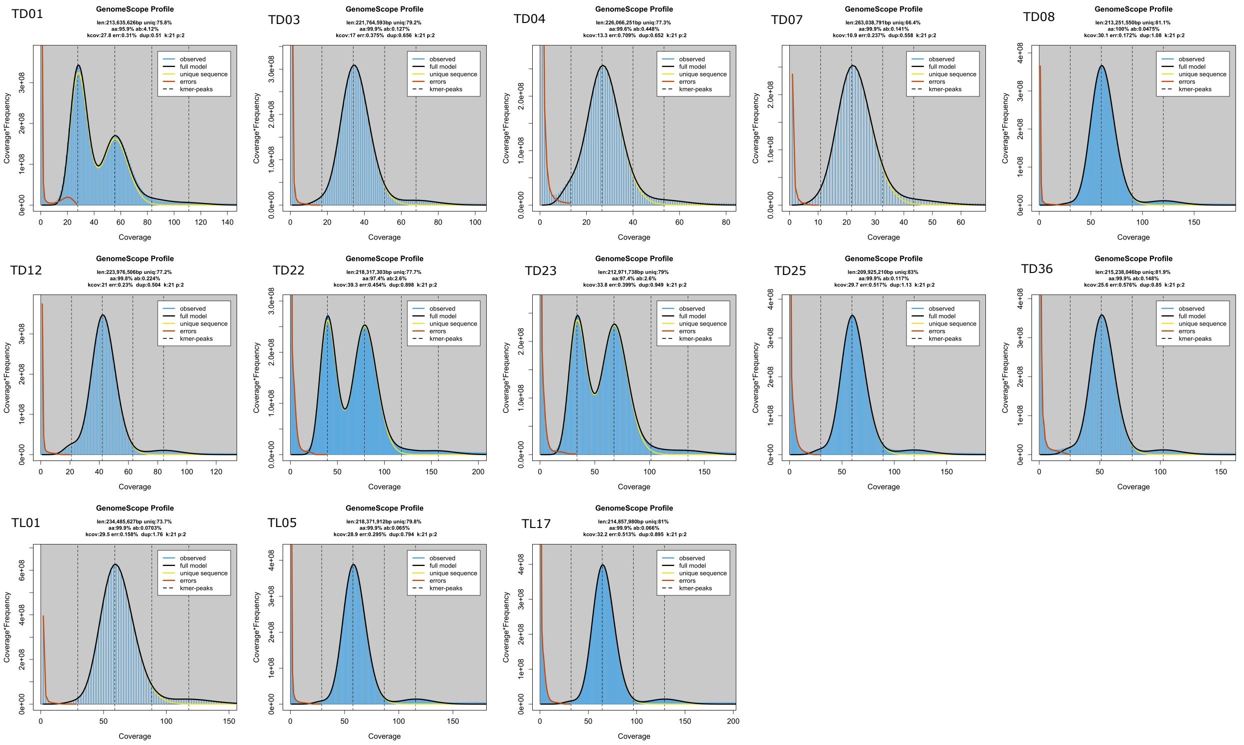

Fig. 1: *k*-mer spectra of Typha genomes. Bimodal distributions indicate heterozygosity.

[1] Guillaume Marcais and Carl Kingsford. A fast, lock-free approach for efficient parallel counting of occurrences of k-mers. *Bioinformatics* 2011

[2] Ranallo-Benavidez T.R. et al. GenomeScope 2.0 and Smudgeplot for reference-free profiling of polyploid genomes. *Nature Communications* 2020

Based on these results, we can conclude that TD01, TD22, and TD23 are heterozygous genotypes while the rest are homozygous.

#### 5.3 *k*-mer indexes

We use `pankmer index` to generate three *k*-mer indexes: *T. latifolia*, *T. domingensis*, and *Typha* super-pangenome. For example, to generate the super-pangenome, calculate an adjacency matrix, and plot a heatmap:

```
pankmer index -g PRJNA742003-ASSEMBL.tar -o typha-index.tar
pankmer adj-matrix -i typha-index.tar -o typha-matrix.csv
pankmer clustermap -i typha-matrix.csv -o typha-ani.svg --metric ani
```

The TD and TL indexes can be extracted from `typha-index.tar` with `pankmer subset`:

```
tar -xvf PRJNA742003-ASSEMBL.tar
pankmer subset -i typha-index.tar -g TD01 TD03 TD04 TD07 TD08 TD12 \
  TD22 TD23 TD25 TD36 -o td-index.tar
pankmer subset -i typha-index.tar -g TL01 TL05 TL17 -o tl-index.tar
```

By examining pairwise Jacard similarity values from each index, we can identify clusters of similar genomes and outliers. For example, TD01 is an outlier relative to the other TD genomes but is highly similar to the group of TL genomes.

We have observed that TD01 is highly heterozygous and also shares sequence with TL genomes. To inspect TD01 more closely, we proceed to *k*-mer conservation analysis.

#### 5.4 Genome anchoring

To anchor in TD01 and generate a plot of TD and TL conservation across contig 23:

```
pankmer anchor-genome -i td-index.tar tl-index.tar \
  -o TD01_23.svg -r TD01.v1.fa.gz -c 23 -t TD01_23.tsv \
  --groups "T. dom" "T. lat" --x-label Contig --legend \
  --legend-title Species --legend-loc outside --bin-size -1
```

Applying `pankmer anchor-genome` to 26 contigs of TD01 that are larger than 2 Mb reveals the presence of large and small introgressions of TL sequence into TD01 (Figure 3). Contigs 23 and 26 are represenative examples (Figure 4). These results suggest a recent TD-TL hybridization event resulting in TD01.

#### 5.5 Appendix: Resource usage

Pangenome was built with 20 threads and 16 memory blocks

Table 3: Resource usage for indexing *Typha* genomes

| Species | n-genomes | peak_mem_gb | time_minutes |
|---|---|---|---|
| *T. latifolia* | 3 | | |
| *T. domingensis* | 10 | | |
| *Typha* super-pangenome | 13 | | |

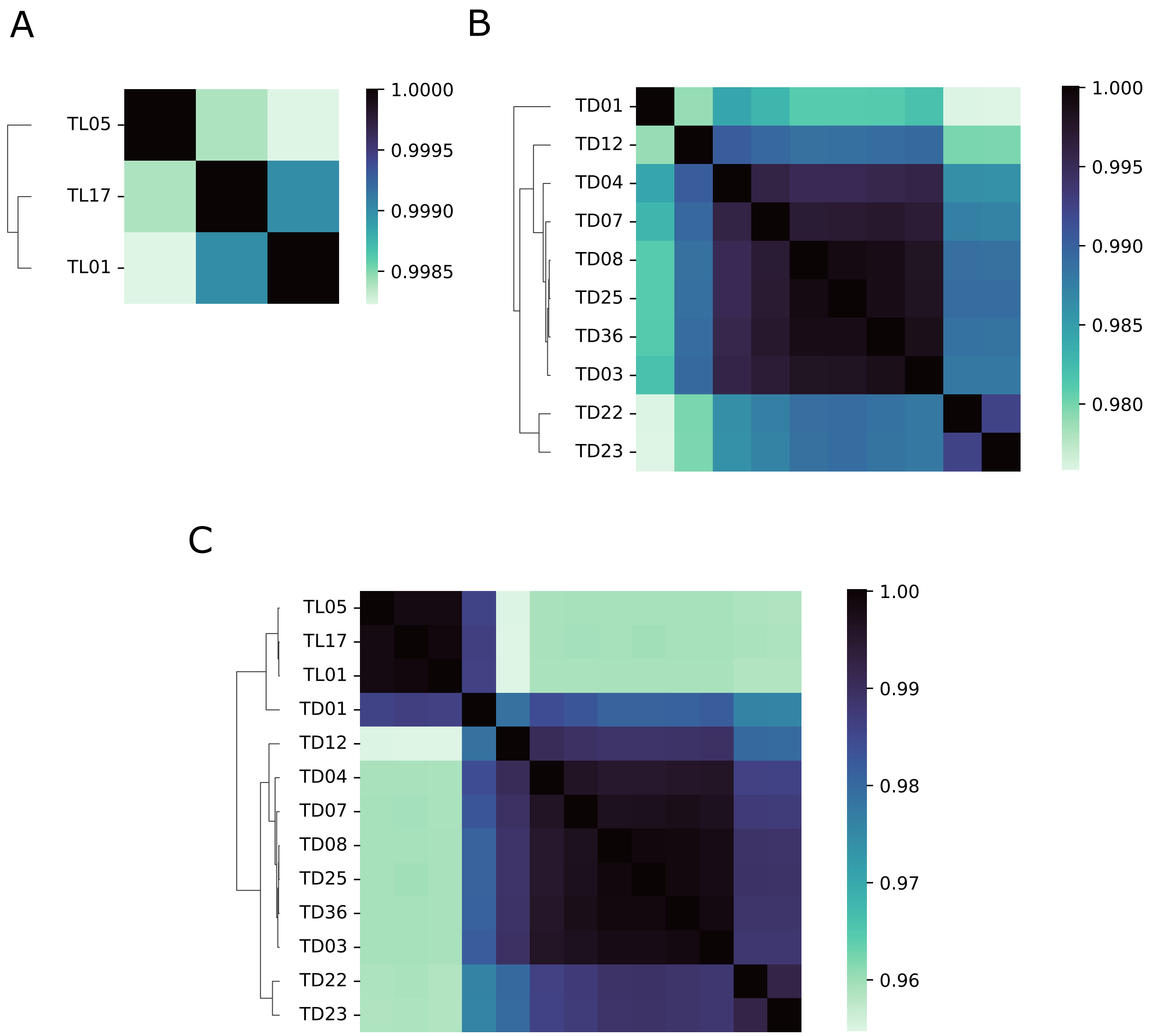

Fig. 2: Pairwise Average Nucleotide Identity values across Typha pangenomes. A) *T. latifolia* pangenome B) *T. domingensis* pangenome C) Typha super-pangenome

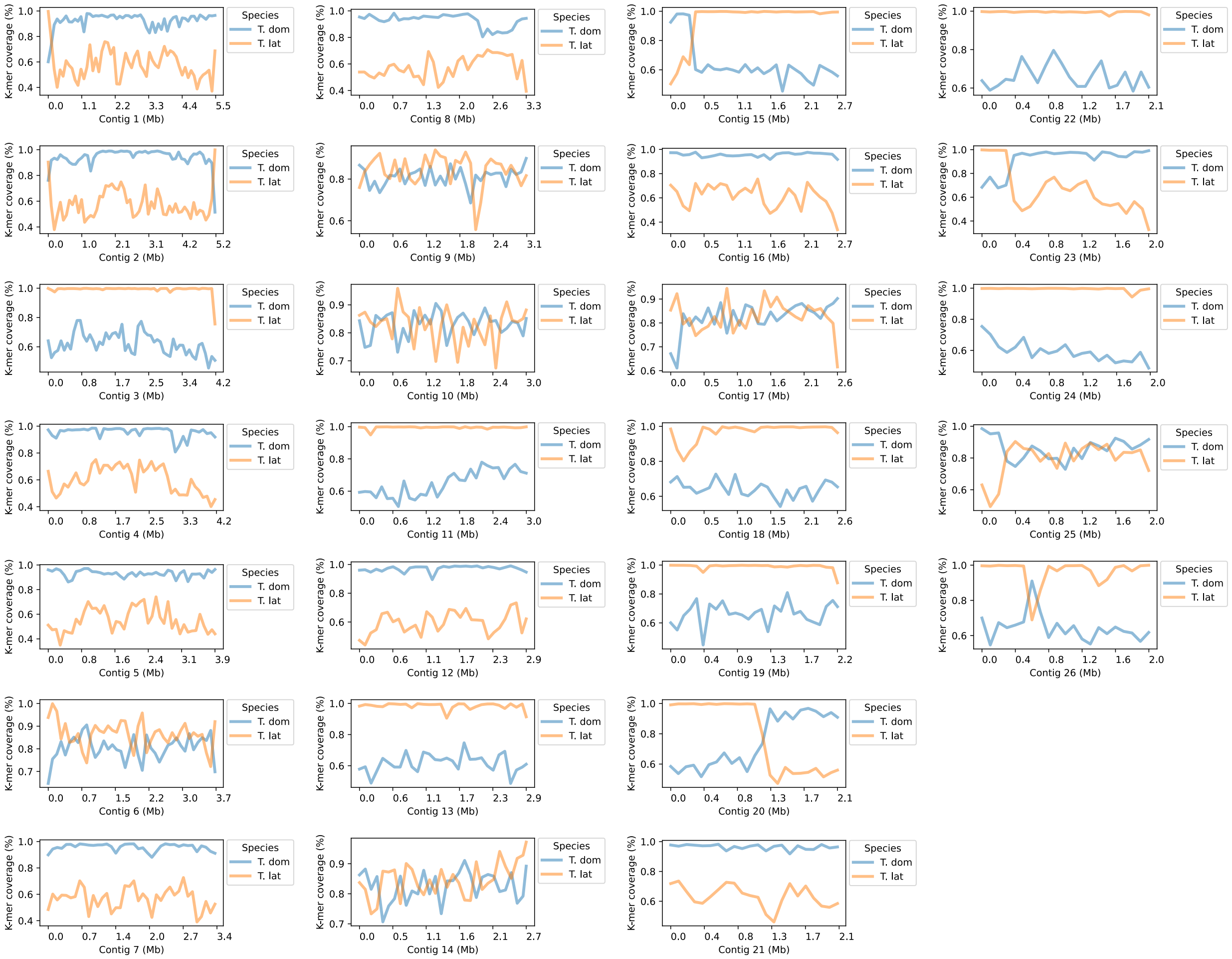

Fig. 3: k-mer conservation of TD01 contigs.

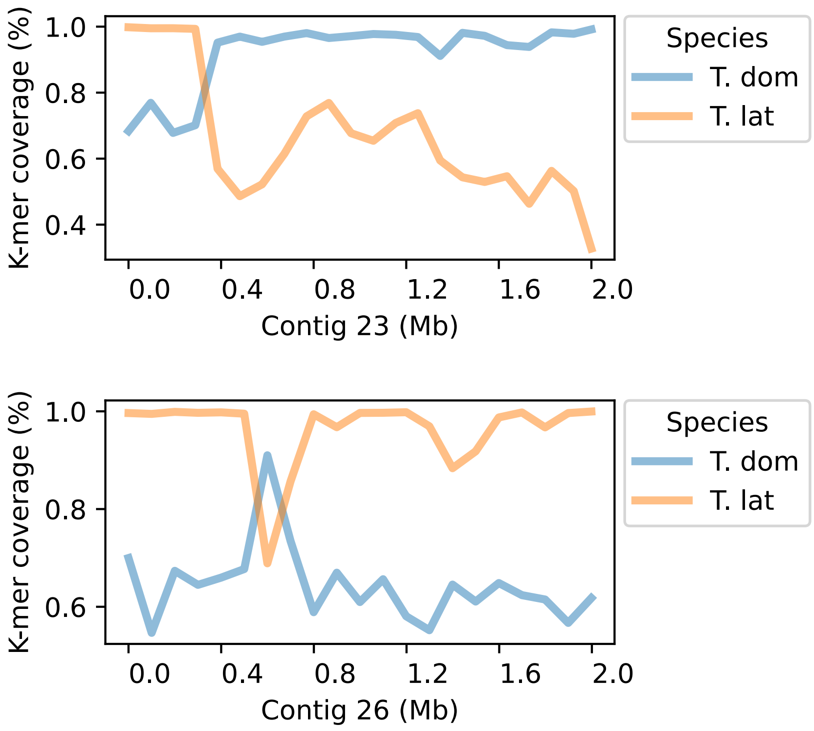

Fig. 4: Representative contigs illustrating large and small introgressions

CHAPTER

SIX

### LARGE PANGENOMES

When building a pangenome of more than a handful of input genomes, you may wish to adjust some settings to save on resource usage.

#### 6.1 *k*-mer counting rounds

A challenge of building the PanKmer index from a large set of eukaryote genomes is the significant memory footprint of the operation. PanKmer includes the `--rounds` option to regulate peak memory usage. The default value of `--rounds` is 1, which means each input genome will be read once by each thread and the entire index will be built on the first pass. Increasing the value will divide the *k*-mer space into discrete blocks and construct a sub-index for each block, in sequence, finally merging all sub-indexes into the complete index. When the number of rounds is greater than 1, this scheme reduces peak memory usage by reducing the number of *k*-mers held in memory, at the expense of increased runtime since multiple *k*-mer decomposition passes must be made over each genome. Hence, when the pangenome size is small or large memory resources are available, fewer rounds are recommended, while more rounds are recommended for large pangenomes or environments with minimal memory.

```
pankmer index --rounds <nrounds> -g <genomes> -o <index[.tar]>
```

Estimate the appropriate number of rounds based on the size of your pangenome and your available memory resources. See the benchmarks below for a starting point. Increasing the number of rounds will decrease memory usage at the expense of longer runtime, while decreasing the number of rounds will decrease runtime at the expense of higher memory usage. Pankmer can use up to hundreds of rounds to achieve low memory usage for large pangenomes.

#### 6.2 GZIP compression

Depending on your pangenome size and computing resources, you may wish to adjust the gzip compression level of indexing. You can do this by:

```
pankmer index --gzip-level <level> -g <genomes> -o <index[.tar]>
```

The gzip level is an integer from 1-9, with a default value of 6. Setting a lower gzip level will shorten runtime but will slightly increase the size of the PanKmer index on disk. Setting a higher gzip level will slightly decrease the size of the index but will increase runtime. For fastest indexing usage `--gzip-level 1`.

#### 6.3 Index completeness

By default, PanKmer records *every* *k*-mer observed in the input sequences, producing an index that is "*k*-mer complete". This can be redundant for some applications. In such cases, you can use the `-q/--qual` argument to reduce completeness of the index.

```
pankmer index -q <Q> -g <genomes> -o <index[.tar]>
pankmer index -q 30 -g <genomes> -o <index[.tar]>
```

In a "Q30" PanKmer index, 80% of k-mers will be discarded instead of recorded. Approx 999 of every 1000 base pairs of input sequence will still be represented in at least 1 recorded k-mer. This allows a significant reduction in required memory and runtime, with only a slight loss in fidelity.

Alternatively, set the fraction of k-mers directly:

```
pankmer index --fraction <float> -g <genomes> -o <index[.tar]>
```

#### 6.4 Benchmarks

PanKmer was benchmarked on several pangenome datasets of various sizes.

1. The Typha super-pangenome dataset presented in the *use case* (13 genomes total)
2. *Solanum* super-pangenome (46 genomes total)
3. *Zea* super-pangenome (54 genomes total)
4. *H. sapiens* pangenome (94 haplotypes total)
5. *A. thaliana* pseudo-pangenome (1,135 pseudo-genomes total)

These datasets are further described in *datasets*. Memory usage and runtime were also compared to Kmer-db . Shell scripts to execute all PanKmer-only benchmarking runs and all comparative PanKmer vs Kmer-db benchmarking runs can be found in the PanKmer GitLab repository .

All pankmer runs were parametrized to execute with 20 threads (`--threads 20`), default gzip level (6), and a *k*-mer complete index result. In all cases except the *H. sapiens* pangenome, the input genomes were compressed. The hard-coded *k*-mer size of 31 bp was in place for all cases. For PanKmer only runs, the `--rounds` parameter was set either to 16 for moderate memory usage with moderate runtimes or to 256 for low memory usage with long runtimes. For PanKmer vs Kmer-db runs, `--rounds` was set to 1 for highest memory usage with shortest runtimes. All Kmer-db runs had parameters `-k 30` to use the maximum k-mer size of 30 and either `-t 20` or `-t 4` to use either 20 or 4 threads.

**Computing environment**

The computer used for benchmarking was of the following configuration:

- 2 Intel Xeon Gold 6326 CPUs, 16 double-threaded cores per CPU, clocked at 2.9 GHz
- 1024 GB RAM
- 12 HDDs of size 2.4 TB each, hdparm -t reported buffered read speed 2317.27 MB/sec
- Ubuntu 20.04 x86-64 OS

PanKmer was compiled with Rust version 1.71.0. Kmer-db binary was installed via bioconda .

Table 1: Sizes of pangenome datasets

| Clade | N. genomes | Mean genome size (Mb) | Total 31-mers ($10^6$) |
|---|---|---|---|
| *Typha* | 13 | 222 | 542 |
| *Solanum* | 46 | 807 | 1181 |
| *Zea* | 54 | 2426 | 8266 |
| *H. sapiens* | 94 | 3015 | 3188 |
| *A. thaliana* | 1135 | 119 | 306 |

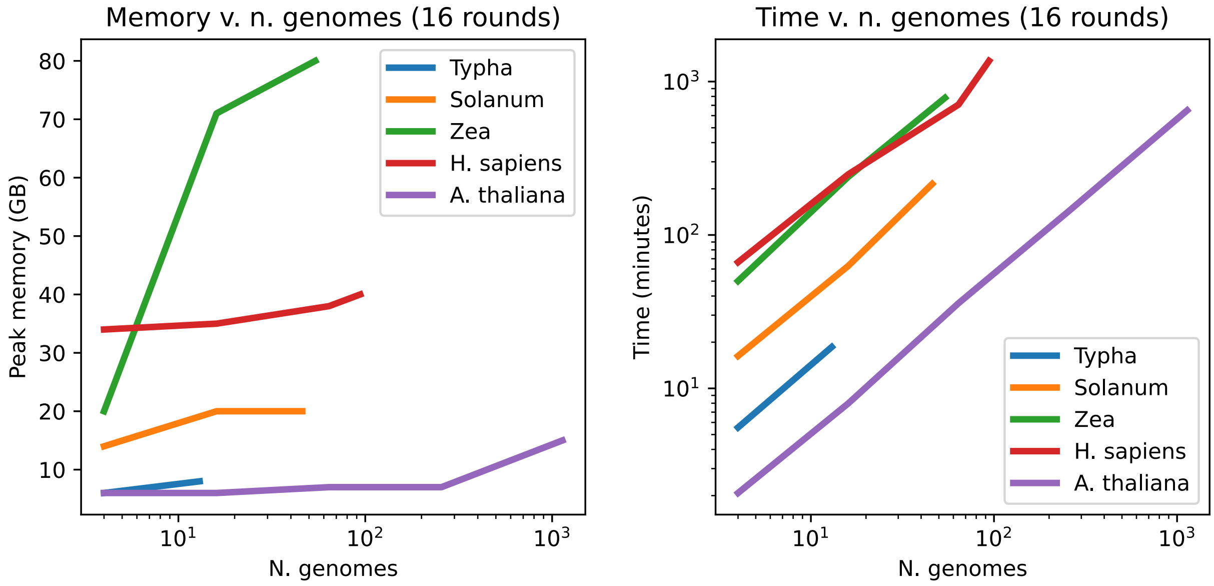

Fig. 1: Benchmark results with 16 rounds

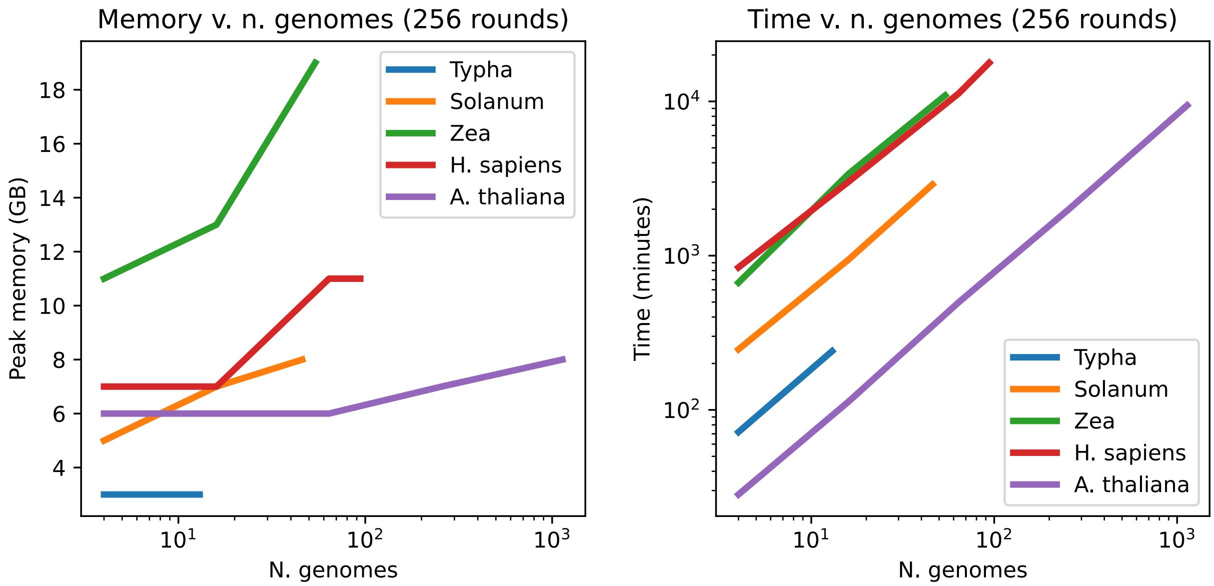

Fig. 2: Benchmark results with 256 rounds

Table 2: Resource usage of PanKmer indexing (v0.17.0) with 16 rounds

| Clade | N. genomes | Peak memory (GB) | Time (minutes) |
|---|---|---|---|
| *Typha* | 4 | 6 | 5.56 |
| *Typha* | 13 | 8 | 18.61 |
| *Solanum* | 4 | 14 | 16.26 |
| *Solanum* | 16 | 20 | 62.87 |
| *Solanum* | 46 | 20 | 215.19 |
| *Zea* | 4 | 20 | 50.12 |
| *Zea* | 16 | 71 | 239.38 |
| *Zea* | 54 | 80 | 783.10 |
| *H. sapiens* | 4 | 34 | 66.45 |
| *H. sapiens* | 16 | 35 | 248.50 |
| *H. sapiens* | 64 | 38 | 707.39 |
| *H. sapiens* | 94 | 40 | 1364.33 |
| *A. thaliana* | 4 | 6 | 2.08 |
| *A. thaliana* | 16 | 6 | 7.98 |
| *A. thaliana* | 64 | 7 | 35.90 |
| *A. thaliana* | 256 | 7 | 142.93 |
| *A. thaliana* | 1135 | 15 | 648.90 |

Table 3: Resource usage of PanKmer indexing (v0.17.0) with 256 rounds

| Clade | N. genomes | Peak memory (GB) | Time (minutes) |
|---|---|---|---|
| *Typha* | 4 | 3 | 71.87 |
| *Typha* | 13 | 3 | 240.40 |
| *Solanum* | 4 | 5 | 246.94 |
| *Solanum* | 16 | 7 | 943.86 |
| *Solanum* | 46 | 8 | 2893.45 |
| *Zea* | 4 | 11 | 668.40 |
| *Zea* | 16 | 13 | 3391.95 |
| *Zea* | 54 | 19 | 10930.97 |
| *H. sapiens* | 4 | 7 | 839.12 |
| *H. sapiens* | 16 | 7 | 3015.26 |
| *H. sapiens* | 64 | 11 | 11268.46 |
| *H. sapiens* | 94 | 11 | 17800.85 |
| *A. thaliana* | 4 | 6 | 28.28 |
| *A. thaliana* | 16 | 6 | 113.29 |
| *A. thaliana* | 64 | 6 | 498.88 |
| *A. thaliana* | 256 | 7 | 2000.42 |
| *A. thaliana* | 1135 | 8 | 9425.92 |

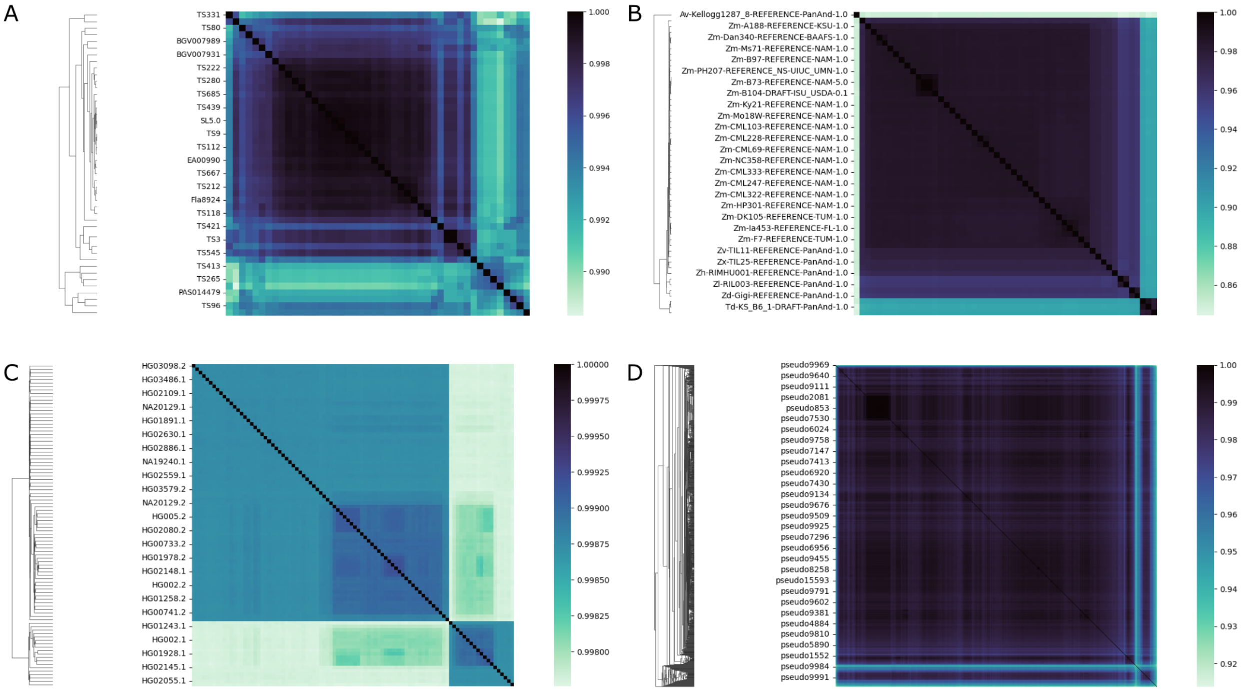

Fig. 3: Heatmaps of PanKmer benchmarking pangenome datasets. A) *Solanum* B) *Zea* C) *H. sapiens* D) *A. thaliana*

#### 6.5 PanKmer vs Kmer-db

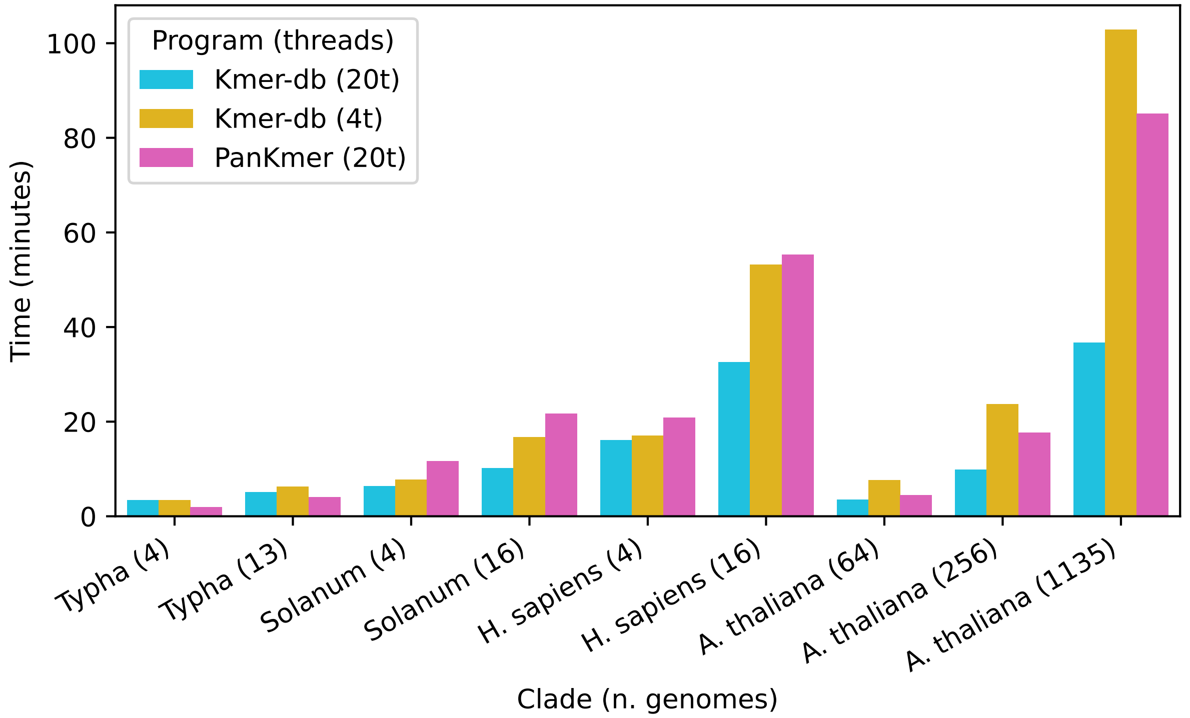

Fig. 4: Kmer-db vs PanKmer runtime

Table 4: Resource usage of Kmer-db (20 threads)

| Clade | N. genomes | Peak memory (GB) | Time (minutes) |
|---|---|---|---|
| *Typha* | 4 | 72 | 3.45 |
| *Typha* | 13 | 88 | 5.15 |
| *Solanum* | 4 | 110 | 6.40 |
| *Solanum* | 16 | 186 | 10.17 |
| *H. sapiens* | 4 | 281 | 16.17 |
| *H. sapiens* | 16 | 535 | 32.65 |
| *A. thaliana* | 64 | 81 | 3.59 |
| *A. thaliana* | 256 | 89 | 9.88 |
| *A. thaliana* | 1135 | 107 | 36.73 |

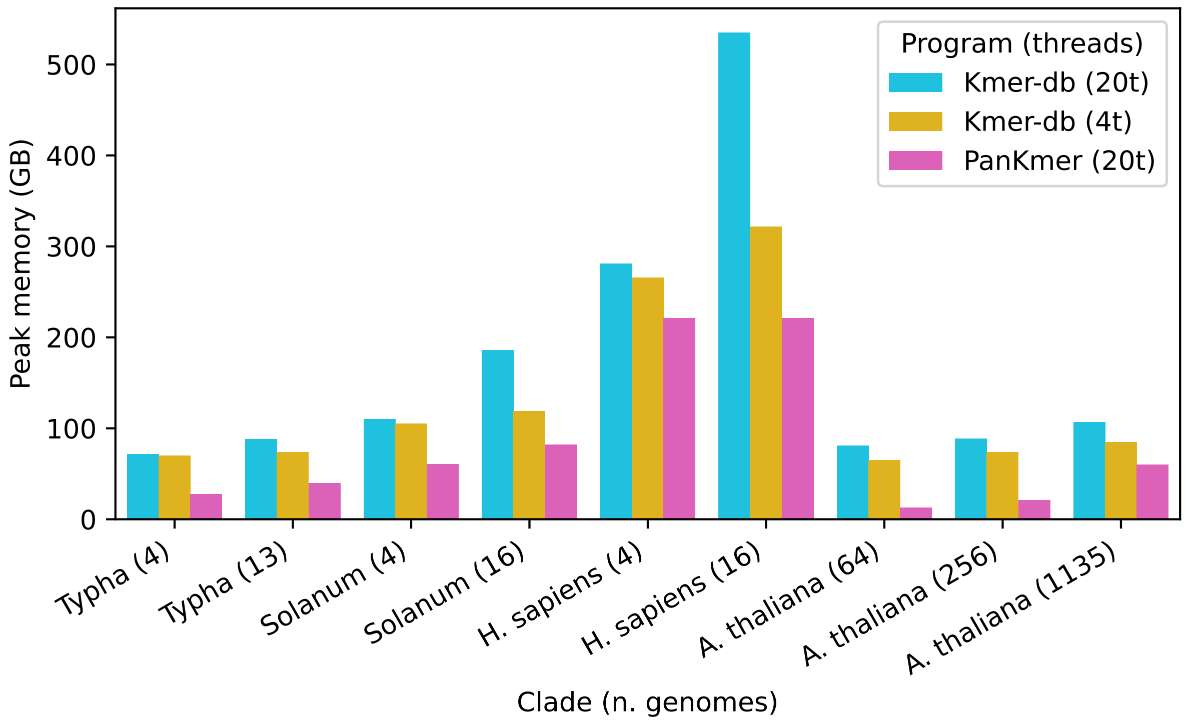

Fig. 5: Kmer-db vs PanKmer peak memory usage

Table 5: Resource usage of Kmer-db (4 threads)

| Clade | N. genomes | Peak memory (GB) | Time (minutes) |
|---|---|---|---|
| *Typha* | 4 | 70 | 3.45 |
| *Typha* | 13 | 74 | 6.28 |
| *Solanum* | 4 | 105 | 7.80 |
| *Solanum* | 16 | 119 | 16.73 |
| *H. sapiens* | 4 | 266 | 17.08 |
| *H. sapiens* | 16 | 322 | 53.25 |
| *A. thaliana* | 64 | 65 | 7.65 |
| *A. thaliana* | 256 | 73 | 23.78 |
| *A. thaliana* | 1135 | 85 | 102.87 |

Table 6: Resource usage of PanKmer (1 round, 20 threads)

| Clade | N. genomes | Peak memory (GB) | Time (minutes) |
|---|---|---|---|
| *Typha* | 4 | 28 | 1.95 |
| *Typha* | 13 | 40 | 4.10 |
| *Solanum* | 4 | 61 | 11.67 |
| *Solanum* | 16 | 82 | 21.76 |
| *H. sapiens* | 4 | 221 | 20.85 |
| *H. sapiens* | 16 | 221 | 55.32 |
| *A. thaliana* | 64 | 13 | 4.49 |
| *A. thaliana* | 256 | 21 | 17.72 |
| *A. thaliana* | 1135 | 60 | 85.13 |

#### 6.6 Appendix: details of memory block scheme

A basic action of PanKmer is to read k-mers from a genome into memory

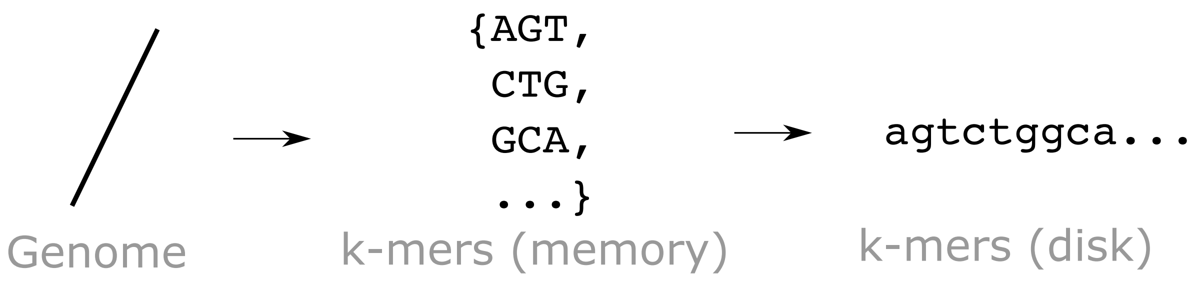

Fig. 6: Simple k-mer counting

However, doing this naively results in excesive memory usage

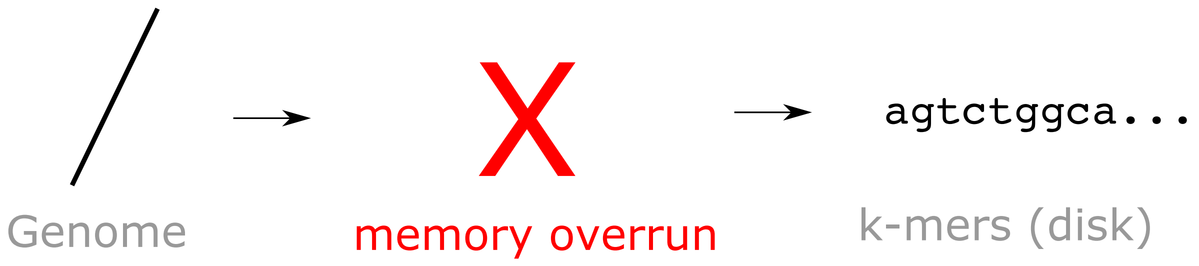

Fig. 7: Memory overrun

As a solution, PanKmer breaks up the space of k-mers into blocks and operates on them sequentially. This limits the peak memory requirement. The number of blocks is determined by the `--split-memory` argument.

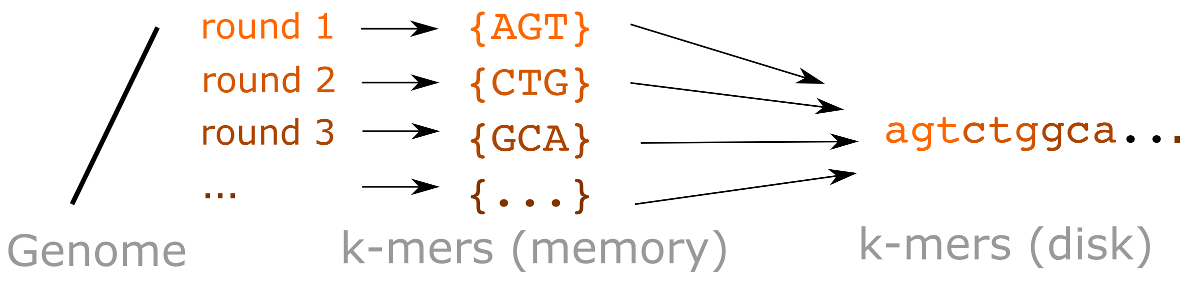

Fig. 8: K-mer counting with memory blocks

Adding multiple genomes and scores fits naturally into this scheme

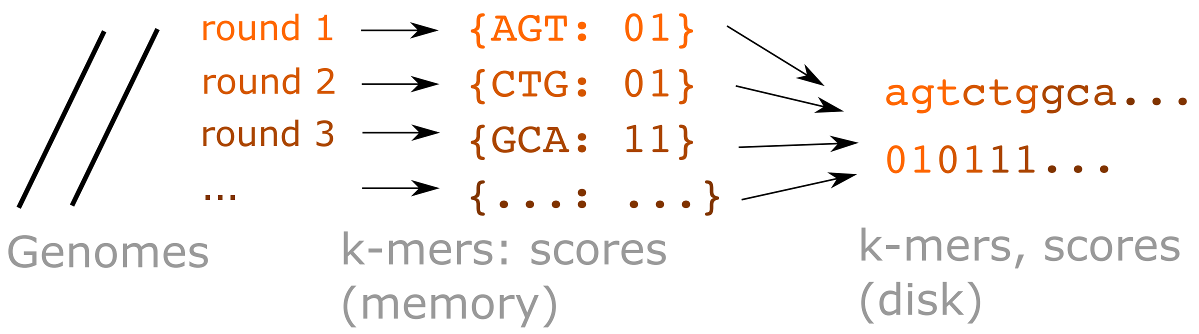

Fig. 9: K-mer counting with memory blocks and scores

CHAPTER

SEVEN

### PANGENOME DATASETS

The `pankmer download-example` subcommand can be used to download genomes from several publicly available pangenome datasets. See the help text:

```
pankmer download-example --help
```

```
usage: pankmer download-example [-h] [-d <dir/>] [-s {Spolyrhiza,Slycopersicum,Zmays,
↪Hsapiens,Bsubtilis,Athaliana}] [-n <int>]

options:
  -h, --help            show this help message and exit
  -d <dir/>, --dir <dir/>
                        destination directory for example data
  -s {Spolyrhiza,Slycopersicum,Zmays,Hsapiens,Bsubtilis,Athaliana}, --species
↪{Spolyrhiza,Slycopersicum,Zmays,Hsapiens,Bsubtilis,Athaliana}
                        download publicly available genomes. Species: max_samples.␣
↪Spolyrhiza: 3, Slycopersicum: 46, Zmays: 54, Hsapiens: 94,    Bsubtilis: 164,␣
↪Athaliana: 1135
  -n <int>, --n-samples <int>
                        number of samples to download, must be less than species max [1]
```

The `-s/--species` option selects the species, and the `-n/--n-samples` option selects the number of samples to download. The maximum number of samples for each species is:

Table 1: Pangenome datasets

| Species | Max samples |
| --- | --- |
| S. polyrhiza | 3 |
| S. lycopersicum | 46 |
| 26. mays | 54 |
| H. sapiens | 94 |
| B. subtilis | 164 |
| A. thaliana | 1135 |

See below a description of each pangenome dataset

#### 7.1 *S. lycopersicum*

46 *Solanum lycopersicum* genomes from the SolOmics database. See also *Nature* article: Graph pangenome captures missing heritability and empowers tomato breeding .

#### 7.2 *Z. mays*

54 *Zea mays* genomes from the downloads page of MaizeGDB.

#### 7.3 *H. sapiens*

94 *Homo sapiens* haplotypes from Year 1 of the Human Pangenome Reference Consortium / Human Pangenome Project. Download details found at the HPRC/HPP github repository. See also *Nature* articles: A draft human pangenome reference , The Human Pangenome Project: a global resource to map genomic diversity.

#### 7.4 *B. subtilis*

164 *B. subtilis* genomes from NCBI.

#### 7.5 *A. thaliana*

1135 *Arabidopsis thaliana* pseudo-genomes from the data center of 1001 Genomes. See also *Cell* article: 1,135 Genomes Reveal the Global Pattern of Polymorphism in Arabidopsis thaliana .

#### 7.6 *S. polyrhiza*

A collection of 3 *Spirodela polyrhiza* clones Sp7498, Sp9509, Sp9512, from the following sources: Sp7498 and Sp9509 sequences were sourced from the following references found at http://spirodelagenome.org:

Sp9509_oxford_v3
NCBI: GCA_900492545.1
CoGe: id51364
This genome was generated with Oxford Nanopore and polished with Illumina, scaffolded against the previous Illumina-based genome Sp9509v3 and validated with BioNano optical maps and multi-color FISH (mcFISH).

Hoang PNT, Michael TP, Gilbert S, Chu P, Motley TS, Appenroth KJ, Schubert I, Lam E. Generating a high-confidence reference genome map of the Greater Duckweed by integration of cytogenomic, optical mapping and Oxford Nanopore technologies. Plant J. 2018 Jul 28.

Sp7498_HiC
CoGe: 55877
This assembly was generated using Oxford Nanopore long reads and Illumina-based HiC scaffolding.

Harkess A, McGlaughlin F, Bilkey N, Elliott K, Emenecker R, Mattoon E, Miller K, Vierstra R, Meyers BC, Michael TP. High contiguity Spirodela polyrhiza genomes reveal conserved chromosomal structure. Submitted.

Sp9512 sequence was sourced from research data for the following in-progress publication:

Pasaribu B, Acosta K, Aylward A, Abramson BW, Colt K, Hartwick NT, Liang Y, Shanklin J, Michael TP, Lam E Genomics of turions from the Greater Duckweed reveal pathways for tissue dormancy and reemergence strategy of an aquatic plant.

Sp9512 can be downloaded from Michael lab AWS storage.

CHAPTER

EIGHT

### RAW SEQUENCING READS

In addition to assemblies in FASTA format, PanKmer can also operate on raw sequencing reads in FASTQ format. They are handled differently to account for the possibility of sequencing errors.

When FASTQ files are included as input genomes, the indexing process is altered so that, during indexing, each k-mer's score is composed of 2 bits per genome instead of 1 bit. This increases the memory footprint but allows tracking whether each k-mer has been observed one, 2, or >2 times in each genome.

### CHAPTER NINE

### INDEXING ALGORITHM

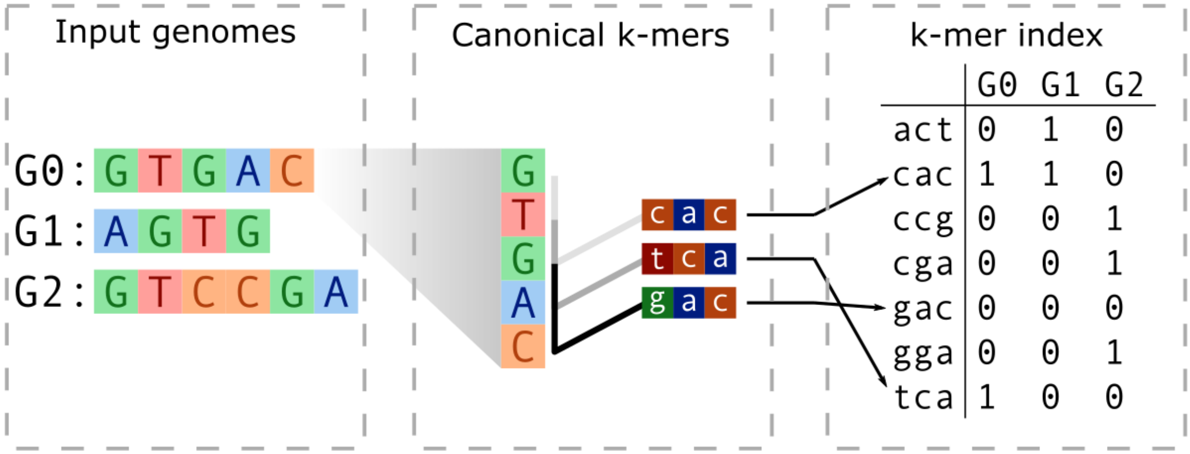

Fig. 1: Diagram of PanKmer functionality.

#### 9.1 *k*-mer decomposition

PanKmer indexing breaks down genomes to unique canonical $k$-mers ($k = 31$), and assigns each $k$-mer a bit vector indicating which samples include it. K-mers are represented as integers as in k-mer counters such as Jellyfish[1] and Kmer-db[2]. For each $k$-mer in a contig, we encode the $k$-mer and its complement as unsigned 64-bit integers. The larger integer (lexicographically smaller $k$-mer) is selected as the canonical $k$-mer. Each base has a two bit substitute:

Table 1: Nucleotide base to 2-bit integer substitute

| Base | 2-bit integer |
|---|---|
| A | 11 |
| C | 10 |
| G | 01 |
| T | 00 |

For example, the 5-mer ACGTC becomes 11-10-01-00-10 which equals 1938 in base 10. The reverse complement GACGT then becomes 01-11-10-01-00 which equals 1508 in base 10. Since 1938 is greater than 1508, it becomes the

[1] Guillaume Marçais , Carl Kingsford, A fast, lock-free approach for efficient parallel counting of occurrences of k-mers, Bioinformatics, Volume 27, Issue 6, March 2011

[2] Deorowicz, S., Gudyś, A., Długosz, M., Kokot, M., Danek, A. (2019) Kmer-db: instant evolutionary distance estimation, Bioinformatics

representative of the two, and is added to the index. Any $k$-mers containing a base that is not one of A, C, G, or T are skipped.

Each $k$-mer is added/updated in a hash table of $k$-mers wherein the keys are $k$-mers and the values are bit vectors called *scores*. A score indicates presence or absence of the $k$-mer in each sample. Scores are $n$ bits where $n$ is the number of samples. Each bit indicates presence/absence with sample 1 at the right most bit and sample $n$ at the left most bit. For example, for a sample set of four genomes:

Table 2: Example score values

| Bit Score | Sample 4 | Sample 3 | Sample 2 | Sample 1 |
|---|---|---|---|---|
| 0011 | Not present | Not present | Present | Present |
| 1101 | Present | Present | Not present | Present |

After all $k$-mers in all contigs have been updated in the hash table, the $k$-mers and scores are written to their corresponding files on disk, along with a metadata file, as shown in the *tutorial*.

#### 9.2 Choice of *k*

PanKmer's implementation represents each k-mer as an unsigned 64-bit integer, as in k-mer counting tools such as Jellyfish and Kmer-db[Page 39, 1][Page 39, 2] . Hence the maximum value of $k$ is 32. Choosing $k \geq 31$ limits the theoretical rate of non-unique k-mers occurring by chance to fewer than 1 per 100 million k-mers[4]. This is important because some of PanKmer's downstream analysis functions, such as genome anchoring, work best when the non-unique k-mer rate is low. Hence, $k = 31$ and $k = 32$ are both suitable values, and PanKmer uses $k = 31$. Furthermore, 31-bp $k$-mers have been applied successfully to define variation in previous studies[3][5][6].

#### 9.3 Assembly quality considerations

The $k$-mer index is robust to varying contiguity of the input assemblies. Chromosome-level assemblies can be compared directly to unscaffolded contigs or even unaligned reads. By the same token, the index is agnostic to the choice of software tools used in the assembly process. However, PanKmer can be confounded by genome sequence datasets that either:

1. Fail to include large sections of the sequenced genome
2. Contain a large number of sequencing errors

In the first case, many $k$-mers will be missing from some genomes in the index. In the second case, many spurious $k$-mers will be injected into some genomes. So long as the input files do not suffer from extensive sequencing errors or missing reads, the index is agnostic to the choice of assembly pipeline or the contiguity of assemblies.

[4] Sheikhizadeh, S., Schranz, M. E., Akdel, M., de Ridder, D. & Smit, S. PanTools: representation, storage and exploration of pan-genomic data. Bioinformatics 32, i487–i493 (2016).

[3] Rahman, A., Hallgrímsdóttir, I., Eisen, M. & Pachter, L. Association mapping from sequencing reads using k-mers. eLife 7, e32920 (2018).

[5] Voichek, Y. & Weigel, D. Identifying genetic variants underlying phenotypic variation in plants without complete genomes. *Nat. Genet.* 52, 534-540 (2020).

[6] Karikari, B., Lemay, M.-A. & Belzile, F. k-mer-Based Genome-Wide Association Studies in Plants: Advances, Challenges, and Perspectives. *Genes* 14, 1439 (2023).

#### 9.4 Numerical complexity, resource usage, & runtime

Indexing has an $O(n \times c)$ memory footprint, where $c$ is the number of unique canonical *k*-mers. To reduce the memory usage, the *k*-mer space can be partitioned based on the leading two bases of the canonical *k*-mers. Since we choose the canonical *k*-mer to have a larger value, the *k*-mer space skews to start with AA rather than TT. Thus, the *k*-mer space can be divided into the following groups or any combination thereof: (TT, TG, TC, TA), (GT, GG, GC, GA), (CT, CG), (CC, CA), (AT), (AG), (AC), (AA).

The conversion from contig to *k*-mers to a canonical representative has $O(b)$ complexity where $b$ is the sum of the genome sizes. The canonical *k*-mers must also be added to the main *k*-mer hash table, which also has $O(b)$ complexity.

CHAPTER

TEN

### LIMITATIONS

#### 10.1 *k*-mers and genetic variants

The presence or absence of 31-mers can be used to detect most cases of SNP, INDEL, and SV including deletion, insertion, inversion, and translocation. See below illustrations of how it works. These figures were published in[1] and[2]
.

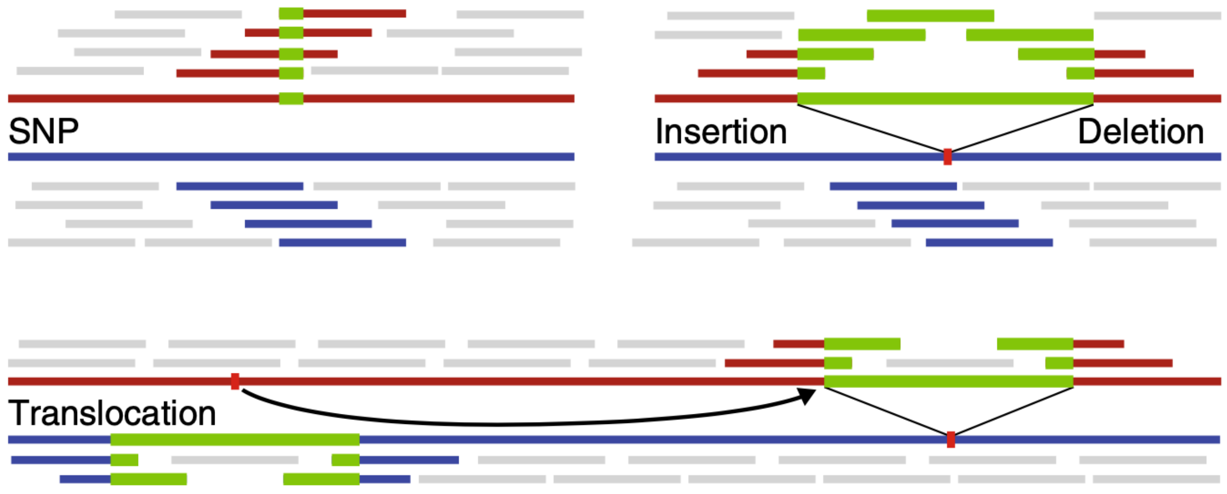

Fig. 1: **k-mers and genetic variants.** The blue and red lines represent two individual genomes. The colored short bars mark *k*-mers unique to each genome, and the gray bars mark *k*-mers shared between genomes. From[1]

[1] Voichek, Y. & Weigel, D. Identifying genetic variants underlying phenotypic variation in plants without complete genomes. *Nat. Genet.* 52, 534-540 (2020).

[2] Karikari B, Lemay M-A, Belzile F. k-mer-Based Genome-Wide Association Studies in Plants: Advances, Challenges, and Perspectives. Genes. 2023; 14(7):1439.

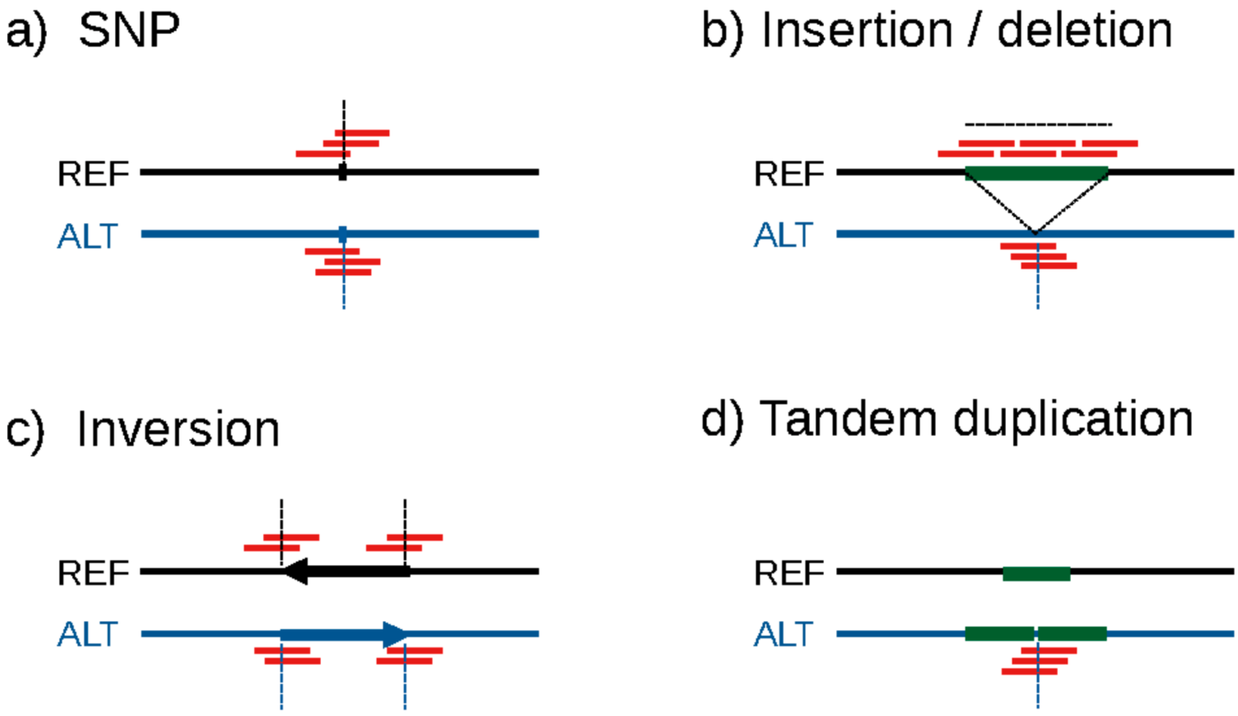

Fig. 2: **k-mers and genetic variants.** Illustration of the location of unique *k*-mers (red lines) originating from a reference genome (solid black lines) and an alternate genome (solid blue lines) depending on the underlying variant type. Dashed lines indicate genomic locations (vertical lines) or ranges of genomic positions (horizontal lines) that will induce unique *k*-mer patterns in the respective sample. (a) A single-nucleotide polymorphism (SNP, indicated by ticks on the genomes) will result in *k*-mers specific to each genome when *k*-mers overlap with the SNP. (b) For insertions/deletions, unique *k*-mers will originate from the breakpoints induced by the variation. In addition, if the sequence insertion is novel (i.e., not found elsewhere in the genome), unique *k*-mers will also originate from within the inserted sequence. (c) For inversions, unique *k*-mers arise at the inversion breakpoints. *k*-mers originating from within the inverted sequence will not differentiate between the two genomes because only the orientation of the sequence will differ. (d) For tandem duplications, novel adjacencies will only occur in the genome bearing the additional copies. From[2]

#### 10.2 Copy number variation

PanKmer and similar methods can be confounded by copy number variations in repetitive sequences. *k*-mer presence/absence cannot tag CNVs unless their junctions produce unique K-mers. In practice, while some special cases can be tagged, a large class of CNVs are invisible to PanKmer. See an illustrative example below:

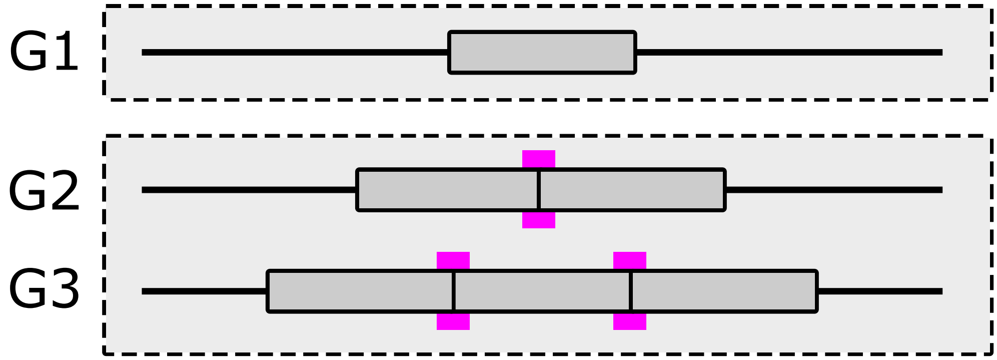

Fig. 3: **Limitations of k-mers for tagging copy number variation.** Genomes G1, G2, and G3 have 1, 2, and 3 tandem copies of a repeat, respectively. K-mers produced by the copy junction highlighted in magenta can distinguish G2 and G3 from G1. However, G2 cannot be distinguished from G3 since the junction is present in both.

CHAPTER

ELEVEN

### TESTS

To run PanKmer's unit tests, download the git repository and use pytest. This assumes you have a conda environment set up as in *Installation*:

```
conda activate pankmer
conda install -c conda-forge pytest fq
git clone https://gitlab.com/salk-tm/pankmer.git
cd pankmer
pip install .
pytest
```

The unit tests are divided into three categories: fast, slow, and very slow. The fast tests have a short runtime and do not require an internet connection. The slow tests have more complete coverage of PanKmer's features, but require an internet connection to download the example dataset and use it as input. The very slow tests have the same requirements as the slow tests but are even slower.

By default, only the fast tests are run. You can run the slow tests by adding the `--runslow` option:

```
pytest --runslow
```

To run the slow and very slow tests, use the `--veryslow` option:

```
pytest --veryslow
```

CHAPTER

### TWELVE

### CLI REFERENCE

#### 12.1 index

```
usage: pankmer index [-h] [-g <genome[s]{.fa,.fq,.tar,/}> [<genome[s]{.fa,.fq,.tar,/}> ..
↪.]]
                     [-p <genome[s]{.fa,.fq,/}> [<genome[s]{.fa,.fq,/}> ...]] -o
↪<output[.tar]> [--split-memory <int>]
                     [-t <int>] [--fraction <float> | -q <int>] [--gzip-level {1,2,3,4,5,
↪6,7,8,9}] [--time]

options:
  -h, --help            show this help message and exit
  -g <genome[s]{.fa,.fq,.tar,/}> [<genome[s]{.fa,.fq,.tar,/}> ...], --genomes <genome[s]
↪{.fa,.fq,.tar,/}> [<genome[s]{.fa,.fq,.tar,/}> ...]
                        paths to input genomes, directories, or tar archive
  -p <genome[s]{.fa,.fq,/}> [<genome[s]{.fa,.fq,/}> ...], --genomes-paired <genome[s]{.
↪fa,.fq,/}> [<genome[s]{.fa,.fq,/}> ...]
                        paths to input genomes or directories (paired)
  -o <output[.tar]>, --output <output[.tar]>
                        output directory or tarfile that will contain the k-mer index
  --rounds <int>        split indexing into multiple rounds to reduce memory usage
  -t <int>, --threads <int>
                        Number of threads to use [1]
  --fraction <float>    Fraction of k-mers to use. By default all k-mers are kept.
  -q <int>, --qual <int>
                        Completeness of index, in terms of bases, expressed as a Phred␣
↪quality score. e.g. --qual 30
                        means approx 1 in 1000 bases will be missed, which is equivalent␣
↪to --fraction 0.2. By default
                        no bases are missed.
  --gzip-level {1,2,3,4,5,6,7,8,9}
                        gzip compression level [6]
  --time                Report the time required to execute
```

#### 12.2 count

```
usage: pankmer count [-h] -i <index[.tar]> [<index[.tar]> ...]

options:
  -h, --help            show this help message and exit
  -i <index[.tar]> [<index[.tar]> ...], --index <index[.tar]> [<index[.tar]> ...]
                        a k-mer index
```

#### 12.3 collect

```
usage: pankmer collect [-h] -i <index[.tar]> [-o <output.{pdf,png,svg}>] [-t <output-
↪table.tsv>] [--title <"Plot title">]
                       [--width <float>] [--height <float>] [--color-palette <#color> [<
↪#color> ...]] [--alpha <float>]
                       [--linewidth <int>] [--conf | --contours <int> [<int> ...]]
                       [--legend-loc {best,upper left,upper right,lower left,lower right,
↪outside}]

options:
  -h, --help            show this help message and exit
  -i <index[.tar]>, --index <index[.tar]>
                        a k-mer index
  -o <output.{pdf,png,svg}>, --output <output.{pdf,png,svg}>
                        destination file for plot
  -t <output-table.tsv>, --table <output-table.tsv>
                        output TSV file containing plotted data
  --title <"Plot title">
                        set the title for the plot
  --width <float>       set width of figure in inches [4]
  --height <float>      set height of figure in inches [3]
  --color-palette <#color> [<#color> ...]
                        color palette to use
  --alpha <float>       transparency value for lines [1.0]
  --linewidth <int>     line width for plot [3]
  --conf                calculate confidence intervals for collection curves
  --contours <int> [<int> ...]
                        set contours for collection curves (in percent)
  --legend-loc {best,upper left,upper right,lower left,lower right,outside}
                        location of legend [best]
```

#### 12.4 upset

```
usage: pankmer upset [-h] -i <index[.tar]> -o <output.{pdf,png,svg}> -g <genome> [
↪<genome> ...] [-v] [-x] [-t <table.tsv[.gz]>]
                     [--show-counts] [--min-subset-size <int>] [--max-subset-size <int>]␣
↪[--time]

options:
  -h, --help            show this help message and exit
  -i <index[.tar]>, --input <index[.tar]>
                        a k-mer index
  -o <output.{pdf,png,svg}>, --output <output.{pdf,png,svg}>
                        destination file for upset plot
  -g <genome> [<genome> ...], --genomes <genome> [<genome> ...]
                        list of genomes to include
  -v, --vertical        draw the plot vertically
  -x, --exclusive       exclude k-mers that occur in genomes other than the input set
  -t <table.tsv[.gz]>, --table <table.tsv[.gz]>
                        write (optionally compressed) table of values in tsv format
  --show-counts         show counts for each subset
  --min-subset-size <int>
                        show only subsets larger than a minimum size
  --max-subset-size <int>
                        show only subsets smaller than a maximum size
  --time                report the time required to execute
```

#### 12.5 subset

```
usage: pankmer subset [-h] -i <input-index[.tar]> -o <output-index[.tar]> -g <genome> [
↪<genome> ...] [-x]
                      [--gzip-level {1,2,3,4,5,6,7,8,9}] [--time]

options:
  -h, --help            show this help message and exit
  -i <input-index[.tar]>, --input <input-index[.tar]>
                        a k-mer index
  -o <output-index[.tar]>, --output <output-index[.tar]>
                        destination file for subset index
  -g <genome> [<genome> ...], --genomes <genome> [<genome> ...]
                        list of genomes to include
  -x, --exclusive       exclude k-mers that occur in genomes other than the input set
  --gzip-level {1,2,3,4,5,6,7,8,9}
                        gzip compression level [6]
  --time                report the time required to execute
```

#### 12.6 adj-matrix

```
usage: pankmer adj-matrix [-h] -i <index[.tar]> -o <adjmatrix.{csv,tsv}> [--time]

options:
  -h, --help            show this help message and exit
  -i <index[.tar]>, --input <index[.tar]>
                        a k-mer index
  -o <adjmatrix.{csv,tsv}>, --output <adjmatrix.{csv,tsv}>
                        destination file for adjacency matrix
  --time                Report the time required to execute
```

#### 12.7 tree

```
usage: pankmer tree [-h] -i <adjmatrix.{csv,tsv}> [-n] [--metric {intersection,jaccard,
↪overlap,qv,qv_symmetric,ani}]
                    [--method {single,complete,average,weighted,centroid}] [--
↪transformed-matrix <matrix.{csv,tsv}>]

options:
  -h, --help            show this help message and exit
  -i <adjmatrix.{csv,tsv}>, --input <adjmatrix.{csv,tsv}>
                        adjacency matrix file
  -n, --newick          output tree in NEWICK format
  --metric {intersection,jaccard,overlap,qv,qv_symmetric,ani}
                        similarity metric [intersection]
  --method {single,complete,average,weighted,centroid}
                        clustering method [complete]
  --transformed-matrix <matrix.{csv,tsv}>
                        Write similarity transformed matrix to file
```

#### 12.8 clustermap

```
usage: pankmer clustermap [-h] -i <adjmatrix.{csv,tsv}> -o <adjmatrix.{pdf,png,svg}>
                          [--metric {intersection,jaccard,overlap,qv,qv_symmetric,ani}]
                          [--method {single,complete,average,weighted,centroid}] [--
↪colormap <color_map>] [--width <float>]
                          [--height <float>] [--heatmap-ticks {left,right}] [--cbar-
↪ticks {left,right}] [--dend-ratio <float>]
                          [--dend-spacer <float>]

options:
  -h, --help            show this help message and exit
  -i <adjmatrix.{csv,tsv}>, --input <adjmatrix.{csv,tsv}>
                        adjacency matrix file
  -o <adjmatrix.{pdf,png,svg}>, --output <adjmatrix.{pdf,png,svg}>
                        destination file for plot
  --metric {intersection,jaccard,overlap,qv,qv_symmetric,ani}
```

(continues on next page)

(continued from previous page)

```
                        similarity metric [intersection]
  --method {single,complete,average,weighted,centroid}
                        clustering method [complete]
  --colormap <color_map>
                        seaborn colormap for plot [mako_r]
  --width <float>       width of plot in inches [7]
  --height <float>      height of plot in inches [7]
  --heatmap-ticks {left,right}
                        Position of heatmap ticks. Must be "left" or "right" [left]
  --cbar-ticks {left,right}
                        Position of color bar ticks. Must be "left" or "right" [left]
  --dend-ratio <float>  Fraction of plot width used for dendrogram [0.2]
  --dend-spacer <float>
                        Fraction of plot width used as spacer between dendrogram and␣
↪heatmap [0.1]
```

#### 12.9 similarity

```
usage: pankmer similarity [-h] -i <adjmatrix.{csv,tsv}> -o <simmatrix.{csv,tsv}>
                          [--metric {jaccard,overlap,qv,qv_symmetric,ani}]

options:
  -h, --help            show this help message and exit
  -i <adjmatrix.{csv,tsv}>, --input <adjmatrix.{csv,tsv}>
                        adjacency matrix file
  -o <simmatrix.{csv,tsv}>, --output <simmatrix.{csv,tsv}>
                        similarity matrix file
  --metric {jaccard,overlap,qv,qv_symmetric,ani}
                        similarity metric [jaccard]
```

#### 12.10 distance

```
usage: pankmer distance [-h] -i <adjmatrix.{csv,tsv}> -o <distmatrix.{csv,tsv}> [--
↪metric {jaccard,overlap,ani}]

options:
  -h, --help            show this help message and exit
  -i <adjmatrix.{csv,tsv}>, --input <adjmatrix.{csv,tsv}>
                        adjacency matrix file
  -o <distmatrix.{csv,tsv}>, --output <distmatrix.{csv,tsv}>
                        distance matrix file
  --metric {jaccard,overlap,ani}
                        distance metric [jaccard]
```

#### 12.11 anchor-region

```
usage: pankmer anchor-region [-h] -i <index[.tar]> [<index[.tar]> ...] -r <reference.fa.
↪gz> -c <chr:start-end>
                             [-o <output.bdg[.gz]>] [-b] [-g <genes.gff3[.gz]>] [-f <int>
↪] [-p <int>]

options:
  -h, --help            show this help message and exit
  -i <index[.tar]> [<index[.tar]> ...], --index <index[.tar]> [<index[.tar]> ...]
                        index
  -r <reference.fa.gz>, --reference <reference.fa.gz>
                        reference
  -c <chr:start-end>, --coords <chr:start-end>
                        genomic coordinates
  -o <output.bdg[.gz]>, --output <output.bdg[.gz]>
                        write to file instead of standard output
  -b, --bgzip           block compress the output file
  -g <genes.gff3[.gz]>, --genes <genes.gff3[.gz]>
                        gff file of gene coordinates
  -f <int>, --flank <int>
                        size of flanking regions
  -p <int>, --processes <int>
                        number of processes to use
```

#### 12.12 anchor-genome

```
usage: pankmer anchor-genome [-h] -i <index[.tar]> [<index[.tar]> ...] -o <output.{pdf,
↪png,svg}> -r <reference.fa.gz> -c <chrX>
                             [<chrX> ...] [-t <output-table.tsv>] [--groups <"Group"> [<
↪"Group"> ...]] [--title <"Plot title">]
                             [--x-label <"Label">] [--legend] [--legend-title <"Title">]
                             [--legend-loc {best,upper left,upper right,lower left,lower␣
↪right,outside}] [--bin-size <int>]
                             [--width <float>] [--height <float>] [--color-palette <
↪#color> [<#color> ...]] [--alpha <float>]
                             [--linewidth <int>] [--processes <int>]

options:
  -h, --help            show this help message and exit
  -i <index[.tar]> [<index[.tar]> ...], --index <index[.tar]> [<index[.tar]> ...]
                        index
  -o <output.{pdf,png,svg}>, --output <output.{pdf,png,svg}>
                        destination file for plot
  -r <reference.fa.gz>, --reference <reference.fa.gz>
                        reference in BGZIP compressed FASTA format
  -c <chrX> [<chrX> ...], --chromosomes <chrX> [<chrX> ...]
                        chromosomes to include
  -t <output-table.tsv>, --table <output-table.tsv>
                        output TSV file containing plotted data
```

(continues on next page)

(continued from previous page)

```
  --groups <"Group"> [<"Group"> ...]
                        list of groups for provided indexes [0]
  --title <"Plot title">
                        set the title for the plot
  --x-label <"Label">   set the x-axis label for the plot
  --legend              include a legend with the plot
  --legend-title <"Title">
                        title of legend
  --legend-loc {best,upper left,upper right,lower left,lower right,outside}
                        location of legend [best]
  --bin-size <int>      Set bin size. The input <int> is converted to the bin size by␣
↪the formula: 10^(<int>+6) bp. The default
                        value is 0, i.e. 1-megabase bins. [0]
  --width <float>       set width of figure in inches [7]
  --height <float>      set height of figure in inches [3]
  --color-palette <#color> [<#color> ...]
                        color palette to use
  --alpha <float>       transparency value for lines [0.5]
  --linewidth <int>     line width for plot [3]
  --processes <int>     number of processes to use
```

#### 12.13 anchor-plot

```
usage: pankmer anchor-plot [-h] -t <genomecov.tsv> -o <output.{pdf,png,svg}> [--groups <
↪"Group"> [<"Group"> ...]]
                           [--loci <"chr:pos:name"> [<"chr:pos:name"> ...]] [--
↪chromsizes <file.chrom.sizes>]
                           [--title <"Plot title">] [--x-label <"Label">] [--legend] [--
↪legend-title <"Title">]
                           [--legend-loc {best,upper left,upper right,lower left,lower␣
↪right,outside}] [--width <float>]
                           [--height <float>] [--color-palette <#color> [<#color> ...]]␣
↪[--alpha <float>] [--linewidth <int>]

options:
  -h, --help            show this help message and exit
  -t <genomecov.tsv>, --table <genomecov.tsv>
                        genomic coverage results
  -o <output.{pdf,png,svg}>, --output <output.{pdf,png,svg}>
                        destination file for plot
  --groups <"Group"> [<"Group"> ...]
                        list of groups for provided indexes
  --loci <"chr:pos:name"> [<"chr:pos:name"> ...]
                        list of loci to mark on plot
  --chromsizes <file.chrom.sizes>
                        chromsizes file of the reference used to generate the table
  --title <"Plot title">
                        set the title for the plot
  --x-label <"Label">   set x-axis label for the plot
  --legend              include a legend with the plot
```

(continues on next page)

(continued from previous page)

```
  --legend-title <"Title">
                        title of legend
  --legend-loc {best,upper left,upper right,lower left,lower right,outside}
                        location of legend [best]
  --width <float>       set width of figure in inches [7]
  --height <float>      set height of figure in inches [3]
  --color-palette <#color> [<#color> ...]
                        color palette to use
  --alpha <float>       transparency value for lines [0.5]
  --linewidth <int>     line width for plot [3]
```

#### 12.14 dryrun

```
usage: pankmer dryrun [-h] [-g <genome[s]{.fa,.fq,.tar,/}> [<genome[s]{.fa,.fq,.tar,/}> .
↪..]]
                      [-p <genome[s]{.fa,.fq,.tar,/}> [<genome[s]{.fa,.fq,.tar,/}> ...]]␣
↪[--split-memory <int>] [-t <int>]

options:
  -h, --help            show this help message and exit
  -g <genome[s]{.fa,.fq,.tar,/}> [<genome[s]{.fa,.fq,.tar,/}> ...], --genomes <genome[s]
↪{.fa,.fq,.tar,/}> [<genome[s]{.fa,.fq,.tar,/}> ...]
                        input genomes
  -p <genome[s]{.fa,.fq,/}> [<genome[s]{.fa,.fq,/}> ...], --genomes-paired <genome[s]{.
↪fa,.fq,/}> [<genome[s]{.fa,.fq,/}> ...]
                        input genomes (paired)
  --rounds <int>        Parallel.This splits the indexing into multiple rounds; reducing␣
↪memory.
  -t <int>, --threads <int>
                        Number of threads to use
```

#### 12.15 download-example

```
usage: pankmer download-example [-h] [-d <dir/>] [-c {Spolyrhiza,Solanum,Zea,Hsapiens,
↪Bsubtilis,Athaliana}] [-n <int>]

options:
  -h, --help            show this help message and exit
  -d <dir/>, --dir <dir/>
                        destination directory for example data
  -c {Spolyrhiza,Solanum,Zea,Hsapiens,Bsubtilis,Athaliana}, --clade {Spolyrhiza,Solanum,
↪Zea,Hsapiens,Bsubtilis,Athaliana}
                        download publicly available genomes. Clade: max_samples.␣
↪Spolyrhiza: 3, Solanum: 46, Zea: 54, Hsapiens:
                        94, Bsubtilis: 164, Athaliana: 1135
  -n <int>, --n-samples <int>
                        number of samples to download, must be less than clade max [1]
```

### CHAPTER THIRTEEN

### API REFERENCE

**class** pankmer.**PKResults**(*results_dir: str*, *threads: int = 1*)

**decode_kmer**(*bits: bytes*) → int

Convert Kmer bytes to an integer (C unsigned long long)

**Parameters**
**bits** (*bytes*) – Bytes representing Kmer generated by PanKmer indexing

**Returns**
Python integer represting the Kmer represented by bytes

**Return type**
int

**decode_score**(*bits: bytes*) → list

Convert score bytes to a list of binary flags

**Parameters**
**bits** (*bytes*) – Score bytes generated by PanKmer indexing

**Returns**
List of binary flags indicating if genome at position x has the corresponding Kmer 0: Genome doesn't have Kmer 1: Genome has Kmer

**Return type**
list

**encode_kmer**(*kmer: str*) → int

Generate a integer represting canonical Kmer

**Parameters**
**kmer** (*str*) – Sequence representing one Kmer only

**Returns**
Python integer representing canonical Kmer

**Return type**
int

**get_blevel_scores**(*scores: list*) → list

Get base level score for each position in the reference

**get_collapsed_regional_scores**(*reference: str*, *regions: dict*) → dict

Retrieve regions from the reference and return their per-position Kmer scores

**Parameters**

- **reference** (*int*) – Path to a reference file in GZIP fasta format.

- **regions** (*int*) – Dictionary of regions. Key = name of contig, values = list of start and end positions to extract. If no start, end positions are given, the function will use the entire contig. example = {'contig_1': [[4, 10], [59, 90]], 'contig_2': []}

**Returns**
_description_

**Return type**
dict

`get_fasta_kmerbits`(*fasta_path: str*, *upper: int | None = None*, *lower: int | None = None*) → set

Get integers representing all canonical Kmers in a fasta file

**Parameters**

- **fasta_path** (*str*) – Path to fasta file
- **upper** (*int, optional*) – Python integer of the upper bound limit of canonical Kmer, by default None
- **lower** (*int, optional*) – Python integer of the lower bound limit of canonical Kmer, by default None

**Returns**
A set of integers representing all canonical Kmers in fasta file

**Return type**
set

`get_kmer_byte`() → bytes

read bytes in size of one kmer from kmers file

**Returns**
bytes representing Kmer

**Return type**
bytes

`get_kmer_scores`(*kmers: dict | None = None*) → dict

Retrieve scores of Kmer in the Kmers dictionary. if no dictionary is supplied then retrieve all scores.

**Parameters**
**kmers** (*dict, optional*) – Dictionary where the keys are target Kmers and values are default values to populate with if the Kmer isn't found in the index, by default None

**Returns**
Populated dictionary where keys are Kmers and values are scores

**Return type**
dict

`get_positional_fasta_kmerbits`(*fasta_path: str*, *contig_header: str = 'id'*) → dict

Get lists of integers representing each canonical Kmer in each position in each contig in the fasta file

**Parameters**

- **fasta_path** (*str*) – Path to fasta file
- **contig_header** (*str, optional*) – Header style to retrieve for contigs (id or description), by default 'id'

**Returns**
A dictionary where keys are contigs' names and values are lists of integers representing each canonical Kmer in each position in the corresponding contig

**Return type**
dict

`get_regional_coverage`(*reference: str*, *regions: dict*) → dict

Get per position coverage for each region in the reference specified by the regions dictionary.

**Parameters**

- **reference** (*str, required*) – A path to the reference file in fasta format
- **regions** (*dict, required*) – A dictionary whose keys are contigs in the refernce and the values are lists of start and end positons

**Returns**

- *A dictionary whose keys are contig names and start and end positions and*
- *values are lists of number of samples in the index that have that position*

`get_regional_scores`(*reference: str*, *regions: dict*, *cpu: int | None = None*) → dict

Retrieve regions from the reference and return their per-position Kmer scores

**Parameters**

- **reference** (*str*) – Path to a reference file in GZIP fasta format.
- **regions** (*dict*) – Dictionary of regions. Key = name of contig, values = list of start and end positions to extract. If no start, end positions are given, the function will use the entire contig. example = {'contig_1': [[4, 10], [59, 90]], 'contig_2': []}
- **cpu** (*int, optional*) – Number of cores to use. This will replicate memory used by parent function., default to PKResults threads.

**Returns**
A nested dictionary of scores. Key = contig, value = dictionary where key = tuple of start and end positons and value = list of per-position scores. example: {'contig_1': {(4, 10): [scores], (59, 90): [scores]}}

**Return type**
dict

`get_score_byte`() → bytes

read bytes in size of one score from scores file

**Returns**
bytes represting what samples have a Kmer

**Return type**
bytes

`get_sequence_kmerbits`(*seq: str*, *upper: int | None = None*, *lower: int | None = None*) → list

Get canonical Kmers making up the sequence

**Parameters**
**seq** (*str*) – Sequence to Kmerize

**Returns**
list of Python integers representing the canonical Kmers

**Return type**
list

`get_sequence_score`(*seq: str*, *kmers: dict | None = None*) → list

Retrieve scores of canonical Kmers in the sequence

**Parameters**

- **seq** (*str*) – Target sequence
- **kmers** (*dict, optional*) – dictionary of Kmer:Score to use to assign scores to Kmers in sequence. If no dictionary is supplied then the index will be used, by default None

**Returns**
List of integers representing score of Kmer ending at postion x in the sequence

**Return type**
list

`get_sequences_scores`(*sequences: dict*, *cpu: int | None = None*) → dict

Get per-position scores for a dictionary of sequences

**Parameters**

- **sequences** (*dict*) – A dictionary of sequences. Key = name, value = sequence
- **cpu** (*int, optional*) – Number of cores to use. This will replicate memory used by parent function.

**Returns**
A dictionary of scores. Key = name, value = list of per-position scores

**Return type**
dict

`reset_iter`()

Move Kmer and Score files' pointers to 0 position

`seek_kmer`(*kmer_num: int*)

Move Kmer file pointer past a number of Kmers

**Parameters**
**kmer_num** (*int*) – Number of Kmers to skip

`seek_score`(*score_num: int*)

Move Score file pointer past a number of scores

**Parameters**
**score_num** (*int*) – Number of scores to skip

`set_threads`(*threads: int*)

Set number of threads to run multithreaded processes

**Parameters**
**threads** (*int*) – Number of threads

`pankmer.index_wrapper`(*outdir: str*, *genomes_input=None*, *genomes_input_paired=None*, *split_memory: int = 1*, *threads: int = 1*, *index: str = ''*, *fraction: float = 1.0*, *gzip_level: int = 6*, *k=31*)

`pankmer.get_adjacency_matrix`(*results*)

Generate an adjacency matrix from a PKResults object

**Parameters**
**results** (PKResults) – a PKResults object

**Returns**
an adjacency matrix

**Return type**
DataFrame

pankmer.**clustermap**(*adj_matrix*, *output: str*, *cmap: str = 'mako_r'*, *width: float = 7.0*, *height: float = 7.0*, *metric='intersection'*, *method='complete'*, *optimal_ordering: bool = True*, *square: bool = True*, *heatmap_tick_pos='right'*, *cbar_tick_pos='left'*, *dendrogram_ratio: float = 0.2*, *dendrogram_spacer: float = 0.1*)

Plot a clustered heatmap of genome similarity values

**Parameters**

- **adj_matrix** (*DataFrame*) – a Pandas data frame representing the adjacency matrix
- **output** (*str*) – path to destination file for plot
- **cmap** (*str*) – color map for the heatmap
- **width** (*float*) – width of the plot in inches
- **height** (*float*) – height of the plot in inches
- **metric** (*str*) – similarity metric to use for clustering. must be “intersection”, “jaccard”, “overlap”, “qv” or “qv_symmetric”
- **method** (*str*) – linkage algorithm for hierarchical clustering. See *scipy.cluster.hierarchy.linkage* for details.
- **optimal_ordering** (*bool*) – If True, the linkage matrix will be reordered so that the distance between successive leaves is minimal. See *scipy.cluster.hierarchy.linkage*
- **square** (*bool*) – If True, the heatmap will be drawn square. This may force the figure dimensions to be altered slightly
- **cbar_tick_pos** (*str*) – Position of color bar ticks. Must be “left” or “right” [left]
- **dendrogram_ratio** (*float*) – Fraction of plot width used for dendrogram [0.2]
- **dendrogram_spacer** (*float*) – Fraction of plot width used as spacer between dendrogram and heatmap [0.1]

pankmer.**anchor_region**(**pk_results*, *ref*, *coords*, *summary_func=<function mean>*, *output_file=None*, *bgzip: bool = False*, *genes=None*, *flank: int = 0*, *processes: int = 1*)

Generate k-mer conservation levels across the input region

**Parameters**

- **pk_results** – a PKResults object
- **ref** (*str*) – path to a reference genome in BGZIP compressed FASTA format
- **coords** (*str*) – genomic coordinates formatted as ctg:start-end
- **summary_func** – function for summarizing the conservation of a kmer across genomes in the index. The default is statistics.mean
- **output_file** – filename or file object to write to
- **bgzip** (*bool*) – if True, output_file will be block compressed
- **genes** (*str*) – path to gff3 file containing gnes
- **flank** (*int*) – size of flanking region

- **processes** (*int*) – number of processes to use

**Yields**

*contig, start, end, *values* – row of bedGraph data

pankmer.**anchor_genome**(**pk_results, output: str, ref: str, chromosomes: list, output_table=None, summary_func=<function mean>, groups=None, loci=None, title: str = 'Genome anchoring', x_label: str = 'Chromosome', legend: bool = False, legend_title='Group', legend_loc='best', bin_size: int = 0, width: float = 7.0, height: float = 3.0, color_palette=['#1f77b4', '#ff7f0e', '#2ca02c', '#d62728', '#9467bd', '#8c564b', '#e377c2', '#7f7f7f', '#bcbd22', '#17becf'], alpha: float = 0.5, linewidth: int = 3, processes: int = 1, xtlabel_rotation: int = 0, xtlabel_ha: str = 'center'*)

Generate a plot of average k-mer conservation values in bins across the input genome

**Parameters**

- **pk_results** – a PKResults object
- **output** (*str*) – path to destination file for the plot
- **ref** (*str*) – path to the input reference genome in BGZIP compressed FASTA format
- **chromosomes** (*list*) – list of chromosomes to include in the plot
- **output_table** – path to write TSV table containing underlying plotting data
- **summary_func** – function for summarizing the conservation of a kmer across genomes in the index. The default is statistics.mean
- **groups** – iterable of group names
- **title** (*str*) – title of the plot
- **x_label** (*str*) – x-axis label
- **legend** (*bool*) – if true, draw a legend for the plot
- **legend_title** (*str*) – title for the plot legend
- **legend_loc** (*str*) – location of legend. must be one of “best”, “upper left”, “upper right”, “lower left”, “lower right”, “outside”
- **bin_size** – set bin size. The input <int> is converted to the bin size by the formula: 10^(<int>+6) bp. The default value is 0, i.e. 1-megabase bins.
- **width** (*float*) – width of the plot in inches
- **height** (*float*) – height of the plot in inches
- **color_palette** – color palette for plot lines
- **alpha** (*float*) – alpha value for plot lines
- **linewidth** (*float*) – width value for plot lines
- **processes** (*int*) – number of processes to use
- **xtlabel_rotation** (*int*) – rotation for x-axis tick labels
- **xtlabel_ha** (*str*) – horizontal alignment for x-axis tick labels (left, center, right)

### INDEX

## A

anchor_genome() (*in module pankmer*), 62
anchor_region() (*in module pankmer*), 61

## C

clustermap() (*in module pankmer*), 61

## D

decode_kmer() (*pankmer.PKResults method*), 57
decode_score() (*pankmer.PKResults method*), 57

## E

encode_kmer() (*pankmer.PKResults method*), 57

## G

get_adjacency_matrix() (*in module pankmer*), 60
get_blevel_scores() (*pankmer.PKResults method*), 57
get_collapsed_regional_scores() (*pankmer.PKResults method*), 57
get_fasta_kmerbits() (*pankmer.PKResults method*), 58
get_kmer_byte() (*pankmer.PKResults method*), 58
get_kmer_scores() (*pankmer.PKResults method*), 58
get_positional_fasta_kmerbits() (*pankmer.PKResults method*), 58
get_regional_coverage() (*pankmer.PKResults method*), 59
get_regional_scores() (*pankmer.PKResults method*), 59
get_score_byte() (*pankmer.PKResults method*), 59
get_sequence_kmerbits() (*pankmer.PKResults method*), 59
get_sequence_score() (*pankmer.PKResults method*), 59
get_sequences_scores() (*pankmer.PKResults method*), 60

## I

index_wrapper() (*in module pankmer*), 60

## P

PKResults (*class in pankmer*), 57

## R

reset_iter() (*pankmer.PKResults method*), 60

## S

seek_kmer() (*pankmer.PKResults method*), 60
seek_score() (*pankmer.PKResults method*), 60
set_threads() (*pankmer.PKResults method*), 60
